# H-NS silences antiviral immunity through 3D chromatin compaction

**DOI:** 10.64898/2026.09.05.749240

**Authors:** Timofey Khvostikov, Polina Iarema, Alexey Gavrilov, Ilya Shamovsky, Vitaly Epshtein, Aleksandr Andriianov, Polina Baikuzina, Elena Shagimardanova, Dmitry Senko, Thomas K Wood, Konstantin Severinov, Evgeny Nudler, Artem Isaev

## Abstract

Bacterial antiviral immunity systems can be toxic to their hosts and must be tightly regulated. Immunity genes are often encoded by AT-rich mobile genetic elements subjected to silencing by nucleoid binding protein H-NS. Using RNA-seq, ChIP-seq, and chromosome 3D reconstruction with Micro-C, we find that H-NS binds and compacts bacterial immunity loci. H-NS deletion reveals anti-phage activity in model *Escherichia coli* strains generally considered phage-sensitive. Extending this approach to environmental isolates, we show that removal of H-NS silencing enhances defense, allowing bacteria to restrict phages with anti-defense proteins. We also discover Madara, a novel immunity system that complements the co-regulated BREX defense. Our results establish H-NS as a master regulator of bacterial immunity and highlight the importance of expression levels for anti-phage activity in native hosts.

## Introduction

Bacterial defense systems demonstrate enhanced mobility compared to core metabolic genes and are frequently associated with mobile genetic elements (MGEs) (*1*). Although beneficial under phage predation, these systems can impose fitness costs through autoimmunity (*2*). Consequently, defense genes exhibit rapid turnover in bacterial populations and require tight regulation (*3*, *4*). While some systems respond to quorum sensing (*5*) or are transcriptionally upregulated during phage infection (*6–8*), the general principles governing regulation of defense genes’ expression remain unknown. Environmental stresses also modulate defense systems’ expression, and the coordinated response of genes within defense islands hints at their integration into a broader regulatory network (*9*). Given the tight association between defense genes and MGEs, we hypothesized that H-NS - a xenogenic silencer that suppresses AT-rich, horizontally acquired DNA - also regulates defense islands.

H-NS, together with its homolog StpA, sits at the top of the hierarchy of nucleoid-associated proteins that compact the bacterial chromosome (*10*). It has a preference for AT-rich, negatively supercoiled regions, and after nucleation can extend on the chromosome, forming filaments that interfere with transcription (*11–13*). In addition, H-NS has a bridging activity that stitches DNA like a zipper, forming compact chromosomal hairpins (CHINs) and hairpin domains (CHIDs) that are inaccessible to RNA polymerase (*14–16*). The Type I-E CRISPR-Cas of *Escherichia coli* K12 is a classic example of an inactive defense system repressed by H-NS (*17*, *18*), and here we explored whether such repression represents a broader phenomenon.

Common laboratory strains BL21, BW25113, and MG1655 are generally considered phage sensitive, although MG1655 encodes an active Type I R-M system EcoKI. We demonstrate that deletion of H-NS activates, while double deletion of H-NS and StpA significantly enhances, the antiviral defense in these strains. Initially focusing on the Eco1 retron system encoded by a cryptic prophage in BL21, we demonstrate that H-NS binds to its promoter and operon body, causing compaction and formation of a CHID. Double deletion of H-NS and StpA activates transcription, enhances msDNA production, and boosts Eco1-specific phage defense. Extending these observations, we demonstrate activation of the Lit protease in BW25113 and MG1655 backgrounds and identify a novel Sie protein that blocks T5 infection. Finally, we leveraged this approach to identify novel immunity systems in environmental isolates from the ECOR collection, showcasing an example of co-regulated BREX and modification-dependent Madara protein that complement each other’s defense spectra. Together, our results establish H-NS as a master regulator of bacterial defense islands.

## Results

### H-NS and StpA control retron Eco1 expression in *E.coli* BL21 via chromatin compaction

Eco1, formerly known as Ec86, is one of the most-studied retron immunity systems (*19– 22*). It is composed of a NAD-depleting toxic effector (Ndt) and a reverse transcriptase (RT) that synthesizes multi-copy single-stranded DNA (msDNA) on a matrix of non-coding RNA (ncRNA). The Ndt is activated when phage infection disrupts the structure of an msDNA-ncRNA hybrid, for example, via Dcm methylation (*23*). The system protects against T5-like phages, yet this activity has always been studied with overexpression vectors (*21*). Eco1 was originally discovered in the common laboratory strain BL21, where it is encoded in a P2-like prophage remnant (*19*, *24*) (**Fig. 1A, S1A)**. We screened a collection of T5 phages for their infectivity on BL21-AI and its ΔEco1 derivative (*21*), and observed only mild Eco1-mediated plaquing reduction for the phage Gostya9, but not the previously reported activity against T5 (**Fig. 1B)**.

**Figure 1.**
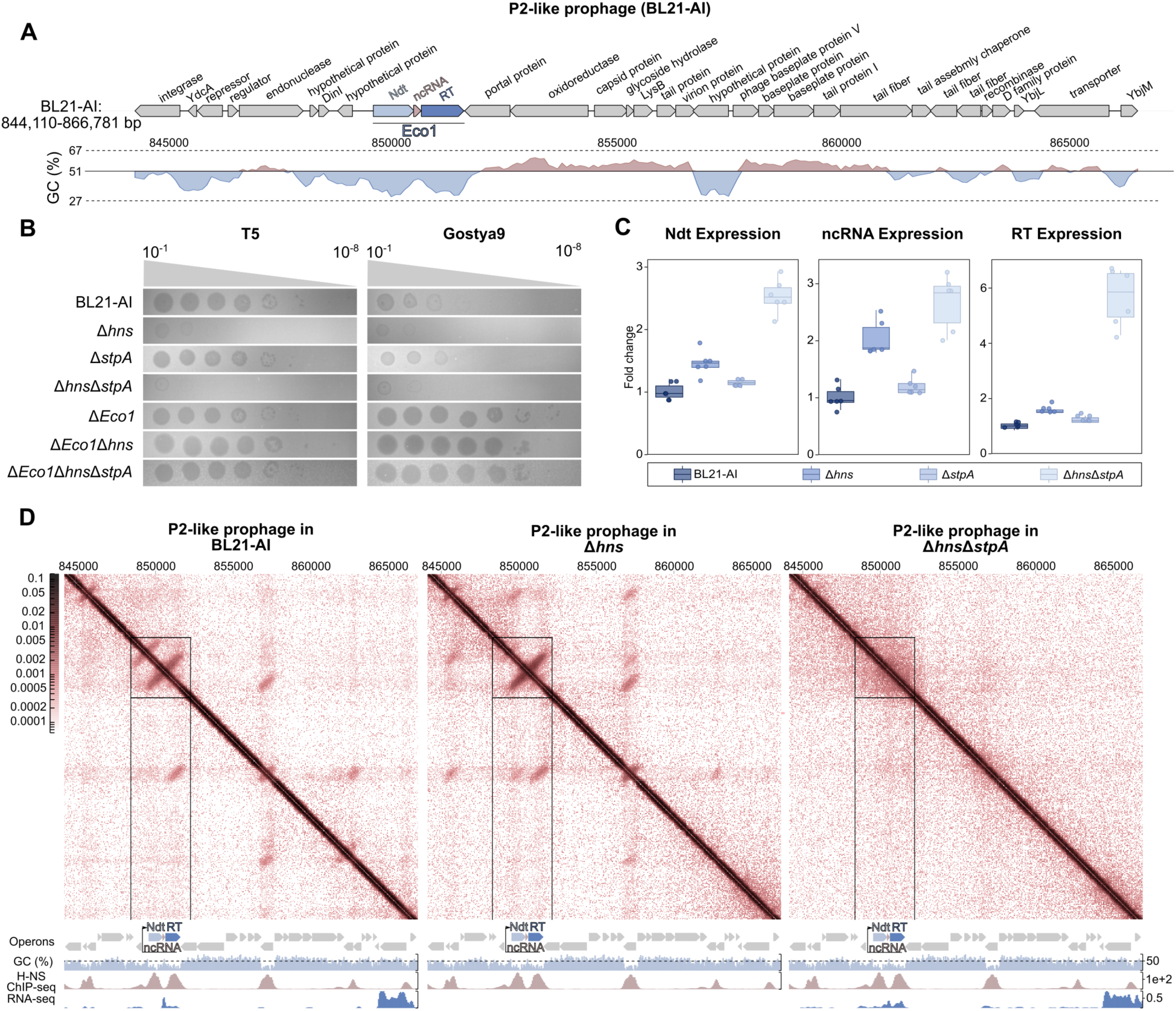
H-NS deletion activates Eco1 retron system in *E. coli* BL21-AI. **(A)** Genetic map and GC-content of the retron Eco1 locus in P2-like prophage of *E. coli* BL21-AI. **(B)** H-NS and the double H-NS/StpA deletions in BL21-AI enhance defense against T5 and Gostya9 phages in EOP assay (30°C)**. (C)** Expression levels of the Eco1 system components in BL21-AI and its derivatives, as determined by RT–qPCR. Signal is normalized to 16S rRNA expression level, with the expression level in WT cells set to 1 (30°C). **(D)** H-NS binding compacts Eco1 locus in BL21-AI and suppresses its transcription. MicroC maps, H-NS binding sites (CPM), and RNA-seq gene expression (log_2_(CPM+1)) of P2-like prophage locus in *E.coli* BL21-AI, *Δhns*, and *ΔhnsΔstpA* derivatives. ChIP-seq data obtained for the WT cells are shown under each panel.

We noticed that the GC content of the Eco1 region was significantly lower compared to the neighboring regions and hypothesized that H-NS repression could be responsible for the lack of anti-phage effect (**Fig. 1A**). H-NS, a major repressor of AT-rich regions, can form homodimers or heterodimers with its less abundant homolog, StpA; therefore, we tested the effects of individual and double H-NS/StpA deletions on the infectivity of phages T5 and Gostya9. In efficiency of plaquing (EOP) assays (**Fig. 1B, S1B**) and liquid culture infections (**fig. S1C**), a single *stpA* deletion had no effect, whereas deletion of *hns* significantly boosted anti-phage activity. This effect was even more pronounced in the double *hns*/*stpA* deletion, whereas none of the deletions affected phage infectivity in the *ΔEco1* strain, suggesting regulation of Eco1 expression by H-NS.

To directly test this possibility, we performed qPCR analysis of RT, ncRNA, and Ndt expression, confirming that deletion of *hns* or the double *hns*/*stpA* deletion upregulates Eco1 transcription (**Fig. 1C, S1D**). The deletions also result in enhanced production of msDNA (**Fig. S1E**). To further show that H-NS directly binds to the Eco1 locus, we performed H-NS ChIP-seq experiments. We identified three significant peaks of H-NS binding within the P2-like prophage, corresponding to the RT, Ndt, and DUF6216 genes, the regions with the lowest GC content (**Fig. 1D**). The same regions demonstrated transcription upregulation in ΔH-NS/ΔStpA culture as revealed via RNA-seq (**Fig. 1D**). Finally, we applied the Micro-C chromatin capture assay that provides 10-nucleotide resolution of 3D contacts in the *E. coli* chromosome (*16*). This analysis identified formation of two H-NS-bound hairpins (CHINs) enclosing the RT and Ndt genes (**Fig. 1D**). In addition, the Eco1 locus together with the upstream region was compacted within a larger chromosomal hairpin domain (CHID). *hns* deletion relaxed the Ndt CHIN and restructured the position of loops in the prophage region, whereas double *hns*/*stpA* deletion completely resolved short-scale inter-DNA contacts, opening the Eco1 locus for transcription. These results provide the first demonstration of immune system regulation via chromatin compaction. H-NS binding and DNA compaction events were noticed for other predicted prophage regions of the BL21 chromosome **(Extended data figure 1, Supplementary table 5)**.

We hypothesized that H-NS repression safeguards the bacterial cell from Eco1 effector autoimmunity, which may result from NAD^+^ depletion. Indeed, bacterial growth at 30°C was slower for the ΔH-NS/StpA strain compared to wild-type and ΔH-NS/StpA/Eco1 culture (**Fig. S1F,G**). *In vivo* NAD^+^ estimation via HPLC-MS demonstrated a notable and significant increase in NAD^+^ concentration in *ΔEco1* cells. This suggests that expression of NAD-depleting immunity systems can be costly, with an even higher cost in ΔH-NS cells (**Fig. S1H**).

### H-NS controls immunity genes in common *E.coli* laboratory strains BW25113 and MG1655

Activation of the Eco1 retron in the *Δhns* BL-21 strain suggested that other common laboratory strains could encode previously undetected immunity systems controlled by H-NS. Model strain BW25113 and a related strain MG1655 of the K-12 *E. coli* lineage are often used as phage-sensitive controls in studies of immunity systems and have been utilized for isolation of large phage collections (*25*). Lit protease and a variant of the Hachiman immunity system called AbpAB have been reported in BW25113, although their anti-viral activity was marginal (*26–28*), while MG1655 additionally encodes an active Type I R-M EcoKI. We compared infectivity of a laboratory phage collection against wild-type or *Δhns* BW25113 and MG1655 hosts and observed severe titer reductions for phages from the T4 and T5 groups in the *Δhns* background (**Fig. 2A**).

**Figure 2.**
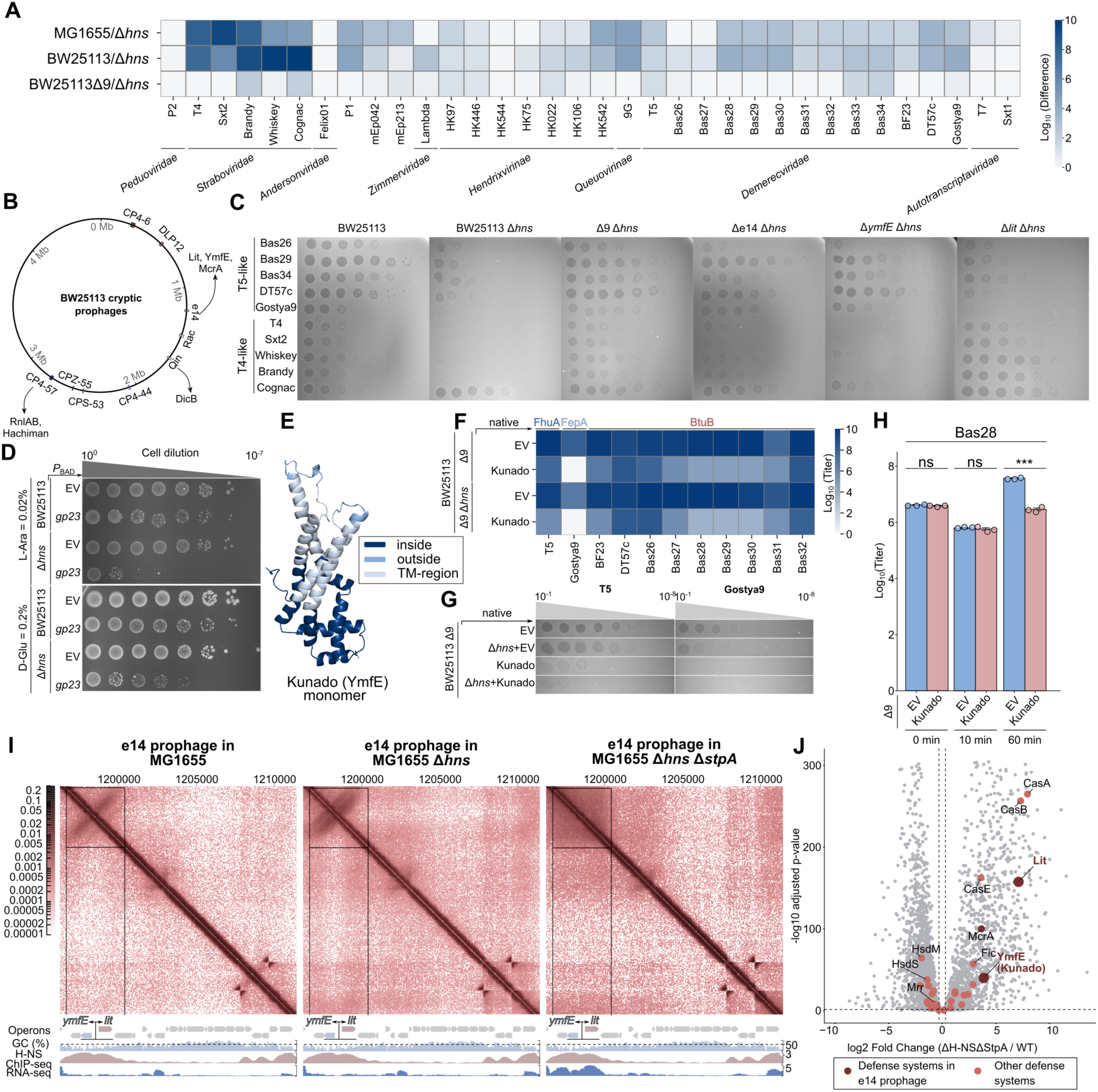
H-NS controls immunity genes in common *E.coli* laboratory strains BW25113 and MG1655. **(A)** Heatmap of the EOP assay demonstrating the difference in the phage titer between an indicated strain and its *Δhns* derivative. **(B)** Schematic illustration of the genomic positions of *E.coli* BW25113 cryptic prophages with a list of encoded defense systems. **(C**) EOP assay demonstrating that H-NS deletion in BW25113 provides defense against T5-like and T4-like phages, mediated by e14-prophage encoded *ymfE* and *lit* genes (30°C). **(D)** Serial dilution test demonstrates Lit-dependent toxicity of gp23 expressed from pBAD. **(E)** AlphaFold model of Kunado (YmfE) monomer with predicted membrane spanning regions. **(F)** Heatmap of the EOP assay demonstrating the defense activity of Kunado (YmfE) expressed from a plasmid under its native promoter. **(G)**. Representative plates from the EOP assay shown in **(F)**. **(H)** Efficiency of the phage Bas28 adsorption and release measured in the presence of Kunado (YmfE). **(I)** H-NS binding compacts defense locus in BL21-AI e14 prophage and suppresses its transcription. MicroC maps, H-NS binding sites (log_2_(IP/input)), and RNA-seq gene expression (log_2_(CPM+1)) of e14 prophage locus in *E.coli* MG1655, *Δhns*, and *ΔhnsΔstpA* derivatives (*16*). ChIP-seq data obtained for the WT cells are shown under each panel. **(J)** Volcano-plot demonstrating relative change of gene expression between MG1655 and the double *Δhns/ΔstpA* deletion strains.

Building on the assumption that H-NS preferentially suppresses horizontally acquired genetic elements, we used the BW25113 Δ9 strain, which lacks all cryptic prophages (*29*) (**Fig. 2B**). Phage infectivity on the *Δhns* lawns was restored in this background, hinting at prophage-encoded immunity systems activated upon H-NS deletion (**Fig. 2C**). We further tested BW25113 derivatives with deletions of individual prophages and localized the immune phenotype to the prophage e14 (**Fig. S2A**). Screening e14 gene deletions from the KEIO collection revealed two hits: deletion of the *lit* protease restored T4 phage infectivity in the *Δhns* strain, whereas restoration of T5-like phage infectivity required deletion of the previously uncharacterized gene *ymfE* (**Fig. 2C**).

Lit protease is activated by the phage T4 major capsid protein gp23, causing nonspecific protein degradation and cellular toxicity (*27*, *30*). We tested whether gp23 expression would cause toxicity in the *Δhns* background. Using cell plating, we confirmed that gp23 expression was not toxic in wild-type cells but caused Lit-dependent toxicity upon H-NS removal, confirming H-NS-mediated control of the Lit activity (**Fig. 2D, S2B**). The second identified protein, YmfE, has predicted transmembrane regions, leading us to hypothesize a role in blocking phage adsorption or injection (**Fig. 2E, S2C**). We cloned *ymfE* and independently confirmed its activity against T5-like phages in the BW25113 Δ9 strain (**Fig. 2F, G, S2D**). YmfE provided defense against phages that use distinct receptors and did not directly block adsorption (**Fig. 2F, H, S2E**), suggesting that it functions at later stages, possibly as a Sie protein that blocks DNA injection. Following the naming convention for novel immunity systems, we renamed this protein Kunado after the Japanese divine spirit that guards boundaries against the entry of malevolent forces.

Notably, *lit* and Kunado (*ymfE*) are neighboring genes encoded in opposite orientation, and their promoters can be co-regulated via compaction in the same hairpin. We confirmed their role in anti-phage defense in another common *E.coli* strain, MG1655, and utilized available RNA-seq, ChIP-seq, and Micro-C data to analyze the e14 prophage locus (**Fig. 2I, J, S2F and Extended data figure 2)**. Similar to the Eco1 locus in BL21, the Lit*/*Kunado locus of MG1655 contained a strong H-NS binding site, was highly compact, and formed a hairpin (CHIN). H-NS deletion restructured the locus, whereas the combined deletion of H-NS and StpA completely relaxed the hairpin and upregulated transcription of both genes. H-NS binding sites and DNA compaction were also noted for other prophages of MG1655 **(Extended data figure 2, Supplementary tables 6, 7)**. Together, the data demonstrate that H-NS/StpA play an important role in compactization and suppression of *E.coli* prophage cargo genes, including active immunity systems.

### Relaxation of H-NS repression allows identification of novel immunity systems in natural *E. coli* isolates

Having established that H-NS controls immunity systems in model laboratory strains of *E. coli*, we decided to test the universality of this phenomenon using natural *E. coli* isolates from the ECOR collection. Among these, ECOR28 and ECOR34 were amenable to P1 generalized transduction, enabling construction of *Δhns* derivatives. H-NS deletion reduced the plaquing efficiency of multiple phages in both ECOR strains (**Fig. 3A, 4A**). In contrast, majority of these phages were not sensitive to H-NS deletion in the BW25113 Δ9 strain (**Fig. 2A**), suggesting that upregulation of immunity systems in ECOR strains is a plausible cause of the titer reduction. To identify H-NS-repressed systems, we performed RNA-seq on wild-type and *Δhns* ECOR28 and ECOR34, quantifying expression of known and putative defense genes predicted by PADLOC and DefenseFinder. As expected, *hns* deletion resulted in upregulation of the type I-E CRISPR-Cas loci (*17*, *18*). Beyond this, we identified several immune systems upregulated upon H-NS deletion. These included PD-λ-4, PDC-S11, and *brxL* gene (a component of the BREX system) in the ECOR34 strain; and ShosTA, Lamassu, Kongming, McrBC, and PDC-S12 in the ECOR28 strain (**Fig. 3B, 4B, Supplementary tables 8, 9**).

**Figure 3.**
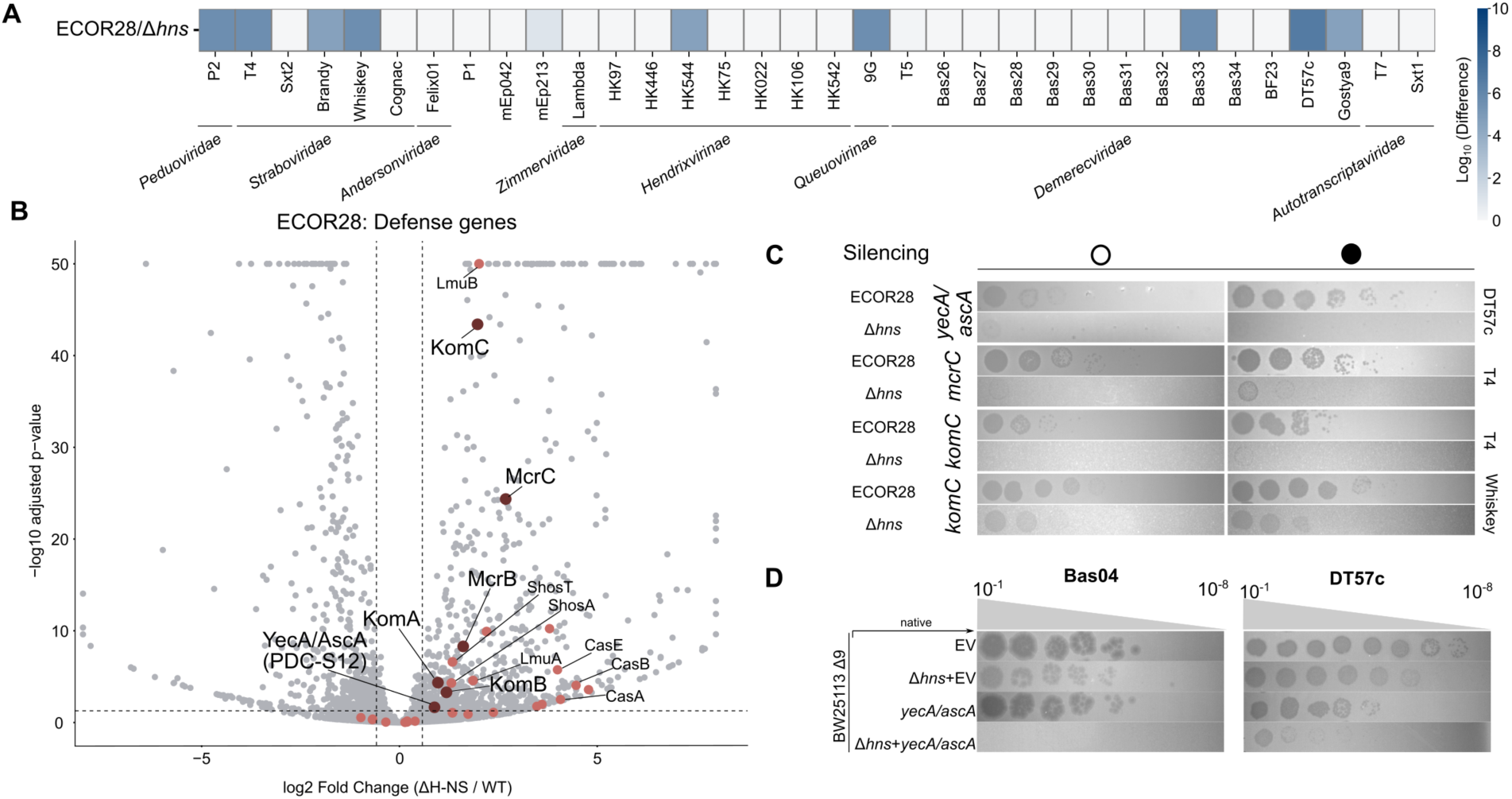
Relaxation of H-NS repression activates anti-phage defence in natural *E. coli* isolate ECOR28. **(A)** Heatmap of the EOP assay demonstrating the difference in the phage titer between ECOR28 and its *Δhns* derivative. **(B)** Volcano-plot demonstrating relative change of gene expression between ECOR28 its *Δhns* derivative. **(C)** EOP with the phages DT57c and T4 performed with ECOR28 and its *Δhns* derivative in conditions of dCas9 silencing of the *mcrC* (R-M IV) or PDC-S12 (*yecA/ascA*) genes. **(D)** EOP assay demonstrating the defense activity of PDC-S12 (*yecA/ascA*) expressed from a plasmid under its native promoter.

We obtained Micro-C maps of the ECOR28 wild type, *Δhns*, and *Δhns*/*ΔstpA* strains, revealing that differentially expressed immunity loci, including Lamassu, ShosTA, Kongming, and McrBC, are compacted, while most of the three-dimensional contacts were relaxed upon double deletion of *hns* and *stpA* (**Fig. S3A-E, Extended data figure 3**). Surprisingly, we also observed the loss of the Kongming-encoding prophage upon double deletion of H-NS and StpA, suggesting that chromatin compaction controls prophage induction, as recently reported for BW25113 (*31*) **(Fig. S3E)**. To assess the role of individual immunity systems in phage suppression in the *Δhns* background, we performed transcriptional knockdowns of upregulated genes using dCas9 silencing.

We were unable to restore phage infectivity by targeting the Lamassu or ShosTA systems. It remains possible that dCas9 silencing did not achieve sufficient repression of these systems or that alternative yet unknown genes contribute to the phage inhibition in the ECOR28 *Δhns* strain. However, we noted increase of T4 phage titer upon dCas9 targeting of the Type IV R-M system McrBC and Kongmin, while T5-like phage DT57c plaquing ability was restored upon targeting of the candidate PDC-S12 immunity system (**Fig. 3C and S4A**). While the classical *E.coli* McrBC system is inactive against glucosylated T4 DNA (*32*), recently reported McrBC homologs were capable to restrict wild-type T4 phage (*33*). Our results suggests that ECOR28 McrBC system, that is distinct from previously studied McrBC complexes, also have specificity towards glucosylated cytocines or targets non-glucosylated DNA during phage replication. As for the PDC-S12 system, dCas9 silencing was efficient only in the wild type cells. Therefore, to further verify H-NS regulation and the immunity function of this gene, we cloned it under native promoter and expressed in BW25113 Δ9 strain. This confirmed protective activity against multiple T5- and T1-like phages, that was enhanced upon *hns* deletion (**Fig. 3D and S4C, S3B**). Strikingly, PDC-S12 is identical to YecA (AscA), an accessory component of the Sec-dependent protein translocation pathway, carrying a Ferrum-binding domain (*34*, *35*) (**Fig. S4B**). BW25113 also encodes the PDC-S12 gene; however, its deletion did not alter plaquing ability for most of the T5- and T1-like phages, except for the phage Bas08 (**Fig. S4D**). Further studies are needed to elucidate whether PDC-S12/YecA/AscA acts as a *bona fide* immunity system or whether its expression affects phage receptors availability.

Next, we focused on immunity genes upregulated in the ECOR34 *Δhns* strain (**Fig. 4A, B**). Here, silencing of the BREX system gene *brxL* revealed an interesting pattern. Phage Sxt1, a close relative of T3 (*36*), was able to plaque on ECOR34, but not on the *Δhns* derivative (**Fig. 4C)**. Silencing of the *brxL* gene restored Sxt1 plaquing ability in the *Δhns* background, directly linking BREX anti-phage activity to H-NS repression. Sxt1 phage encodes an anti-BREX protein SAMase that depletes the S-adenosyl-methionine co-factor required for BREX activity (*37*). Consequently, a SAMase deletion mutant (Sxt1*Δ0.3*) was completely repressed by BREX in the ECOR34 strain, while *brxL* silencing permitted Sxt1*Δ0.3* growth (**Fig. 4C)**. Therefore, we uncover that the native expression level of the BREX system is sufficient to suppress Sxt1*Δ0.3* phage, while the wild type phage overcomes BREX defense due to the activity of SAMase. However, elevated BREX expression in the *Δhns* background helps to restrict the phage armed with an inhibitor. A similar effect of elevated defense required to overcome the phage with an anti-BREX protein was recently linked to the ssDNA-sensing WYL regulator (BrxR/CapW) associated with BREX and CBASS immunity loci (*8*, *38*, *39*). Our results confirm that expression of the immunity genes can be fine-tuned to cope with the anti-immune proteins.

**Figure 4.**
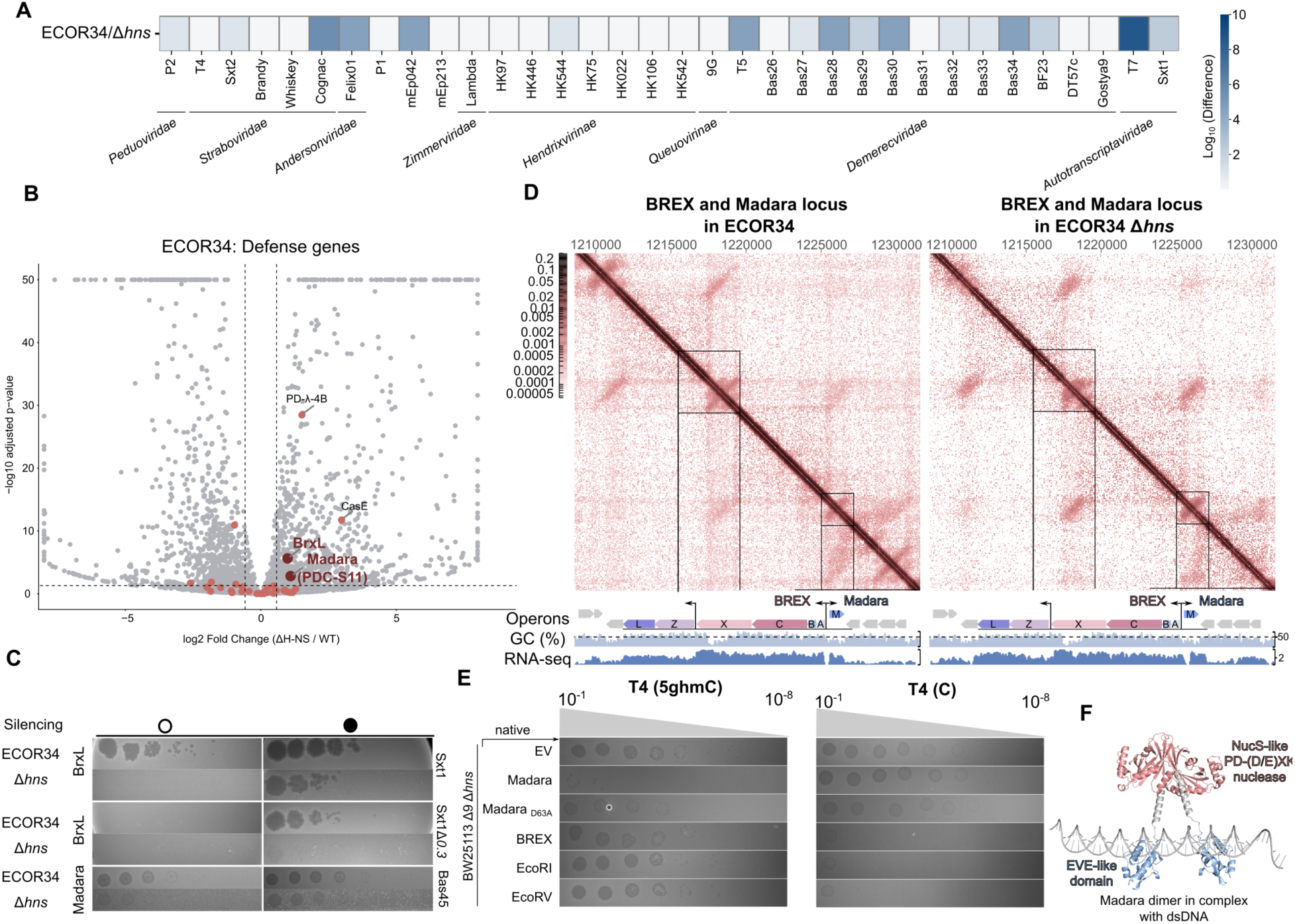
Relaxation of H-NS repression activates anti-phage defence in natural *E. coli* isolate ECOR34. **(A)**. Heatmap of the EOP assay demonstrating the difference in the phage titer between ECOR34 and its *Δhns* derivative. **(B)** Volcano-plot demonstrating relative change of gene expression between ECOR34 its *Δhns* derivative. **(C**) EOP with the phages Sxt1, Sxt1Δ0.3 and Bas45 performed with ECOR34 and its *Δhns* derivative in conditions of dCas9 silencing of the *brxL* (BREX) or PDC-S11 (Madara) immunity systems. **(D)** 3D chromosome compactization, GC content and gene expression levels (log_2_(CPM+1)) of the BREX/Madara defense locus in the ECOR34, and its *Δhns* derivative. **(E)** EOP assay with wt T4 (5ghmC) and modification-deficient T4 (C) performed on BW25113 Δ9*Δhns* in the presence of plasmid-expressed Madara, BREX or Type II R-M systems. **(F)** AF3 model and predicted domain organization of Madara (PDC S-11) dimer in complex with DNA.

We further compared Micro-C maps of ECOR34 and its *Δhns* derivative **(Fig. 4D and S5A, B, Extended data figure 4)**. Two operons of the BREX system (*brxABCX* and *brxZL*) were each compacted in a CHID, with the *brxA* promoter region and *brxX* gene region enclosed in a hairpin. H-NS deletion partially relaxed these three-dimensional contacts and upregulated transcription of the *brxZL* operon (**Fig. 4D**). Unfortunately, we were unable to generate a double *hns*/*stpA* deletion, which would be required for complete DNA decompaction, suggesting that derepression of the chromosome in this strain could be toxic. Notably, the gene body and promoter of the differentially expressed PDC-S11 system was enclosed in the hairpin with the *brxA* promoter, mirroring the scenario of the *lit/ymfE* locus. To test whether these systems might be co-regulated due to complementary defense profiles, we screened the entire BASEL phage collection against dCas9-silenced *brxL* or PDC-S11 **(Fig. 4C, S6A)**. Phages Bas39, Bas40, and Bas45 -all belonging to the *Tevenvirinae* subfamily, known for DNA hypermodification - were sensitive to PDC-S11, but not to the BREX system (**Fig. S6A**). We therefore rename this system Madara, after a demon-protector from Japanese mythology.

BREX system is inhibited by DNA glucosylation of T4-like phages (*40*), and could be coupled with a modification-specific Type IV R-M protein (BrxU), providing backup defense (*41*). To confirm that Madara provides similar complementary defense against T4-like phages, we cloned the system under its native promoter and compared the BASEL phages’ sensitivity profiles towards ECOR34 Madara and the BREX system from HS strain expressed in the BW25113 Δ9 strain. Madara restricted multiple BREX-resistant *Tevenvirinae* phages, an activity that was enhanced upon *hns* deletion (**Fig. S6B**). Consistently, Madara restricted wild type T4 (5ghmC) that was resistant to BREX and Type II R-M defense, but was inactive against the T4 (C) strain, devoid of cytosine modification, and therefore sensitive to BREX and Type II R-M systems (**Fig. 4E**).

Analysis of AF3-predicted Madara structure revealed an N-terminal PD-(D/E)XK nuclease, similar to that of the NucS DNA repair enzyme (**Fig. 4F**). D63A mutation of the nuclease catalytic motif abolished phage defense (**Fig. 4E**). The nuclease NucS-like domain was connected via a linker to a C-terminal domain distantly resembling an EVE fold. EVE domains are widespread across modification-sensing immunity systems (*42*, *43*), and we suppose that within Madara, the EVE-like domain could recognize modified T4 cytosines, triggering PD-(D/E)XK domains dimerization and further phage DNA cleavage or a non-specific abortive response. Together, our data reveal an example of a co-regulated BREX/Type IV R-M Madara defense island controlled by H-NS DNA compaction and providing a complementary defense against modified and non-modified phages.

## Discussion

By regulating horizontal gene transfer and mediating the selection of phage-resistant variants, anti-phage immunity systems profoundly influence bacterial evolution, yet the principles governing their regulation remain largely unexplored. While these systems are essential for survival, their constitutive expression can impose fitness costs (*2*). Here, we identify the nucleoid-associated protein H-NS as a master silencer of defense islands. We show that H-NS compacts the chromatin of defense loci, repressing the expression of immunity genes. H-NS is a well-known xenogenic silencer that targets horizontally acquired AT-rich DNA and defense genes frequently associate with mobile elements and prophages (*44*, *45*). Notably, while lytic genes are tightly controlled by phage-encoded repressors, immunity genes must remain phenotypically active to confer protection. We propose that prophages may therefore delegate the regulation of these genes to the bacterial host.

We discovered that common laboratory strains of *E. coli*, typically used as immunity-negative controls, harbor functional but silent immunity genes. These genes are robustly activated upon deletion of *hns* or double deletion of *hns* and *stpA*. H-NS-dependent repression was also observed in the natural isolates ECOR28 and ECOR34, suggesting it may represent a general mechanism for bacteria encoding H-NS or its functional homologs. Notably, the inhibition of plasmid conjugation was also recently linked to the putative H-NS regulated modules restricting incoming plasmid DNA (*46*). Whereas the activated systems in BL-21 (retron Eco1) and BW25113/MG1655 (Lit and Kunado) are prophage-encoded, many genes upregulated in the ECOR strains were located outside annotated prophage regions **(Extended data figures, Supplementary tables 5-9)**. While we cannot rule out their association with other mobile genetic elements, H-NS binding to immunity genes might represent an independent regulatory adaptation, distinct from its well-established role in silencing of the insertion hotspots of mobile elements (*47*). Analysis of all predicted immunity genes across complete bacterial genomes revealed a significant shift towards increased AT-content, suggesting that immunity genes could represent a preferred substrate for H-NS binding **(Fig. S7)**.

A critical question is whether H-NS repression is alleviated under specific conditions to bolster phage defense. Given that H-NS activity is stress-responsive (*48*, *49*) and growth conditions were shown to modulate coordinated defense island expression (*9*), H-NS may be responsible for previously observed global effects. Phages themselves encode H-NS inhibitors (*50–53*), hinting at a single-cell response model where infection relieves repression. Furthermore, phage-induced lysis releases metabolites that could be sensed by neighboring cells, potentially serving as a population-level signal (*54*, *55*). H-NS repression could be integrated into such population-level pathways (*56*), and recent evidence implicates small metabolites in modulating H-NS function (*57*). This regulatory paradigm mirrors the transcriptional activation of innate immune systems in higher eukaryotes, achieved at both the single-cell and population levels via interferon signaling (*58*).

Exploration of the H-NS-controlled genes revealed an interesting phenomenon: H-NS might co-regulate two inversely oriented operons by stabilizing a hairpin structure in their shared promoter region **(Fig. 5)**. This was evident for the Lit protease and Kunado (*ymfE*) and for the BREX and Madara (PDC-S11) loci. Madara is a novel modification-dependent immunity system that appears to cooperate with BREX, analogous to the previously reported BREX-BrxU partnership (*41*). In addition, Madara could be an example of a DNA repair protein repurposed for immunity, as its PD-(D/E)XK nuclease domain resembles that of an archaeal NucS (*59*). Although a NucS-like domain was recently identified in a family of Type IV restriction-modification nucleases (*60*), the predicted modification-sensing domain of Madara is distinct, and its mechanism warrants further investigation.

**Figure 5.**
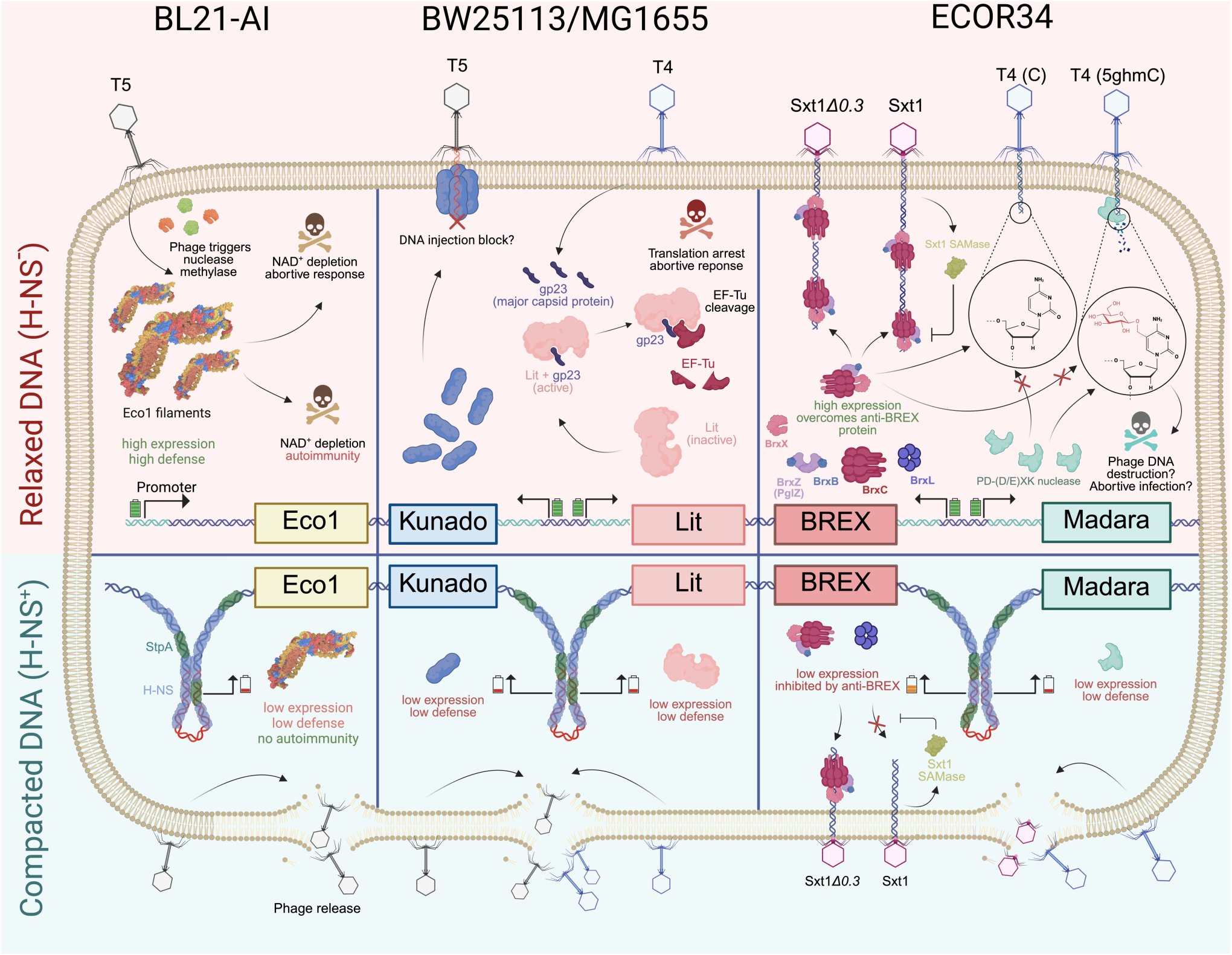
Proposed model for the multiple roles of H-NS in silencing of bacterial antiviral immunity systems.

Critically, removal of H-NS repression was sufficient to enhance bacterial defense and overcome a phage expressing an anti-defense protein. This observation aligns with the gene dosage model of phage-bacterial conflict (*61*, *62*), in which the expression levels of immune and anti-immune components are tightly balanced. However, elevated immunity gene expression is also linked to toxicity, making constitutive activation costly in the absence of phage pressure. We propose that dedicated regulators (e.g., BrxR/CapW) (*8*, *38*, *39*) and master silencers like H-NS provide a dynamic range of expression, enabling bacteria to finely balance immunity system activity to mitigate its protective benefits and fitness costs (**Fig. 5 and S7**).

## Materials and Methods

### Strains and Phages

The bacterial strains used in this study are listed in **Supplementary table 1**. Bacteria were cultured in LB medium (10 g/L Tryptone, 5 g/L Yeast Extract, 10 g/L NaCl, supplemented with appropriate antibiotics) under aerobic conditions (180 rpm/min) or on solid LB medium (supplemented with 1.5% agar). The phage strains, along with their sources, are provided in **Supplementary table 2**.

### Plasmids

The genes encoding Madara and Kunado were amplified via PCR from ECOR34 and BW25113 genomes and then cloned into the pBR322 expression vector using the Gibson assembly method. The gp23 gene was amplified via PCR from the T4 genome and then cloned into the pBAD expression vector using the Gibson assembly method. Plasmids for dCas9 silencing were constructed using the Golden Gate method in the pFD152 expression vector. All used plasmids and oligos are listed in **Supplementary table 3 and 4.**

### Strains construction

Strains with deletions were generated by transducing the corresponding alleles from the Keio collection. using phage P1_vir_, followed by selection for kanamycin resistance. To eliminate FRT-flanked Kmr cassette from the strain, the pCP20 plasmid was used. Ampicillin-resistant transformants were selected at 30 °C, then purified without selection at 37 °C to cure the helper plasmid. Successful removal of all antibiotic resistance markers was confirmed. All deletions were confirmed using PCR with specific gene-flanking primers.

### Construction of the Sxt1Δ0.3 mutant phage

To delete the *0.3* gene from the Sxt1 genome, we used a SpCas9-based counterselection system combined with homologous recombination, following previously described protocols (*63*). *E. coli* BW25113 cells carrying three plasmids, pSpCas9, pKDsgRNA expressing a guide RNA specific to the Sxt1 *0.3* gene, and a donor plasmid carrying 300 bp homology arms flanking the deletion region, were used as the host (*64*). An overnight culture of the triple-plasmid strain was mixed with molten 0.6% LB top agar and poured onto LB agar plates. Wild-type Sxt1 phages were spotted onto the solidified top agar, and plates were incubated overnight at 37 °C. Individual plaques were picked, resuspended in LB medium, and replated for a second round of single-plaque purification. From the second round, individual plaques were picked and screened by PCR using primers flanking the *0.3* locus. To confirm the deletion functionally, the mutant phages were tested in vivo against *E. coli* MG1655, which carries the EcoKI restriction-modification system. Phages lacking the *0.3* gene, which encodes the SAMase protein that protects against this system, were sensitive to EcoKI, whereas wild-type Sxt1 remained resistant. Confirmed clones were propagated in 5 mL of BW25113 culture, and lysates were treated with chloroform and stored at 4 °C.

### LC-MS NAD+ analysis

Cultures were grown under the following conditions: 30°C with shaking. Briefly, 15 mL of culture (OD_600_=1) was transferred to Eppendorf tubes and centrifuged (4000 g, 5 min, 4 °C). The supernatant was removed, and the cell pellets were stored at −80 °C. For the metabolite extraction, samples were homogenized in 1 mL of ice-cold methanol/water (80:20, v/v) containing 0.1% formic acid. After sonication (10 min, 4 °C), samples were shaken on an orbital shaker (10 min, 4 °C) and centrifuged (12700 g, 10 min, 4 °C). The supernatant was transferred to new tubes, and the extraction was repeated with an additional 0.5 mL of methanol/water (80:20, v/v). The combined supernatants were pooled, and the solvent was removed under reduced pressure. Prior to UPLC-MS/MS analysis, the samples were resuspended in 100 μL of acetonitrile/water (75:25, v/v). For targeted metabolomics analysis, 6 μL of the resuspended sample was injected into a Waters ACQUITY UPLC system (Waters). Separation was achieved on a SeQuant® ZIC®-HILIC column (2.1 × 100 mm, 3.5 μm) equipped with a guard column of the same type, maintained at 45 °C. Mobile phase A consisted of 10 mM ammonium formate in water containing 0.125% formic acid, and mobile phase B consisted of 10 mM ammonium formate in acetonitrile containing 0.125% formic acid. The flow rate was set to 0.4 mL/min. The gradient program was as follows: 0–1 min, 95% B; 1–10 min, linear gradient from 95% to 5% B; 10–12 min, 5% B; 12–12.5 min, linear return to 95% B; and 12.5–15 min, re-equilibration at 95% B. Mass spectrometric analysis was performed on a Thermo Fisher Q Exactive mass spectrometer equipped with a heated electrospray ionization (HESI) source (Thermo Fisher Scientific) using parallel reaction monitoring (PRM). The electrospray ionization conditions were as follows: spray voltage, 4.5 kV; sheath gas flow rate, 45 arbitrary units; auxiliary gas flow rate, 20 arbitrary units; sweep gas flow rate, 4 arbitrary units; capillary temperature, 250 °C; auxiliary gas heater temperature, 350 °C; and S-lens RF level, 70. Full MS scans were acquired at a resolution of 70,000, AGC target of 5e5, maximum injection time of 100 ms, and scan range of 50–700 m/z. PRM scans were acquired at a resolution of 17,500, AGC target of 2e5, maximum injection time of 100 ms, isolation window of 1.2 m/z, and stepped normalized collision energies of 15, 35, and 55. The target ions monitored were NAD+, m/z 664.116; ADPr, m/z 560.079; cADPr, m/z 542.068; and nicotinamide, m/z 123.055. Fragment ions used for quantification were m/z 136.061 for NAD+, 159.027 for ADPr, 134.019 for cADPr, and 80.050 for nicotinamide. Peak areas and peak heights were quantified using Thermo Xcalibur Quan Browser software (Thermo Fisher Scientific).

### Library construction and sequencing

The quality of total RNA was evaluated using the Bioanalyzer 2100 (Agilent, Santa Clara, CA, USA). 160 ng of RIN ≥7 of total RNA was used for rRNA depletion by means of the Ribo-Zero Plus rRNA Depletion kit (Illumina, San Diego, CA) following the manufacturer’s instructions. The depleted RNA was purified using RNAClean XP beads (Beckman Coulter, Brea, CA). Resulting RNA was then used for library construction using NEBNext® Ultra II™ Directional RNA Library Prep Kit for Illumina (New England Biolabs, Ipswich, MA, USA) according to the manufacturer’s instructions. The quality of libraries was verified using the Bioanalyzer 2100 (Agilent, Santa Clara, CA, USA) and the yield was validated by qPCR. Sequencing was performed on NovaSeq6000 (Illumina, San Diego, CA, USA) with pair-end 150 bp reading at the Genomics and Bioimaging Core Facility.

### Toxicity assay

Lit toxicity in the presence of gp23 (T4) or empty vector was measured using a spot-test assay. Native Lit^+^ (BW25113 or BW25113*Δhns*) and Lit^−^ (BW25113*Δlit* or BW25113*ΔlitΔhns*) cultures carrying the pBAD vector-encoding-indicated trigger were grown overnight in 10 ml of LB at 37 °C. Stationary cultures were diluted to OD_600_ of 0.6 and plated on LB agar plates supplemented with 0.02% L-Ara by serial tenfold dilution. Control plates without induction contained 0.02% D-Glu to prevent leakage of the araBAD promoter.

### H-NS Chip-seq analysis

For ChIP-seq experiment, overnight LB MG1665 (or BL21-AI) culture was diluted 1:100 with LB (total volume 600 mL) and grown to OD=0.6, followed by adding 16% formaldehyde to 0.25% for 20 minutes at +37C with shaking. Cells were collected by centrifugation 5 minutes at 5000g and re-suspended in 25mL Lysis buffer 3’ [10mM Tris pH8.0; 100mM NaCl; 1mM EDTA; 0.5mM EGTA] supplemented with human lysozyme 1mg/ml and protease inhibitor cocktail (Roche). Sample was sonicated in an ice-bath for 10 min with constant pulse at a midi-tip (power output 70%) in a metal beaker. The resulting lysate was spun for 30 minutes at 30 000 g and 20 uL of the supernatant was taken as an input control. The rest of the lysate was mixed with 2500 units of EndoCleava (Benzonase, CCNBio) and incubated for 1 hour on ice. 0.1% Sodium Deoxycholate; 0.5% N-lauroylsarcosine and 1% Triton X-100 were added to the sample before 20ug anti-H-NS antibodies (Cusabio) were added as well. The sample was rotated overnight at +4C in 50mL Falcone tubes.

Next day protein A/G magnetic beads (100ul - dry bed. Pierce) pre-equilibrated with 1ml of ice cold 1X PBS+0.5% BSA (blocking solution) were added to the sample and incubated with rotation for 1 hour at +4C. Beads were separated on magnetic stand and transferred into fresh 1.7 mL Eppendorf tube in 1ml of RIPA buffer [50 mM HEPES pH7.5; 250 mM LiCl; 1mM EDTA; 1% NP40; 0.7% Sodium Deoxycholate]. Beads were rotated for 10 minutes at +4C, separated on a magnetic stand, and the wash was repeated for a total of five times. Beads were washed once more in 1ml of TE buffer as above before reversal of crosslinked-sonicated-chromatin. Washed sample and input control were mixed with 250 uL of TE buffer and incubated overnight at 65°C shaking with 1% SDS and 2 mg/mL proteinase K [from 20 mg/mL stock solution in water, GoldBio].

Next day samples were briefly spun down, beads separated on a magnetic stand and DNA was purified by using Chip DNA Clean and concentrator kit (Zymo-research). DNA concentration was measured by Qubit.

### Micro-C

Micro-C was performed as previously described (*16*), with the exception of the DNA sonication step. Briefly, cells at OD₆₀₀ ∼0.6 were cross-linked with formaldehyde and DSG, then treated with lysozyme and MNase. DNA ends were biotin-labeled and religated. The DNA was then purified and sonicated using a Bioruptor Pico sonicator (five 30-second pulses). Following biotin pull-down, DNA ends were repaired, A-tailed, and ligated to Illumina adapters. Libraries were amplified with 9–10 PCR cycles, purified with AMPure XP beads, and sequenced (paired-end) on an Illumina NextSeq 2000 platform. Sequencing reads were processed as described (*16*).

### Visualization of the GC content

Analyses of the GC content showcased in Figures 1 and S1 was performed in R 4.4.0 with Biostrings 2.74.1, gggenomes 1.1.3, and ggplot2 4.0.3. The *E. coli* BL21-AI chromosome (NZ_CP047231.1; GCF_009832985.1) and *E. coli* BW25113 chromosome (CP009273.1) were downloaded in FASTA format. BL21-AI (844,110–866,781 bp or 838,322-871,221 bp) and BW25113 (878,248–890,352 bp) loci were defined, and features fully within each interval were selected. Coordinates were converted to locus-relative positions by subtracting the respective locus start, and feature types were mapped to fixed colors for plotting. Using Biostrings, locus sequences were extracted and GC content in the BL21-AI locus was calculated in sliding windows of 1,000 bp stepped every 200 bp; genome-wide minimum, mean, and maximum GC were obtained from the BL21-AI chromosome using 1,000 bp windows stepped every 1,000 bp, with GC defined as the fraction of G+C bases per window and window midpoints expressed in absolute genomic coordinates. Genomic features were visualized with gggenomes, and homologous genes between BW25113 and BL21-AI loci were connected using a links table plotted with geom_link. GC content in BL21-AI was shown in a separate ggplot2 panel as a filled area plot, with values above the genome-wide mean filled in red and values below the mean in blue, horizontal lines at genome-wide minimum, mean, and maximum GC, and axes displaying GC content as percentage and absolute genomic position aligned to the locus.

### Global GC content analysis

Coding sequences were obtained from Prodigal annotations of concatenated bacterial genome assemblies (132383 replicons)(*65–67*). Each gene was classified as defense-associated when its replicon-specific master identifier occurred in either the DefenseFinder-derived mapping or PADLOC output; all remaining genes were classified as non-defense. Defense-associated and non-defense genes were encoded as `1` and `0`, respectively. Putative operons were defined as consecutive genes on the same strand separated by intergenic distances of ≤ 100bp. An operon was classified as defense-associated if it contained at least one defense-associated gene. GC content was calculated for each gene and operon as the fraction of G and C nucleotides, with values ranging from 0 to 1. Mean GC content was calculated independently for each replicon. Feature-specific GC deviation was then calculated as:

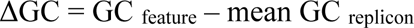

where positive and negative values indicate GC enrichment and depletion, respectively, relative to the corresponding replicon. GC distributions of defense-associated and non-defense genes and operons were compared separately using normalized kernel-density estimates, such that the area under each group-specific curve equaled 1. This enabled distributional comparisons independent of unequal group sizes.

### msDNA isolation and visualization

A 1:100 dilution of ON culture was grown in a flask with 25 ml of LB. The culture was grown for 2h at 37°C (or 30°C) to reach OD_600_ = 0.6. msDNA was extracted from cell lysates using a Monarch plasmid miniprep (NEB, England) from 2 ml of culture of cells (OD_600_ = 0,6) encoding the Eco1 system (BL21-AI, BL21-AI*Δhns*, BL21-AI*ΔstpA*, BL21-AI*ΔhnsΔstpA*) and eluted in a volume of 100 ul. 5 ul of the purified msDNA was then analyzed on 15% PAGE-TBE urea gels, with a 1× TBE running buffer that was heated to ∼50°C before loading. The gels were run (45 min at 200 V) in a TBE running buffer. Gels were stained with Sybr Gold (Thermo Fisher) and imaged on a Gel Doc imager.

### Monitoring bacterial growth in liquid culture

Individual colonies of *E. coli* strains were cultured in LB medium to the stationary phase and then diluted with fresh LB medium to an OD_600_ of approximately 0.06. Cultures were grown at 37 °C (or 30 °C) until reaching an OD_600_ of approximately 0.4. A 150 µl aliquot of each culture, further diluted to an OD_600_ of approximately 0.01, was transferred to 96-well plates and incubated at 37 °C (or 30 °C) with vigorous shaking. Optical density was monitored for 20 h using the EnSpire Multimode Plate Reader (PerkinElmer, USA). Growth curves were plotted based on the collected data.

### Phage liquid culture infection experiments

Overnight bacterial cultures were diluted 100-fold in LB medium with appropriate antibiotics and grown at 37°C. The medium was supplemented with 10 mM CaCl_2_, and 5 mM MgCl_2_. After OD_600_ reached 0.6, 200 μl aliquots were transferred to 96-well plates and infected at the desired MOI. Optical density was monitored for 20 h using the EnSpire Multimode Plate Reader (PerkinElmer, USA). All infections were performed in three biological replicates.

### Phage plaque assays

Double-layer plaque assays were performed with *E. coli* host strains grown to stationary phase at 37°C in LB. A total of 200 μl of the culture was mixed with 10 ml top agar (LB, 0.5% agar, 5 mM MgSO_4_, 10 mM CaCl_2_, 10 mM maltose and the appropriate antibiotic) and poured onto 12 × 12 cm LB-agar plates containing 5 mM MgSO_4_ and the corresponding antibiotic. Ten-fold serial dilutions of phages in liquid LB (10 μl) were spotted onto the plates, and plaques were counted after overnight incubation at 37°C or 30°C. When individual plaques were too small to count, the most concentrated dilution at which no plaques were visible was recorded as containing a single plaque.

### Phage adsorption

Bacterial cultures were grown to mid-exponential phase (OD_600_=0.6) and infected with phage at the MOI=0.01 (approximately 40 µL of diluted phage suspension per 7.5 mL culture). At specified time points (0, 10 and 60 minutes), 1000 µL samples were withdrawn and immediately centrifuged at 12,000 × g for 3 min at 4 °C. A 100 µL aliquot of each supernatant was mixed with 10 µL chloroform, vortexed for 30 s, and incubated at 4 °C. Samples were then centrifuged again under the same conditions. The aqueous phase was collected, and the titer of unadsorbed phages was determined by plaque assay in three independent replicates on BW25113Δ9 culture.

### CRISPRi-mediated silencing of native defense systems

To assess the contribution of native bacterial defense systems to phage restriction, CRISPR interference (CRISPRi) was performed using pFD152, a chloramphenicol-resistant plasmid encoding dCas9 and a defense-system-targeting spacer under anhydrotetracycline (aTc)-inducible control. Targeting constructs were introduced into relevant *E. coli* ECOR strain backgrounds, including ECOR28, ECOR34, and their respective *Δhns* derivatives, by transformation. Transformants were selected on chloramphenicol-containing medium (34 μg ml^−1). Overnight liquid cultures carrying either defense-system-targeting or non-targeting control pFD152 constructs were grown in the presence or absence of 200 ng ml^−1 aTc. Induction was used to activate dCas9 and spacer expression and thereby silence the cognate native defense system. Cultures were subsequently used for efficiency-of-plating (EOP) experiments as described separately. Following overnight incubation of double-layer agar plates at 37°C, phage titers were compared between induced and uninduced cultures. The experimental design leveraged the role of H-NS as a repressor of bacterial immune functions. Accordingly, CRISPRi-mediated silencing was expected to produce limited or no change in phage titer in parental ECOR backgrounds, in which defense activity is comparatively repressed. In contrast, a greater increase in phage titer upon induction was expected in the corresponding *Δhns* backgrounds, where native defense-system activity is enhanced. An increased phage titer on induced, defense-system-targeted cultures relative to uninduced and non-targeting controls was interpreted as evidence that the targeted native defense system contributes to restriction of the tested phage.

### RT-qPCR analysis of gene expression

Total RNA was extracted using the RNA solo kit (Evrogen, Russia), and RNA quality was assessed by agarose gel electrophoresis. RNA quantitation was performed with the Qubit RNA BR Assay Kit. Complementary DNA synthesis was performed using 0.25 µg of total RNA, MMLV reverse transcriptase (Evrogen) and random hexamers according to the manufacturer’s instructions. RT–qPCR was performed using qPCRmix-HS SYBR (Evrogen) on a QuantStudio 5 Real-Time qPCR system (Applied Biosystems). rpoC (RNA polymerase β′-subunit) served as housekeeping gene. For qPCR data analysis, ΔΔCt values were calculated for the two qPCR targets for each replicate. The 2^^-ΔΔCt^ values were counted, after which the 2^^-ΔΔCt^ values of the experimental strains were normalized to the wild-type 2^^-ΔΔCt^ values. The mean fold change of three biological replicates was plotted.

### RNA-seq analysis

RNA-seq reads from *E. coli* ECOR28, ECOR34, BL21-AI, and MG1655 were aligned to strain-specific reference genomes with STAR. Gene-level counts were generated from sorted BAM files using featureCounts and the corresponding GTF/GFF annotations. Paired-end fragments were counted using strand-specific settings appropriate for directional libraries. ECOR28 and ECOR34 comparisons included three biological replicates per condition; BL21-AI and MG1655 samples were analyzed in their respective pairwise comparisons. Differential expression was assessed using DESeq2. Genes with fewer than 10 total counts in a comparison were excluded. Size-factor normalization, dispersion estimation, and negative-binomial modeling were performed using condition as the design variable. *P* values were adjusted by the Benjamini– Hochberg method, and log_2_ fold changes were shrinkage-estimated. Genes were considered differentially expressed at adjusted *P* <0.05 and ∣ log_2_FC ∣≥ 0.58 for ECOR28, ECOR34, and BL21-AI analyses, or ∣ log_2_FC ∣≥ 0.30 for the MG1655 double mutant-versus-WT analysis. Regularized-log-transformed counts were used for principal-component analysis, and normalized counts were used for heat maps. Volcano plots show shrunken log_2_ fold changes versus −log_10_(adjusted *P* values). Genes located within annotated prophage and defense-system loci were identified by genomic-coordinate overlap and highlighted in the plots. For BL21-AI display figures, plotted fold changes and significance values were visually truncated; these display limits did not alter statistical testing or differential-expression calls.

### ONT sequencing and genome assembly of the ECOR strains

Genomic DNA of ECOR28 and ECOR34 was purified from 2 mL of overnight cultures grown in LB at 37 °C with Monarch Genomic DNA Purification Kit (NEB). Total DNA libraries were prepared from DNA using the Native Barcoding Kit 24 V14 (SQK-NBD114-24) with enrichment of long fragments using the Long Fragment Buffer according to the manufacturer’s instructions. DNA library was sequenced using R10.4.1 flow cell (FLO-MIN114) on the MinION device with MinKNOW v23.11.2.Oxford Nanopore Technologies (ONT) long-read datasets from *E. coli* strains ECOR28 and ECOR34 were assembled *de novo* using Unicycler in long-read-only mode. Draft assemblies were subsequently polished with NextPolish v1.4.1 using the corresponding ONT long reads. The quality and completeness of the polished assemblies were evaluated using the gVolante web server, based on the coverage of a 440-gene conserved ortholog set. Assembly completeness was reported using BUSCO-style categories: complete (C), single-copy complete (S), duplicated complete (D), fragmented (F), and missing (M) orthologs. Both assemblies showed 100% recovery of complete or partial core genes.

## Acknowledgments

We sincerely thank Seth Shipman for sharing materials and for discussion of the Eco1 system. We thank Igor Granovsky for providing phage T4 and Anna Trofimova for the help with ONT sequencing of ECOR strains. We thank Oksana Kotovskaya for the discussion of the resulst of this work. RNA sequencing was performed at the Genomics Core Facility, with support of the internal PhD student grants to P.I.

## Authors Contribution

A. I., T.K., and P.I. conceived the study; T.K., and P.I. performed experiments; A. G., I. S., V. E. performed Micro-C and ChIP-seq experiments; P.B., and E.S. performed RNA-sequencing; I.S., V.E., and P.I. analyzed NGS data; A.A. constructed Sxt*Δ0.3* phage; A. I., K. S., and E. N. secured resources and supervised the study; P.I., and T.K. prepared illustrations; A.I. and T.K. wrote the manuscript. All authors reviewed, edited, and approved the final version.

## Data, code, and materials availability

All unique bacterial strains, phages, and plasmids generated in this study are available from the lead contact without restriction. Further information and requests for resources and reagents should be directed to and will be fulfilled by the lead contact, Artem Isaev.

## Declaration of generative AI and AI-assisted technologies in the manuscript preparation process

During the preparation of this work, the author(s) used DeepSeek to assist with grammar checking of the assembled manuscript. The author(s) reviewed and edited the output as needed and take full responsibility for the content of the published article.

