## Extended data for "H-NS silences antiviral immunity through 3D chromatin compaction"

**Extended data figure 1**

3D chromosome compactization, GC content and gene expression levels of the prophages and defense loci in the BL21-AI, Δ*hns*, and Δ*hns*/Δ*stpA* derivatives

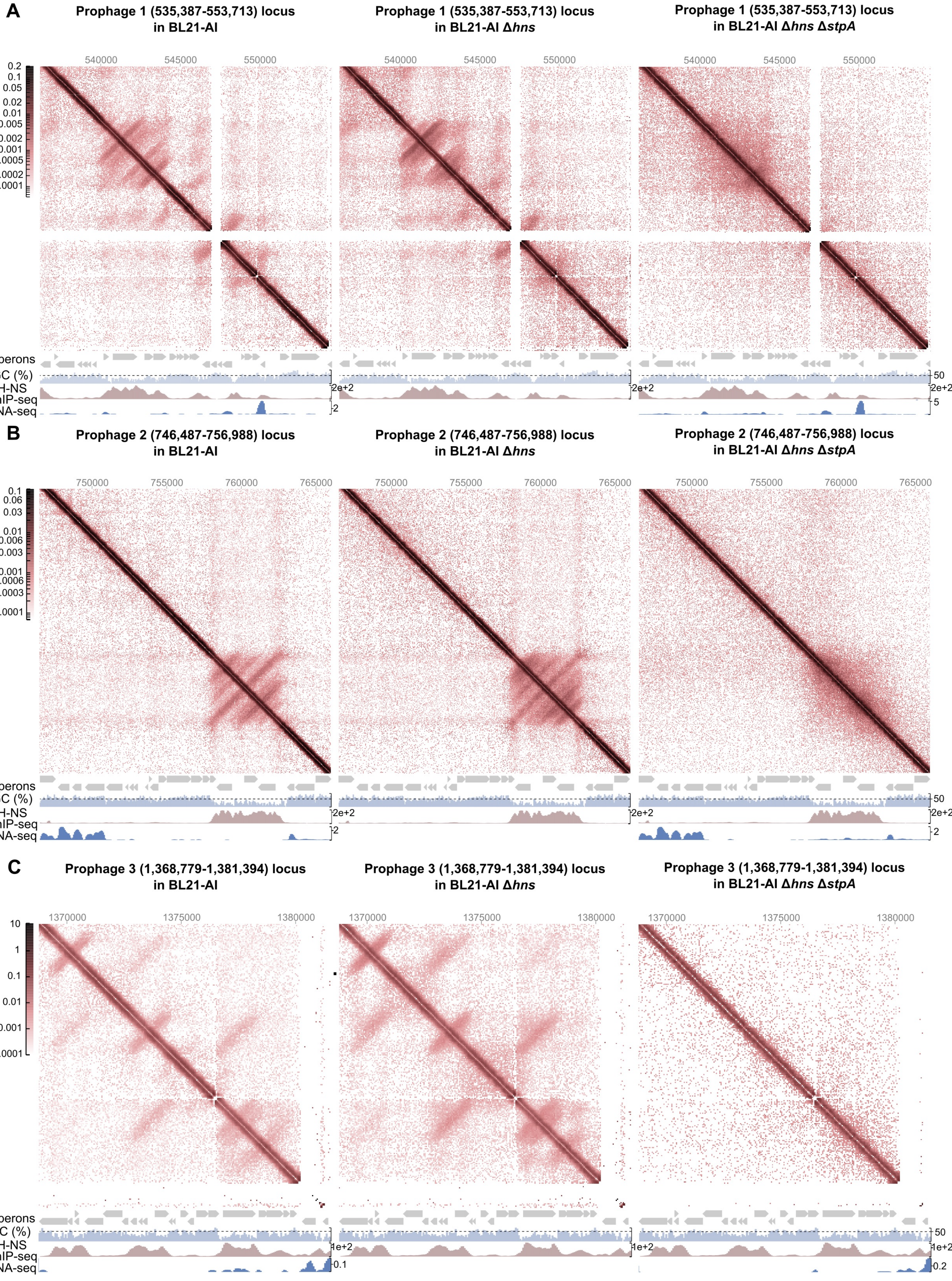

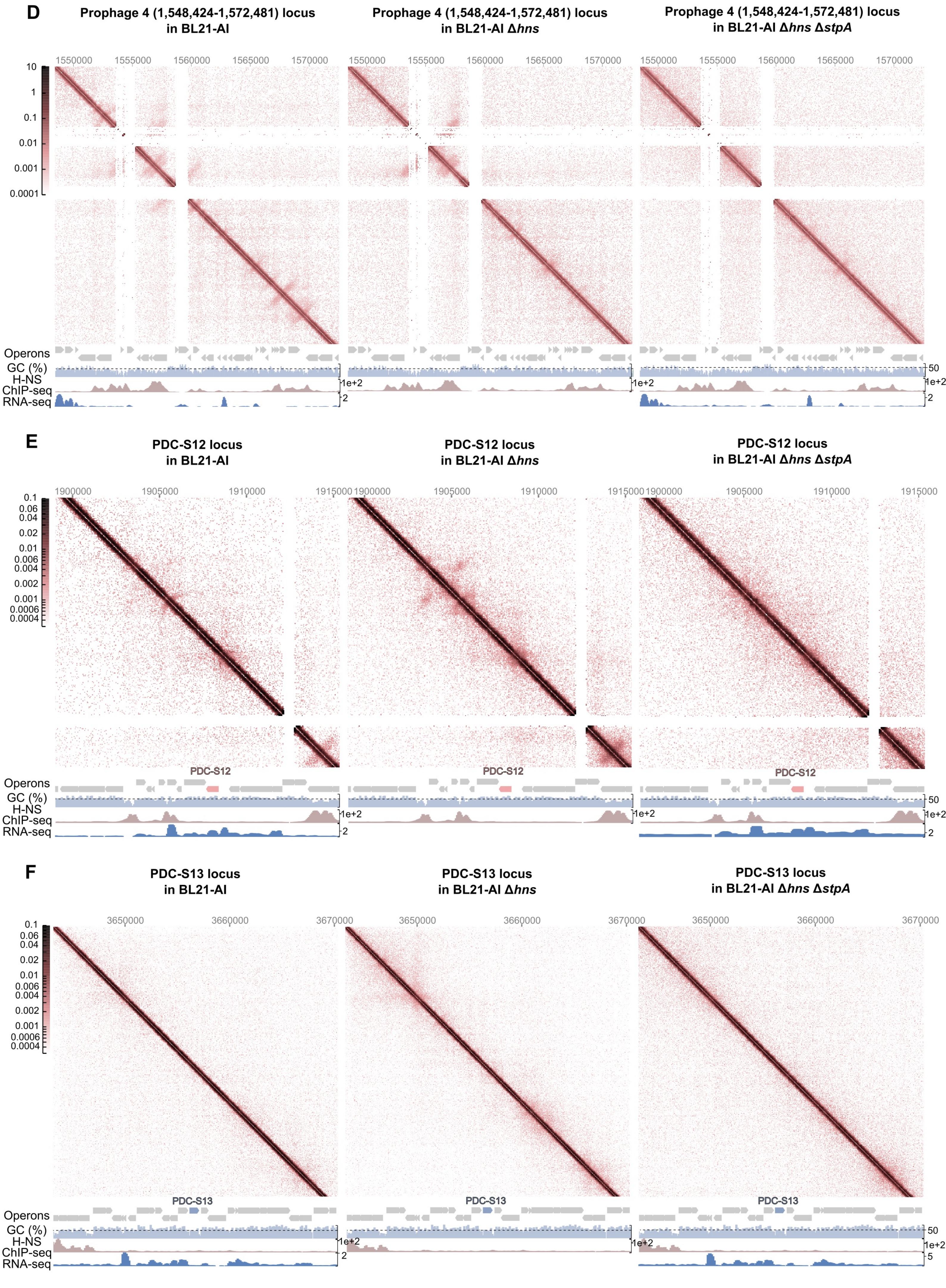
D
Prophage 4 (1,548,424-1,572,481) locus in BL21-AI
Prophage 4 (1,548,424-1,572,481) locus in BL21-AI Δhns
Prophage 4 (1,548,424-1,572,481) locus in BL21-AI Δhns ΔstpA
1550000
1555000
1560000
1565000
1570000
10
1
0.1
0.01
0.001
0.0001
Operons
GC (%)
H-NS ChIP-seq
RNA-seq
50
1e+2
2
E
PDC-S12 locus in BL21-AI
PDC-S12 locus in BL21-AI Δhns
PDC-S12 locus in BL21-AI Δhns ΔstpA
1900000
1905000
1910000
1915000
0.1
0.06
0.04
0.02
0.01
0.006
0.004
0.002
0.001
0.0006
0.0004
PDC-S12
Operons
GC (%)
H-NS ChIP-seq
RNA-seq
50
1e+2
2
F
PDC-S13 locus in BL21-AI
PDC-S13 locus in BL21-AI Δhns
PDC-S13 locus in BL21-AI Δhns ΔstpA
3650000
3660000
3670000
0.1
0.06
0.04
0.02
0.01
0.006
0.004
0.002
0.001
0.0006
0.0004
PDC-S13
Operons
GC (%)
H-NS ChIP-seq
RNA-seq
50
1e+2
2
5

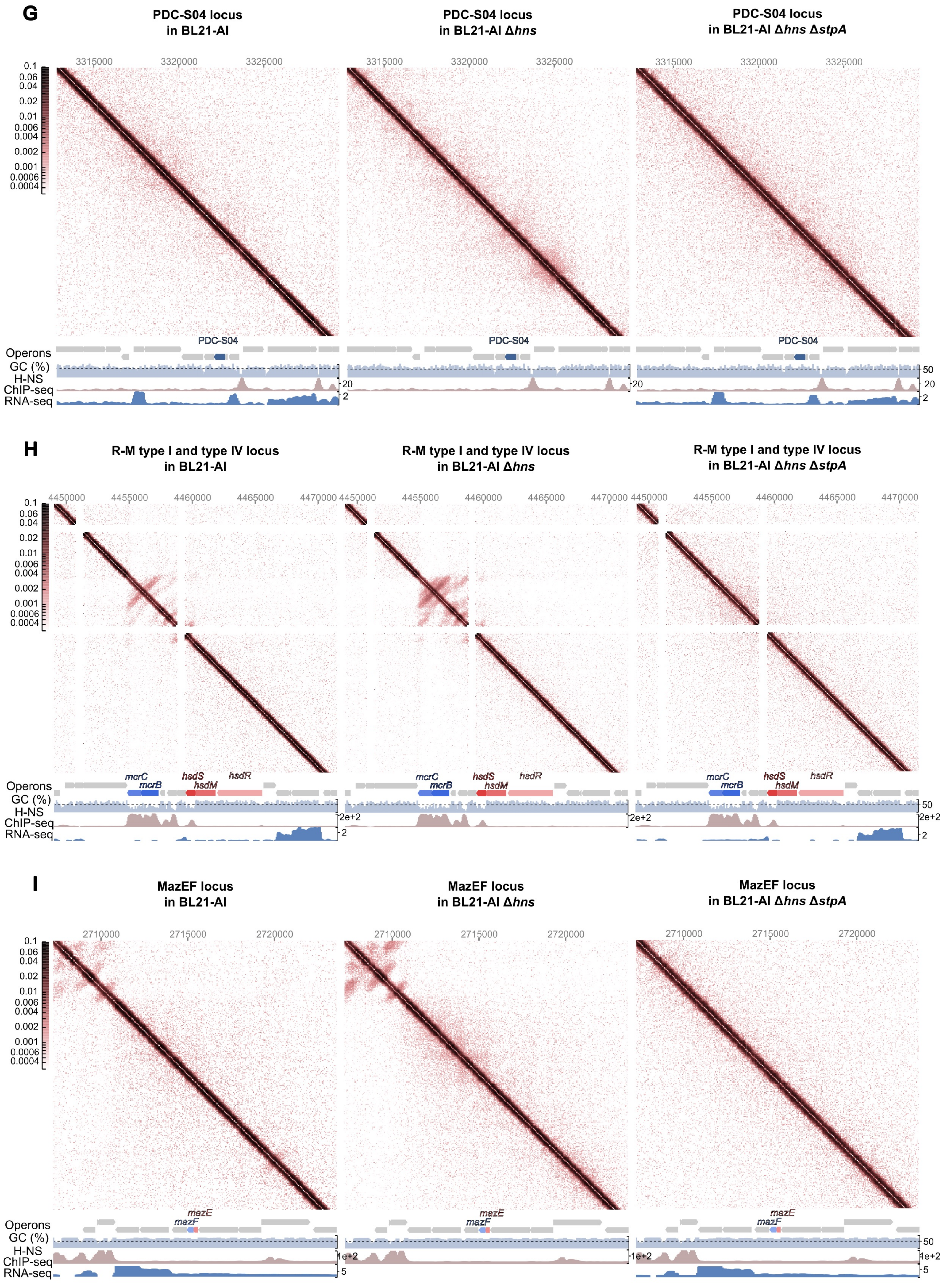
G
PDC-S04 locus
in BL21-AI
PDC-S04 locus
in BL21-AI Δhns
PDC-S04 locus
in BL21-AI Δhns ΔstpA
3315000
3320000
3325000
0.1
0.06
0.04
0.02
0.01
0.006
0.004
0.002
0.001
0.0006
0.0004
PDC-S04
Operons
GC (%)
H-NS
ChIP-seq
RNA-seq
50
20
2
H
R-M type I and type IV locus
in BL21-AI
R-M type I and type IV locus
in BL21-AI Δhns
R-M type I and type IV locus
in BL21-AI Δhns ΔstpA
4450000
4455000
4460000
4465000
4470000
mcrC
mcrB
hsdS
hsdM
hsdR
2e+2
I
MazEF locus
in BL21-AI
MazEF locus
in BL21-AI Δhns
MazEF locus
in BL21-AI Δhns ΔstpA
2710000
2715000
2720000
mazE
mazF
1e+2
5

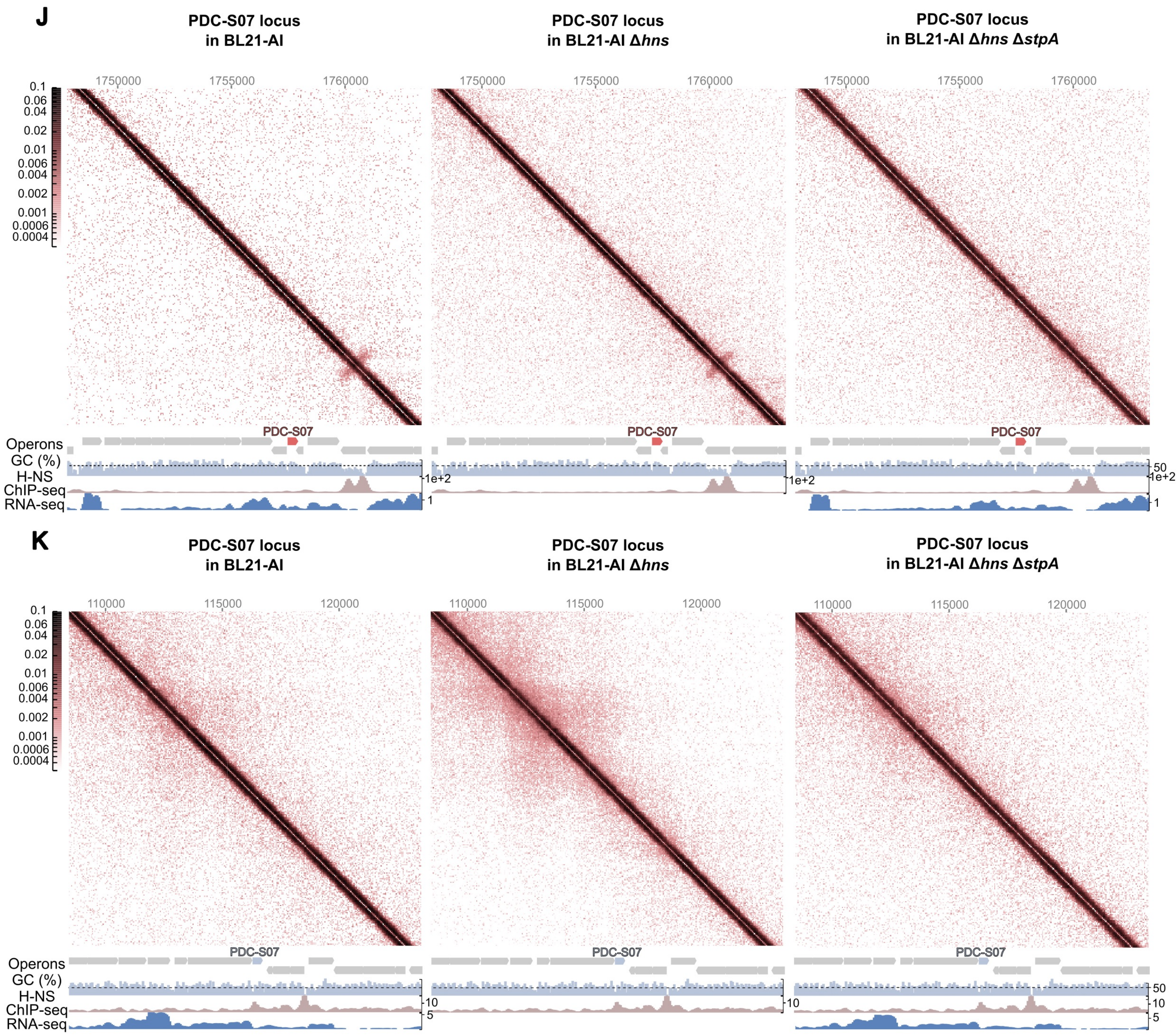
J
PDC-S07 locus in BL21-AI
PDC-S07 locus in BL21-AI Δhns
PDC-S07 locus in BL21-AI Δhns ΔstpA
1750000
1755000
1760000
0.1
0.06
0.04
0.02
0.01
0.006
0.004
0.002
0.001
0.0006
0.0004
PDC-S07
Operons
GC (%)
H-NS
ChIP-seq
RNA-seq
50
1e+2
1
K
PDC-S07 locus in BL21-AI
PDC-S07 locus in BL21-AI Δhns
PDC-S07 locus in BL21-AI Δhns ΔstpA
110000
115000
120000
10
5

**Extended data figure 2**

3D chromosome compactization, GC content and gene expression levels of the prophages and defense loci in the MG1655, Δ*hns*, and Δ*hns*/Δ*stpA* derivatives

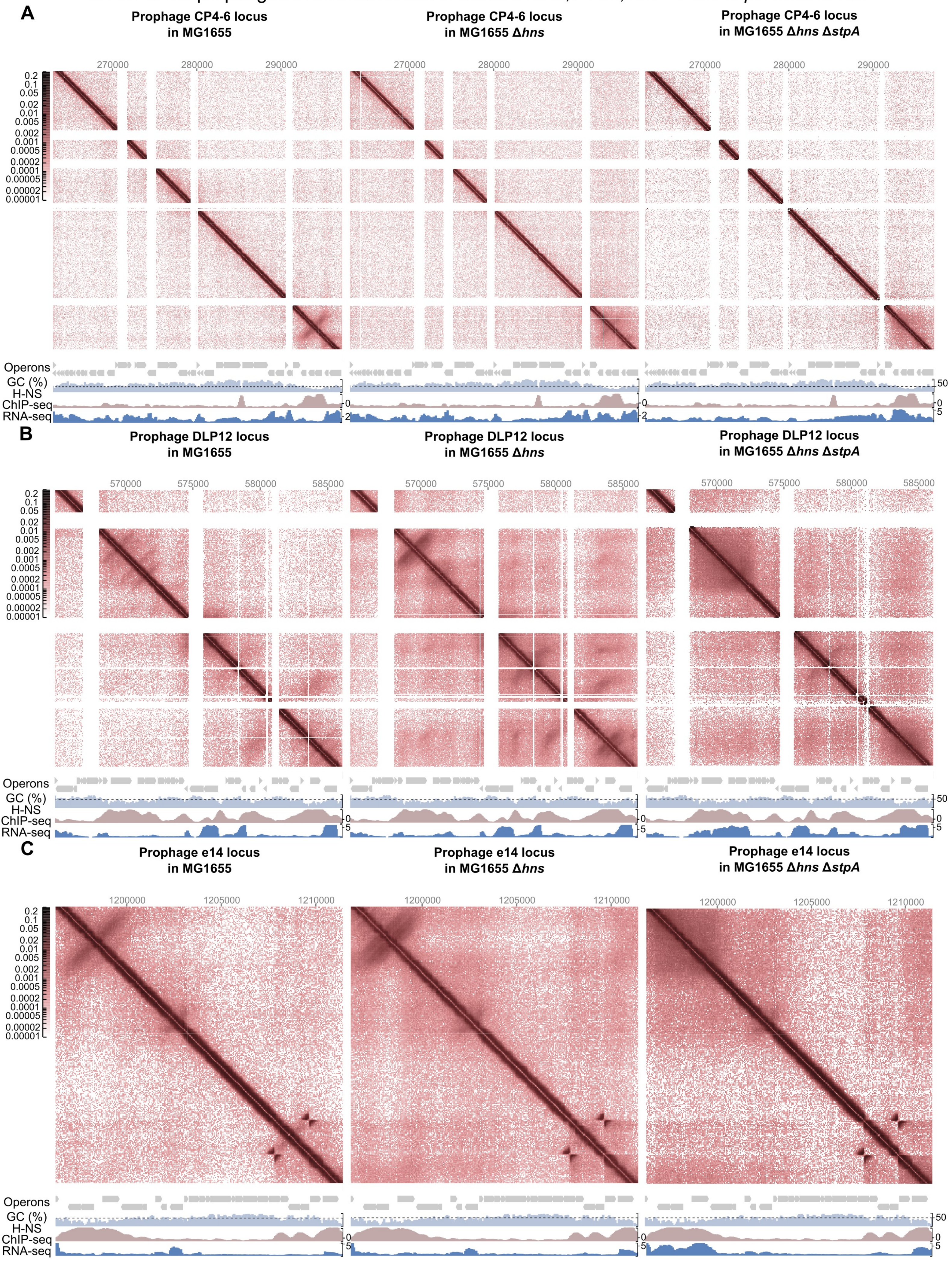

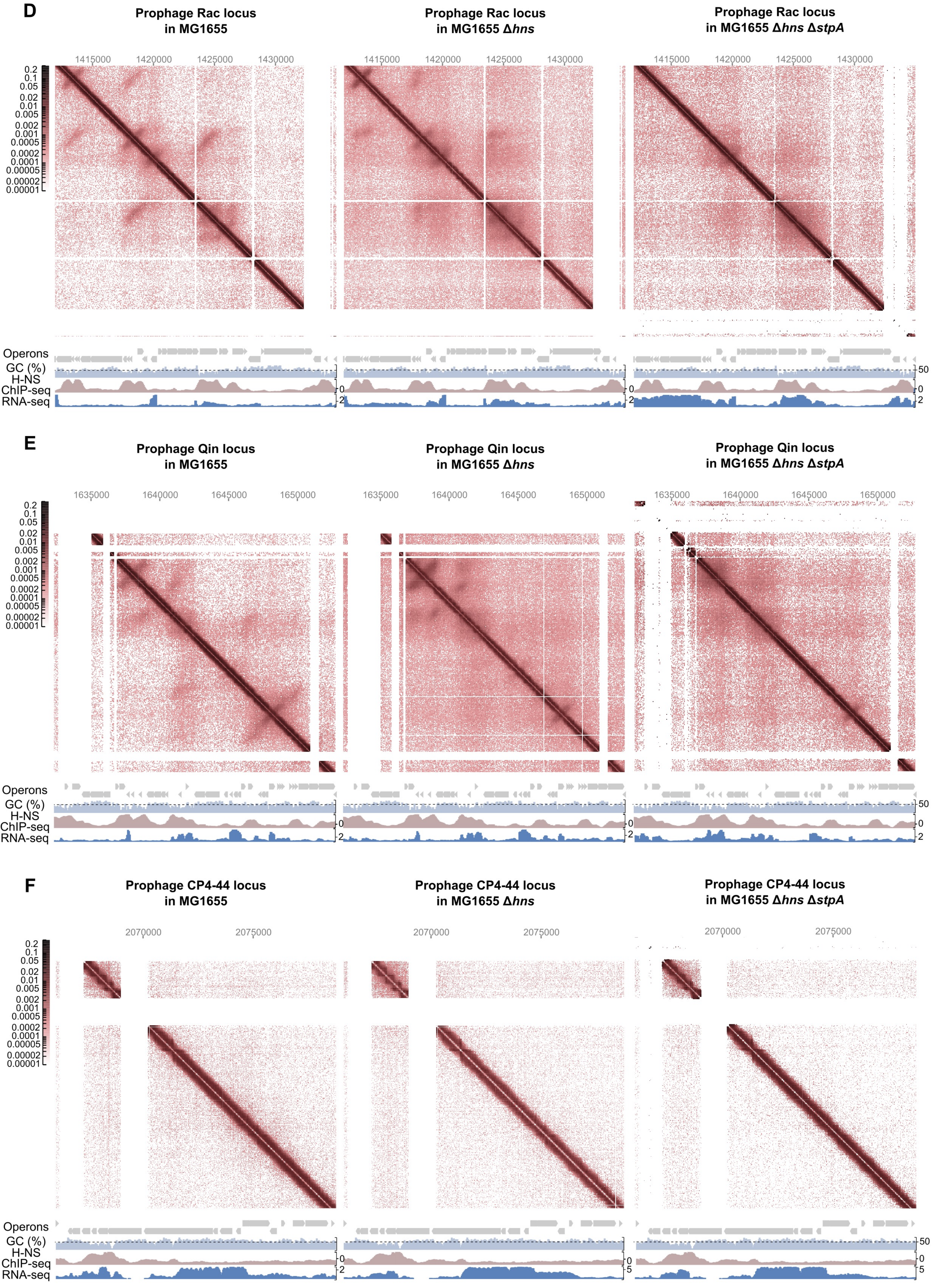
D
Prophage Rac locus in MG1655
Prophage Rac locus in MG1655 Δhns
Prophage Rac locus in MG1655 Δhns ΔstpA
1415000
1420000
1425000
1430000
0.2
0.1
0.05
0.02
0.01
0.005
0.002
0.001
0.0005
0.0002
0.0001
0.00005
0.00002
0.00001
Operons
GC (%)
H-NS ChIP-seq
RNA-seq
50
0
2
E
Prophage Qin locus in MG1655
Prophage Qin locus in MG1655 Δhns
Prophage Qin locus in MG1655 Δhns ΔstpA
1635000
1640000
1645000
1650000
F
Prophage CP4-44 locus in MG1655
Prophage CP4-44 locus in MG1655 Δhns
Prophage CP4-44 locus in MG1655 Δhns ΔstpA
2070000
2075000
5

G

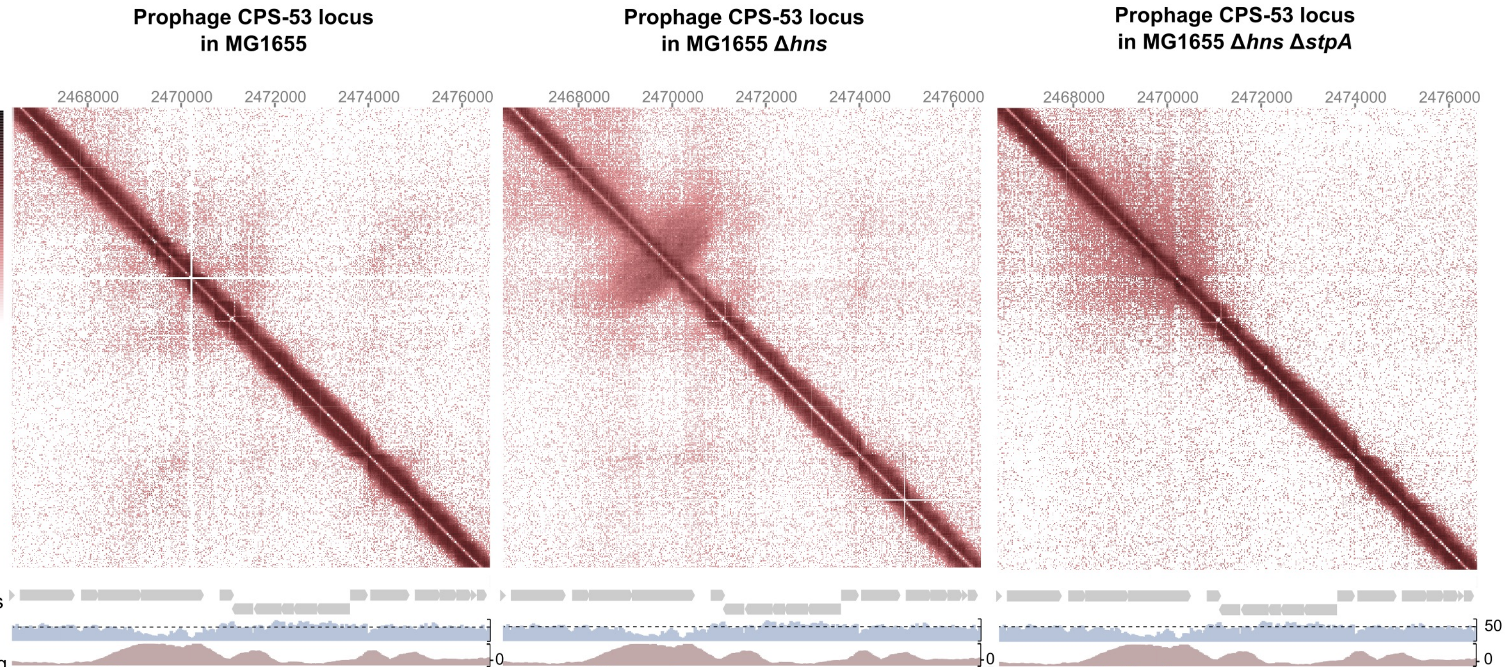

Prophage CPS-53 locus in MG1655
Prophage CPS-53 locus in MG1655 Δhns
Prophage CPS-53 locus in MG1655 Δhns ΔstpA
2468000
2470000
2472000
2474000
2476000
0.2
0.1
0.05
0.02
0.01
0.005
0.002
0.001
0.0005
0.0002
0.0001
0.00005
0.00002
0.00001
Operons
GC (%)
H-NS
ChIP-seq
RNA-seq
50
0
2
5


H

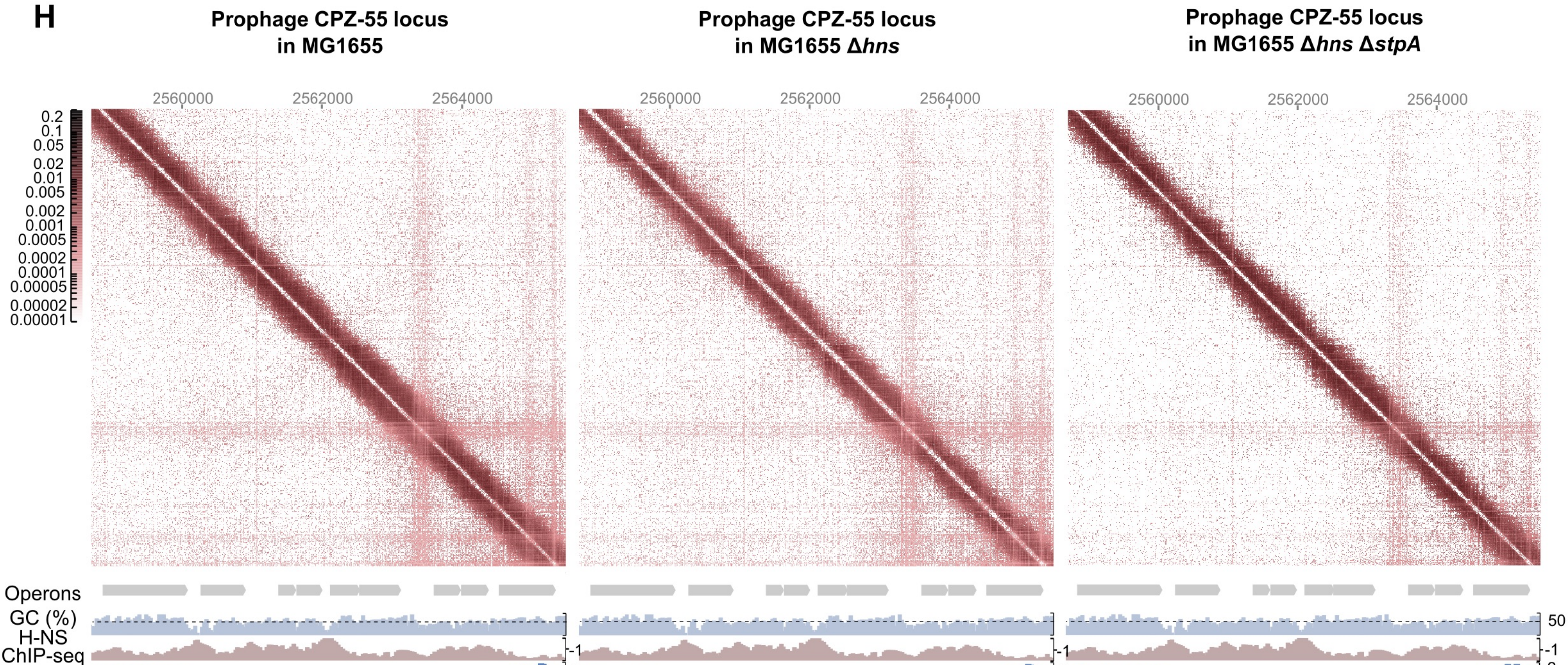

Prophage CPZ-55 locus in MG1655
Prophage CPZ-55 locus in MG1655 Δhns
Prophage CPZ-55 locus in MG1655 Δhns ΔstpA
2560000
2562000
2564000
0.2
0.1
0.05
0.02
0.01
0.005
0.002
0.001
0.0005
0.0002
0.0001
0.00005
0.00002
0.00001
Operons
GC (%)
H-NS
ChIP-seq
RNA-seq
50
-1
2


I

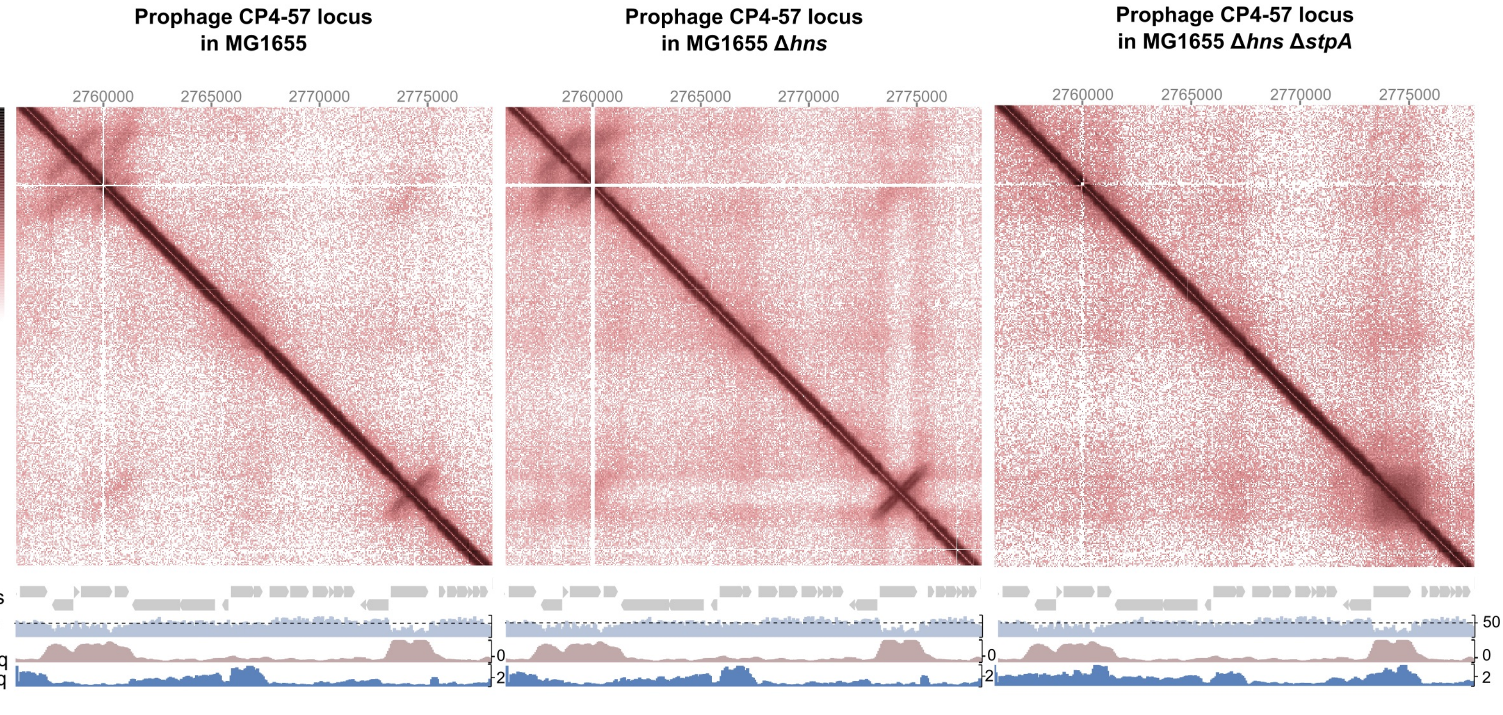

Prophage CP4-57 locus in MG1655
Prophage CP4-57 locus in MG1655 Δhns
Prophage CP4-57 locus in MG1655 Δhns ΔstpA
2760000
2765000
2770000
2775000
0.2
0.1
0.05
0.02
0.01
0.005
0.002
0.001
0.0005
0.0002
0.0001
0.00005
0.00002
0.00001
Operons
GC (%)
H-NS
ChIP-seq
RNA-seq
50
0
2

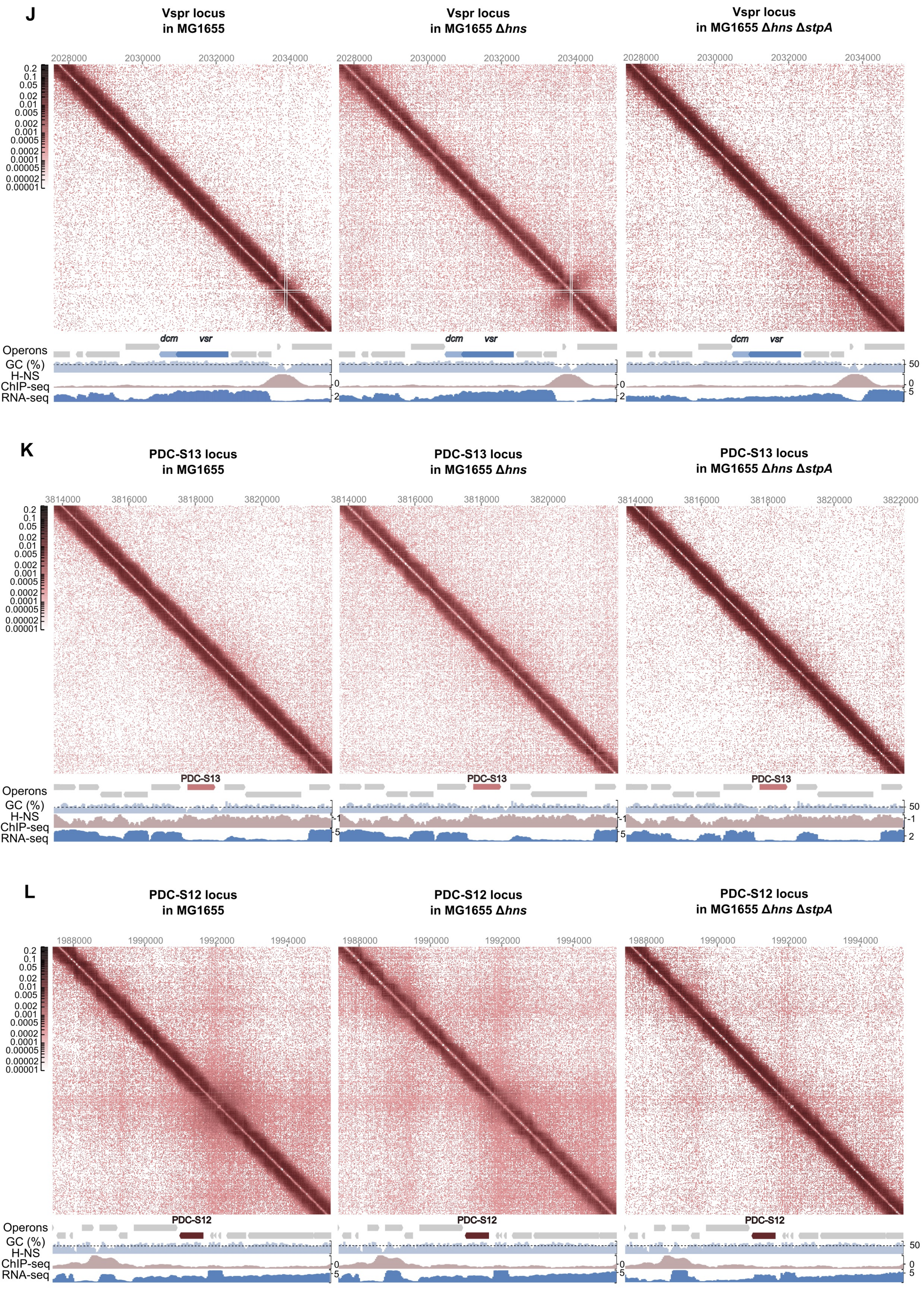

J
Vspr locus
in MG1655
Vspr locus
in MG1655 Δ*hns*
Vspr locus
in MG1655 Δ*hns* Δ*stpA*
2028000
2030000
2032000
2034000
0.2
0.1
0.05
0.02
0.01
0.005
0.002
0.001
0.0005
0.0002
0.0001
0.00005
0.00002
0.00001
dcm
vsr
Operons
GC (%)
H-NS
ChIP-seq
RNA-seq
50
0
2
5
K
PDC-S13 locus
in MG1655
PDC-S13 locus
in MG1655 Δ*hns*
PDC-S13 locus
in MG1655 Δ*hns* Δ*stpA*
3814000
3816000
3818000
3820000
3822000
PDC-S13
-1
L
PDC-S12 locus
in MG1655
PDC-S12 locus
in MG1655 Δ*hns*
PDC-S12 locus
in MG1655 Δ*hns* Δ*stpA*
1988000
1990000
1992000
1994000
PDC-S12

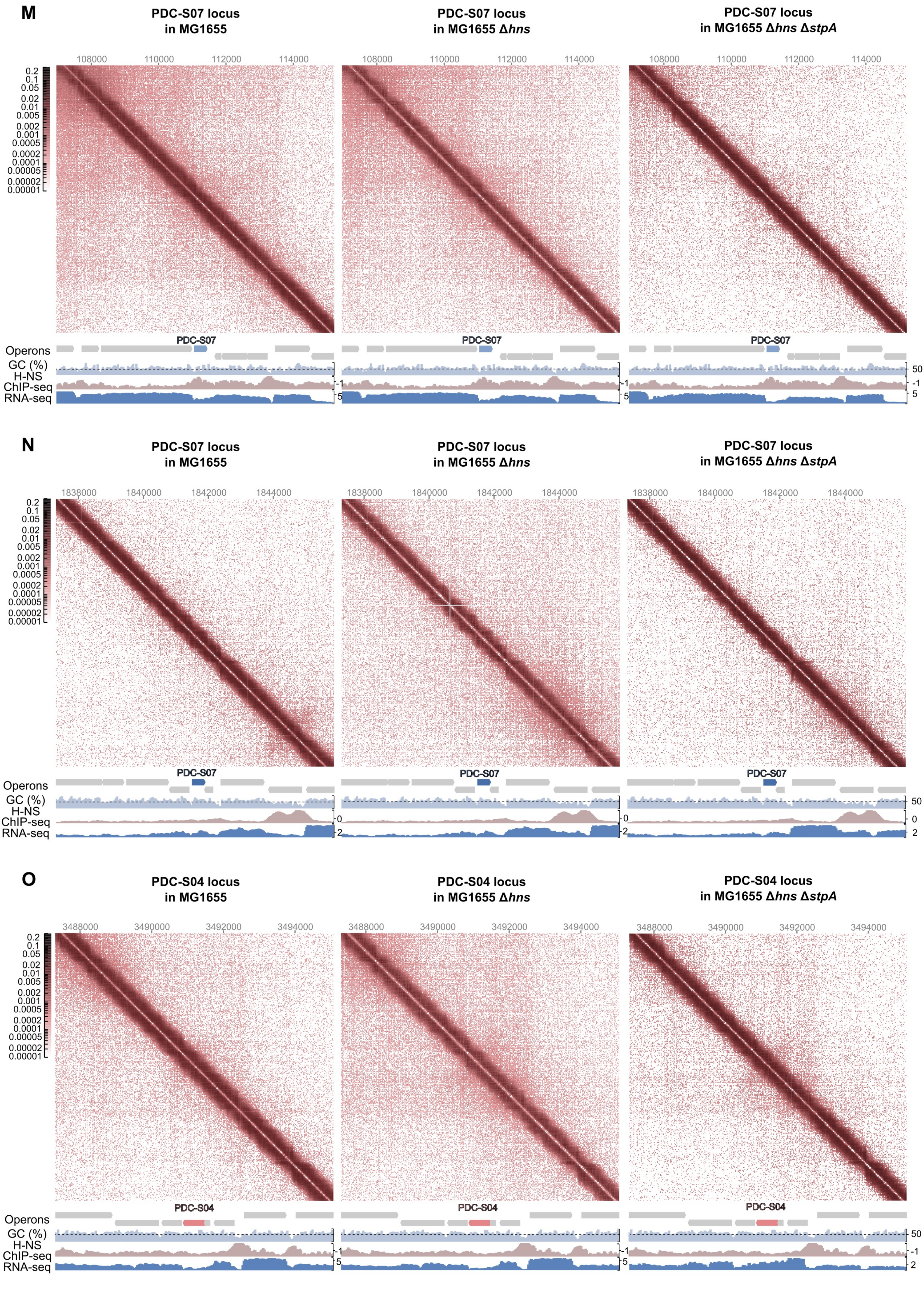

M
PDC-S07 locus in MG1655
PDC-S07 locus in MG1655 Δhns
PDC-S07 locus in MG1655 Δhns ΔstpA
108000
110000
112000
114000
0.2
0.1
0.05
0.02
0.01
0.005
0.002
0.001
0.0005
0.0002
0.0001
0.00005
0.00002
0.00001
Operons
PDC-S07
GC (%)
H-NS
ChIP-seq
RNA-seq
50
-1
5
N
PDC-S07 locus in MG1655
PDC-S07 locus in MG1655 Δhns
PDC-S07 locus in MG1655 Δhns ΔstpA
1838000
1840000
1842000
1844000
PDC-S07
0
2
O
PDC-S04 locus in MG1655
PDC-S04 locus in MG1655 Δhns
PDC-S04 locus in MG1655 Δhns ΔstpA
3488000
3490000
3492000
3494000
PDC-S04
50
-1
5
2

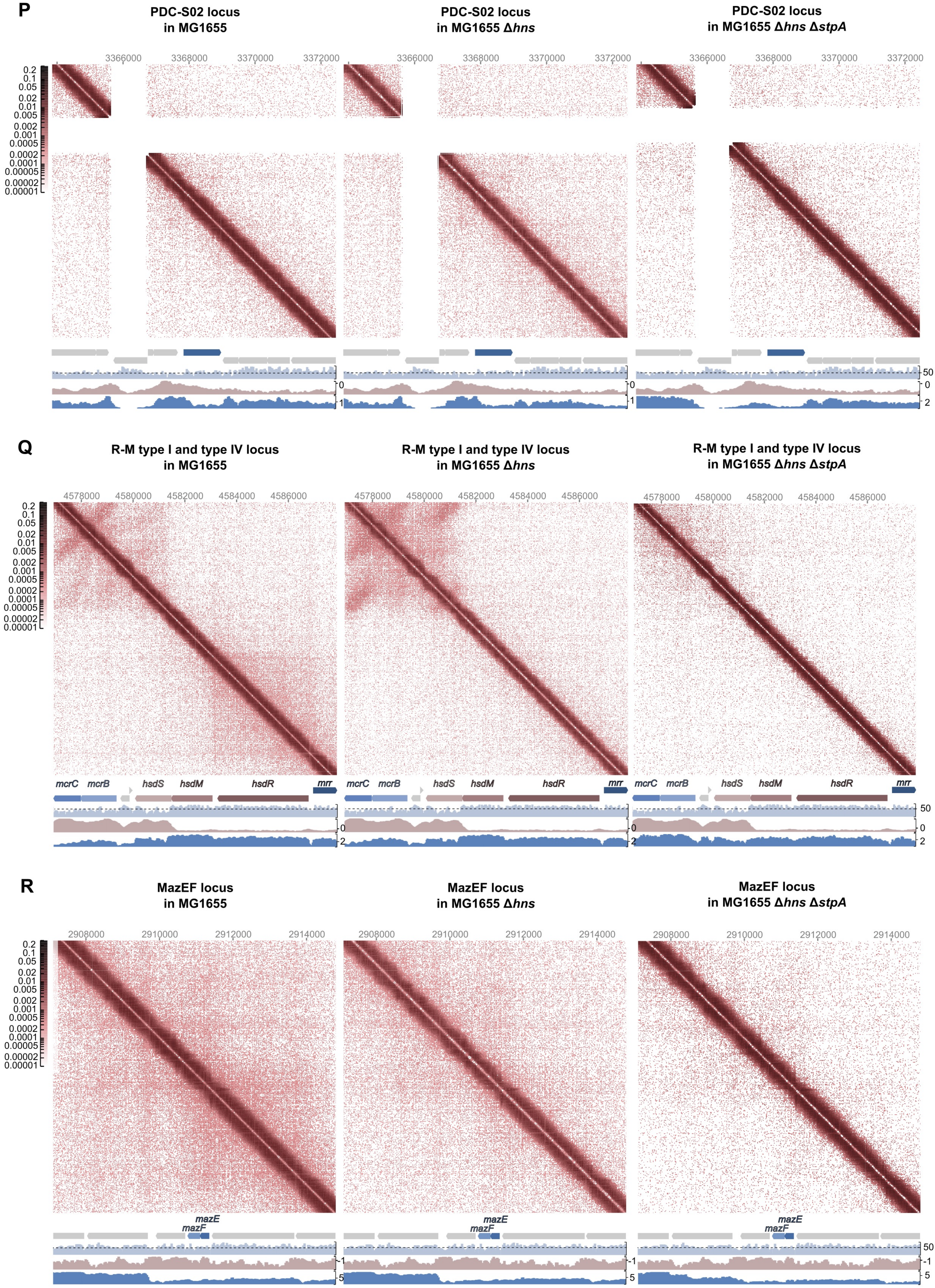
P
PDC-S02 locus in MG1655
PDC-S02 locus in MG1655 Δhns
PDC-S02 locus in MG1655 Δhns ΔstpA
3366000
3368000
3370000
3372000
Q
R-M type I and type IV locus in MG1655
R-M type I and type IV locus in MG1655 Δhns
R-M type I and type IV locus in MG1655 Δhns ΔstpA
4578000
4580000
4582000
4584000
4586000
mcrC
mcrB
hsdS
hsdM
hsdR
mrr
R
MazEF locus in MG1655
MazEF locus in MG1655 Δhns
MazEF locus in MG1655 Δhns ΔstpA
2908000
2910000
2912000
2914000
mazE
mazF

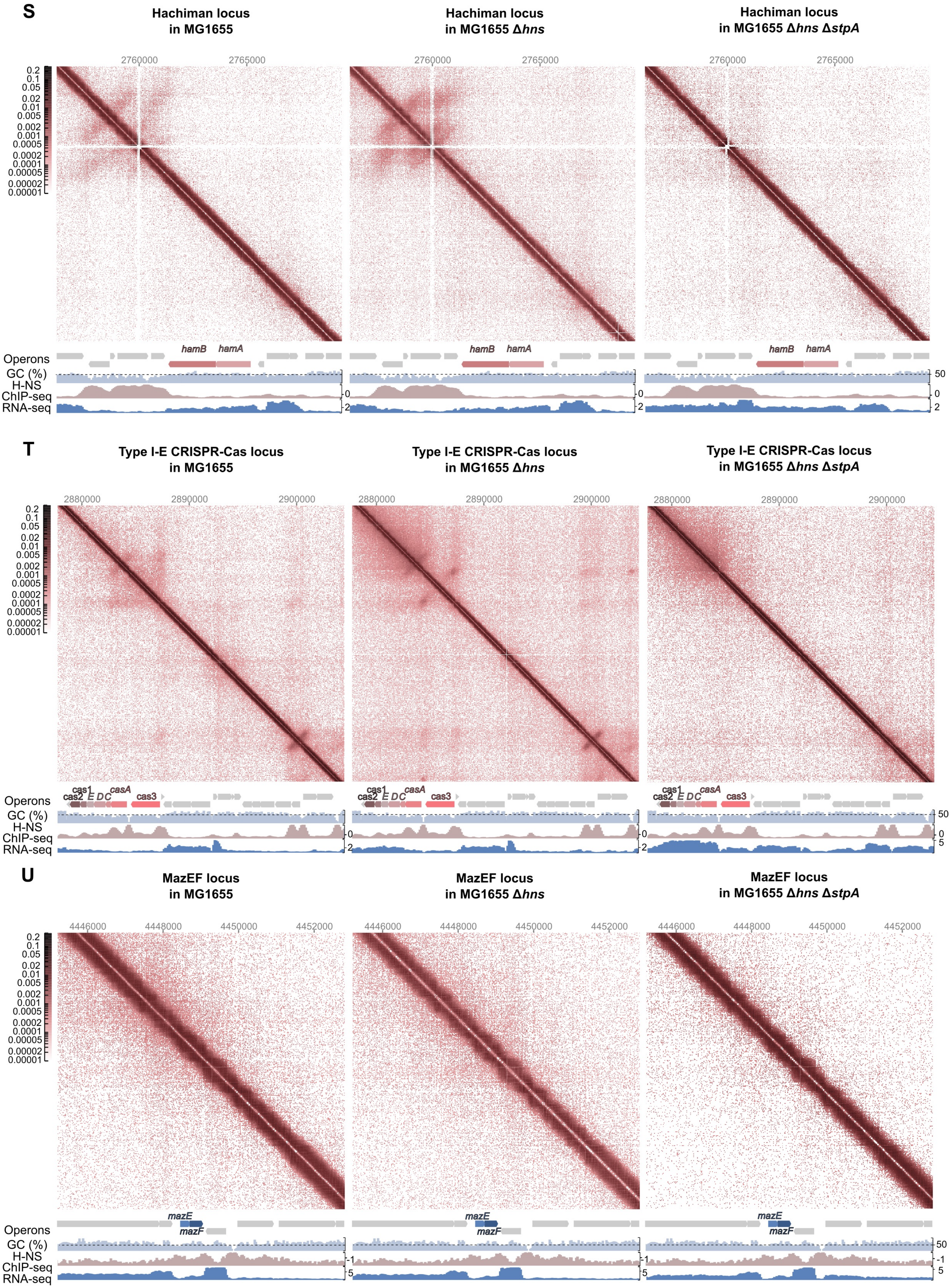
S
Hachiman locus
in MG1655
Hachiman locus
in MG1655 Δhns
Hachiman locus
in MG1655 Δhns ΔstpA
2760000
2765000
0.2
0.1
0.05
0.02
0.01
0.005
0.002
0.001
0.0005
0.0002
0.0001
0.00005
0.00002
0.00001
Operons
hamB
hamA
GC (%)
H-NS
ChIP-seq
RNA-seq
50
0
2
T
Type I-E CRISPR-Cas locus
in MG1655
Type I-E CRISPR-Cas locus
in MG1655 Δhns
Type I-E CRISPR-Cas locus
in MG1655 Δhns ΔstpA
2880000
2890000
2900000
cas1
cas2
E
D
C
casA
cas3
5
U
MazEF locus
in MG1655
MazEF locus
in MG1655 Δhns
MazEF locus
in MG1655 Δhns ΔstpA
4446000
4448000
4450000
4452000
mazE
mazF
-1

### Extended data figure 3

3D chromosome compactization, GC content and gene expression levels of the prophages and defense loci in the ECOR28, Δ*hns*, and Δ*hns*/Δ*stpA* derivatives

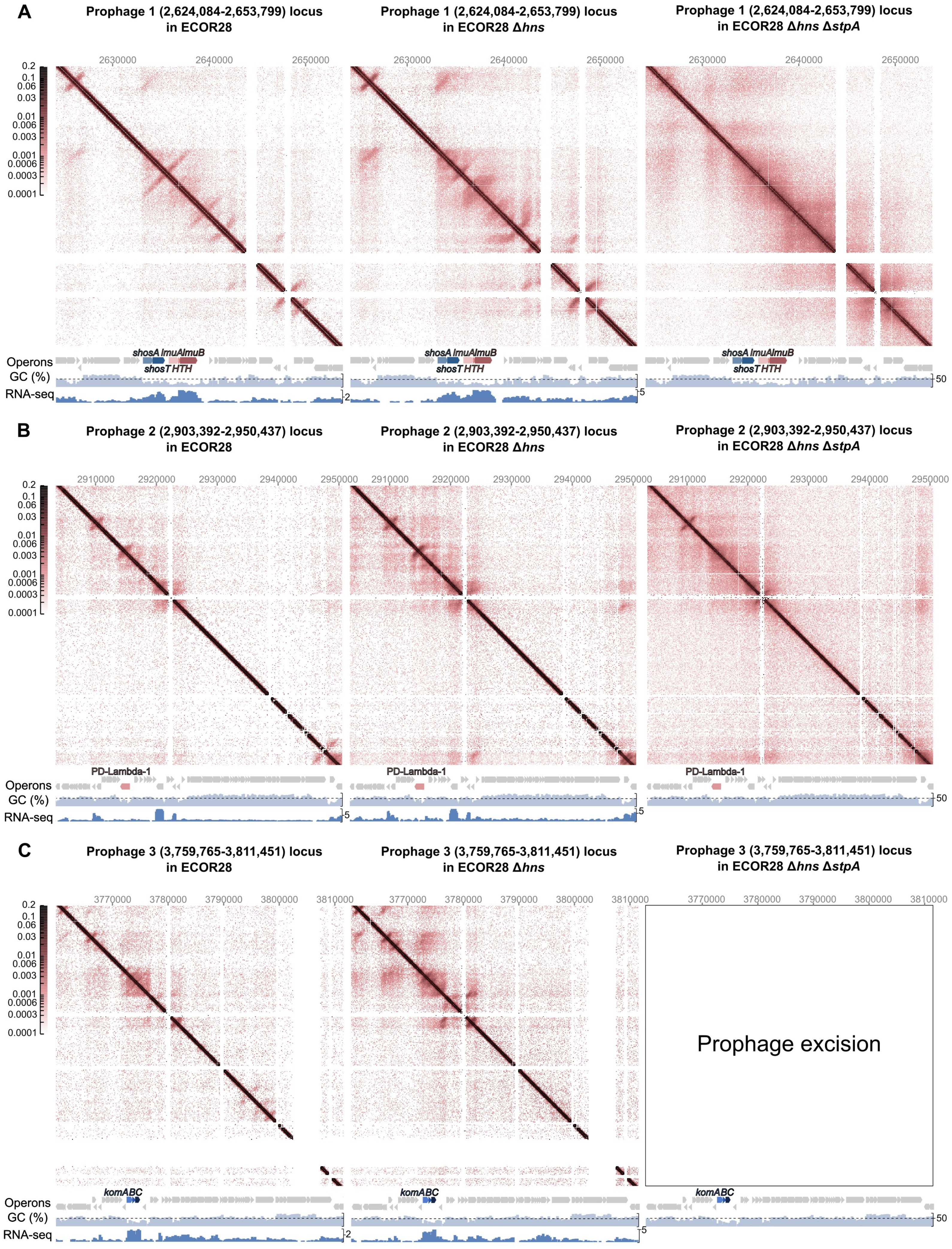

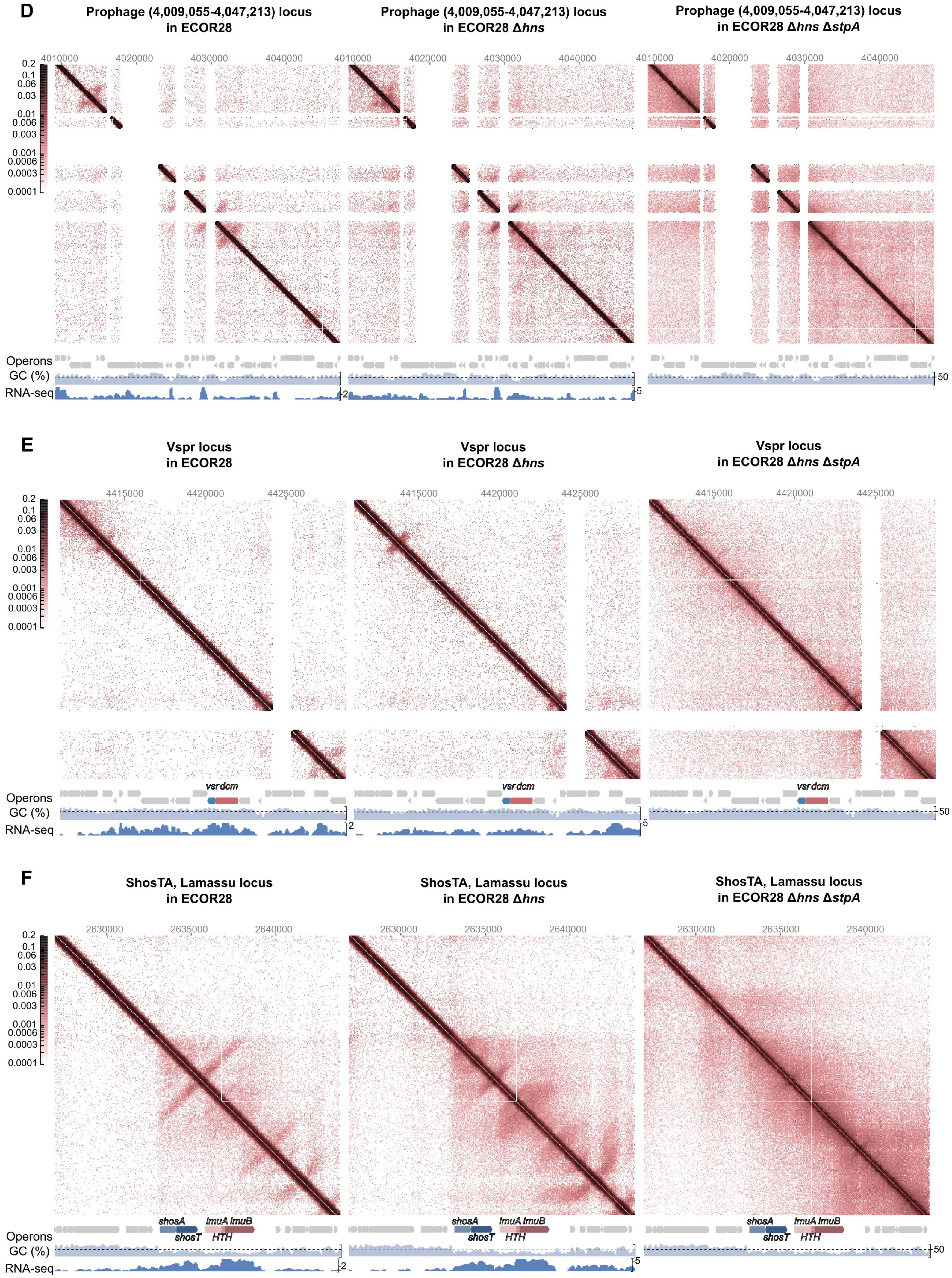
D
Prophage (4,009,055-4,047,213) locus in ECOR28
Prophage (4,009,055-4,047,213) locus in ECOR28 Δhns
Prophage (4,009,055-4,047,213) locus in ECOR28 Δhns ΔstpA
4010000 4020000 4030000 4040000
0.2 0.1 0.06 0.03 0.01 0.006 0.003 0.001 0.0006 0.0003 0.0001
Operons
GC (%)
RNA-seq
50
E
Vspr locus in ECOR28
Vspr locus in ECOR28 Δhns
Vspr locus in ECOR28 Δhns ΔstpA
4415000 4420000 4425000
vsr dcm
F
ShosTA, Lamassu locus in ECOR28
ShosTA, Lamassu locus in ECOR28 Δhns
ShosTA, Lamassu locus in ECOR28 Δhns ΔstpA
2630000 2635000 2640000
shosA shosT lmuA lmuB HTH

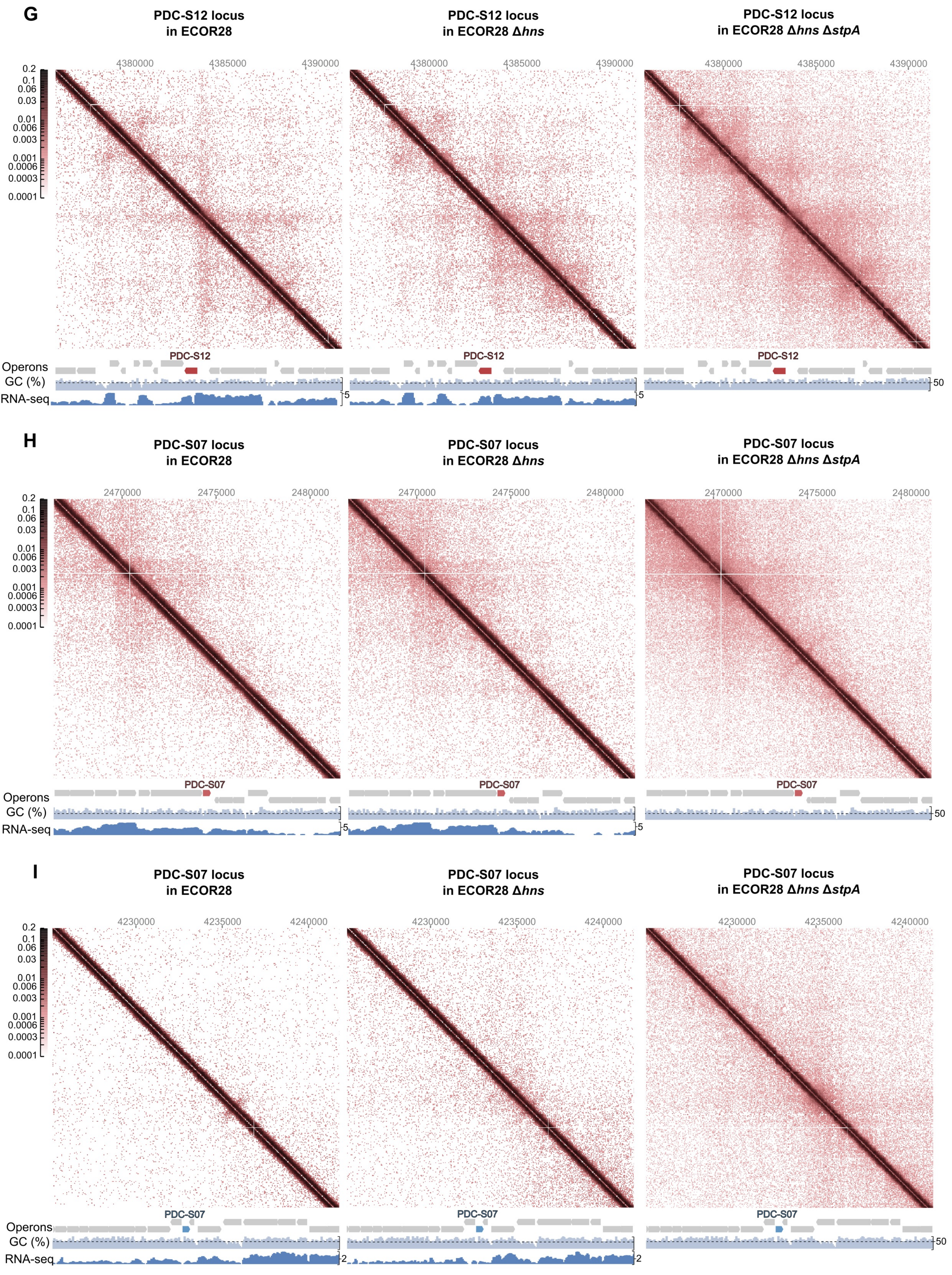
G
PDC-S12 locus in ECOR28
PDC-S12 locus in ECOR28 Δ*hns*
PDC-S12 locus in ECOR28 Δ*hns* Δ*stpA*
4380000
4385000
4390000
0.2
0.1
0.06
0.03
0.01
0.006
0.003
0.001
0.0006
0.0003
0.0001
PDC-S12
Operons
GC (%)
RNA-seq
5
50
H
PDC-S07 locus in ECOR28
PDC-S07 locus in ECOR28 Δ*hns*
PDC-S07 locus in ECOR28 Δ*hns* Δ*stpA*
2470000
2475000
2480000
PDC-S07
I
PDC-S07 locus in ECOR28
PDC-S07 locus in ECOR28 Δ*hns*
PDC-S07 locus in ECOR28 Δ*hns* Δ*stpA*
4230000
4235000
4240000
2

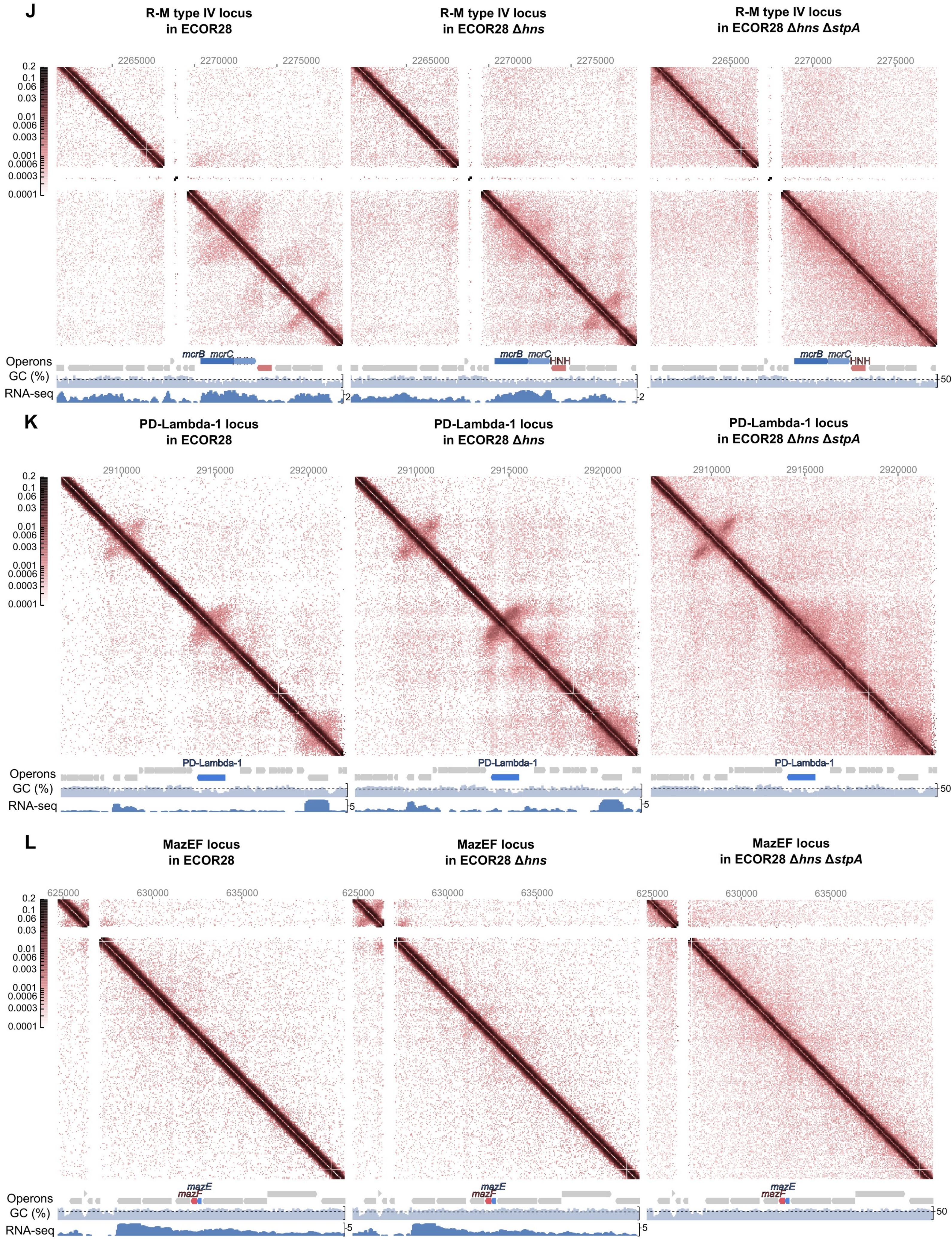

J
R-M type IV locus in ECOR28
R-M type IV locus in ECOR28 Δhns
R-M type IV locus in ECOR28 Δhns ΔstpA
2265000
2270000
2275000
0.2
0.1
0.06
0.03
0.01
0.006
0.003
0.001
0.0006
0.0003
0.0001
mcrB
mcrC
HNH
Operons
GC (%)
RNA-seq
2
50
K
PD-Lambda-1 locus in ECOR28
PD-Lambda-1 locus in ECOR28 Δhns
PD-Lambda-1 locus in ECOR28 Δhns ΔstpA
2910000
2915000
2920000
PD-Lambda-1
5
L
MazEF locus in ECOR28
MazEF locus in ECOR28 Δhns
MazEF locus in ECOR28 Δhns ΔstpA
625000
630000
635000
mazE
mazF

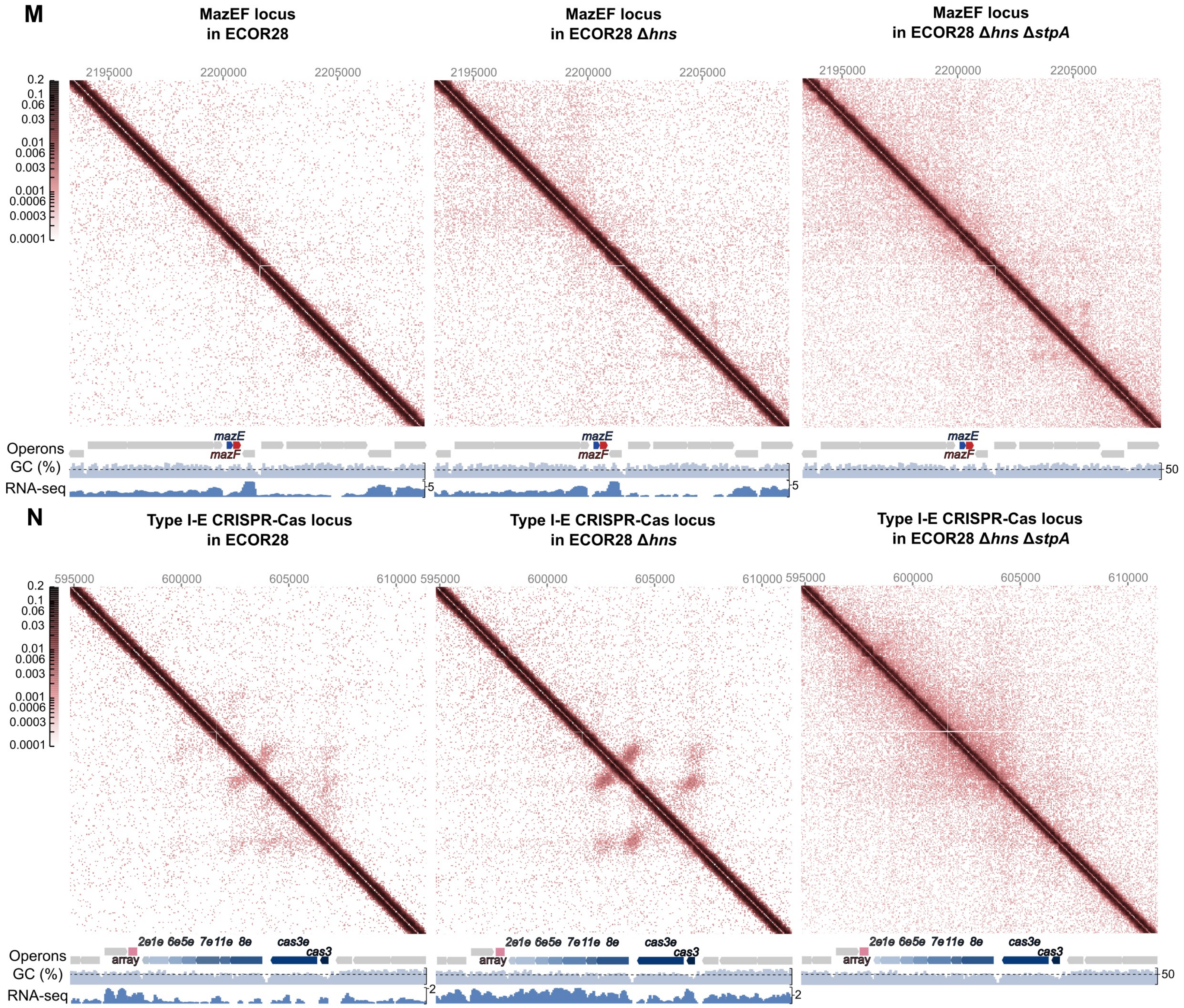

M
MazEF locus in ECOR28
MazEF locus in ECOR28 Δhns
MazEF locus in ECOR28 Δhns ΔstpA
2195000
2200000
2205000
0.2
0.1
0.06
0.03
0.01
0.006
0.003
0.001
0.0006
0.0003
0.0001
Operons
GC (%)
RNA-seq
mazE
mazF
5
50
N
Type I-E CRISPR-Cas locus in ECOR28
Type I-E CRISPR-Cas locus in ECOR28 Δhns
Type I-E CRISPR-Cas locus in ECOR28 Δhns ΔstpA
595000
600000
605000
610000
array
2e 1e 6e 5e 7e 11e 8e
cas3e
cas3
2

**Extended data figure 4**

3D chromosome compactization, GC content and gene expression levels of the prophages and defense loci in the ECOR34, Δ*hns* derivative

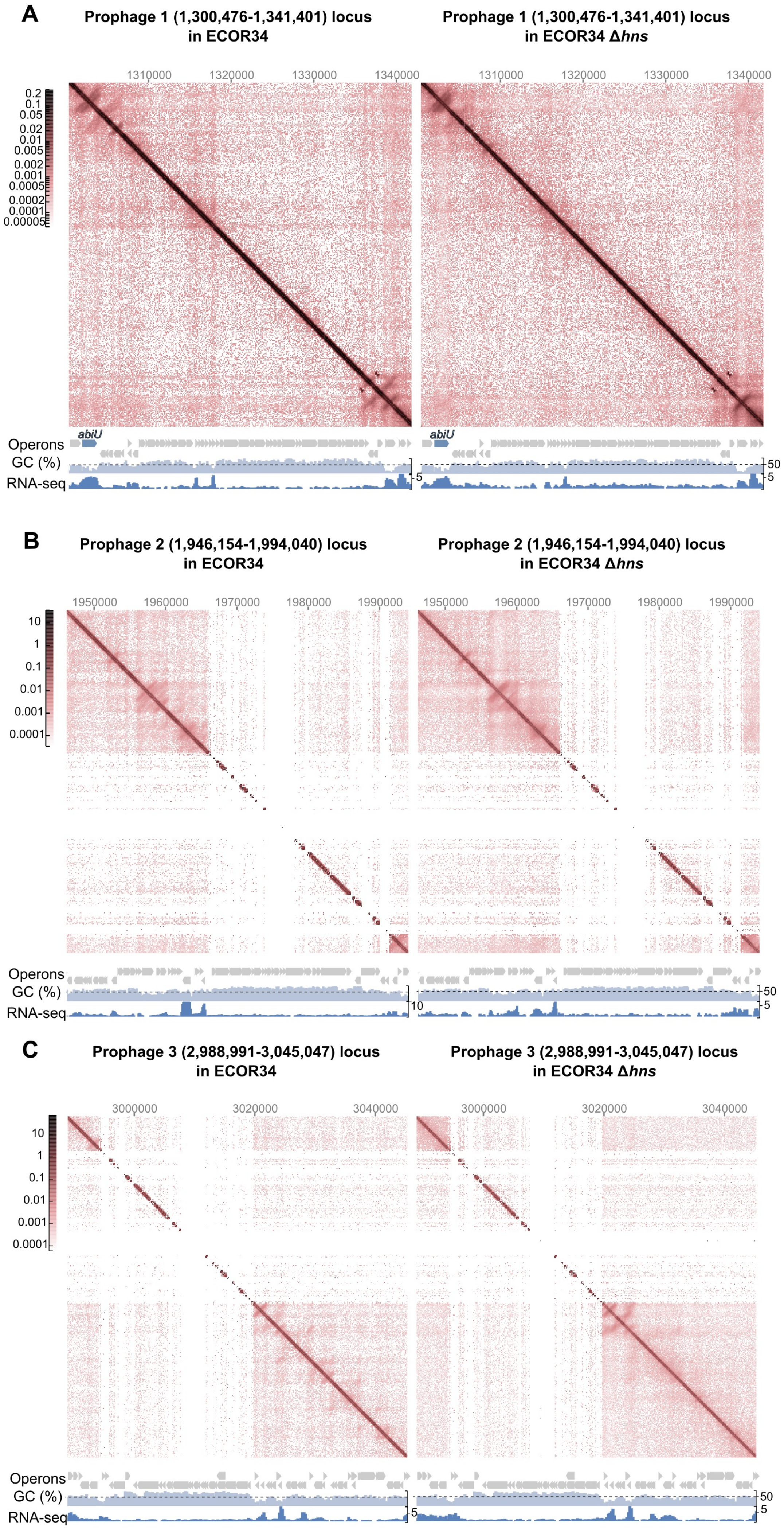

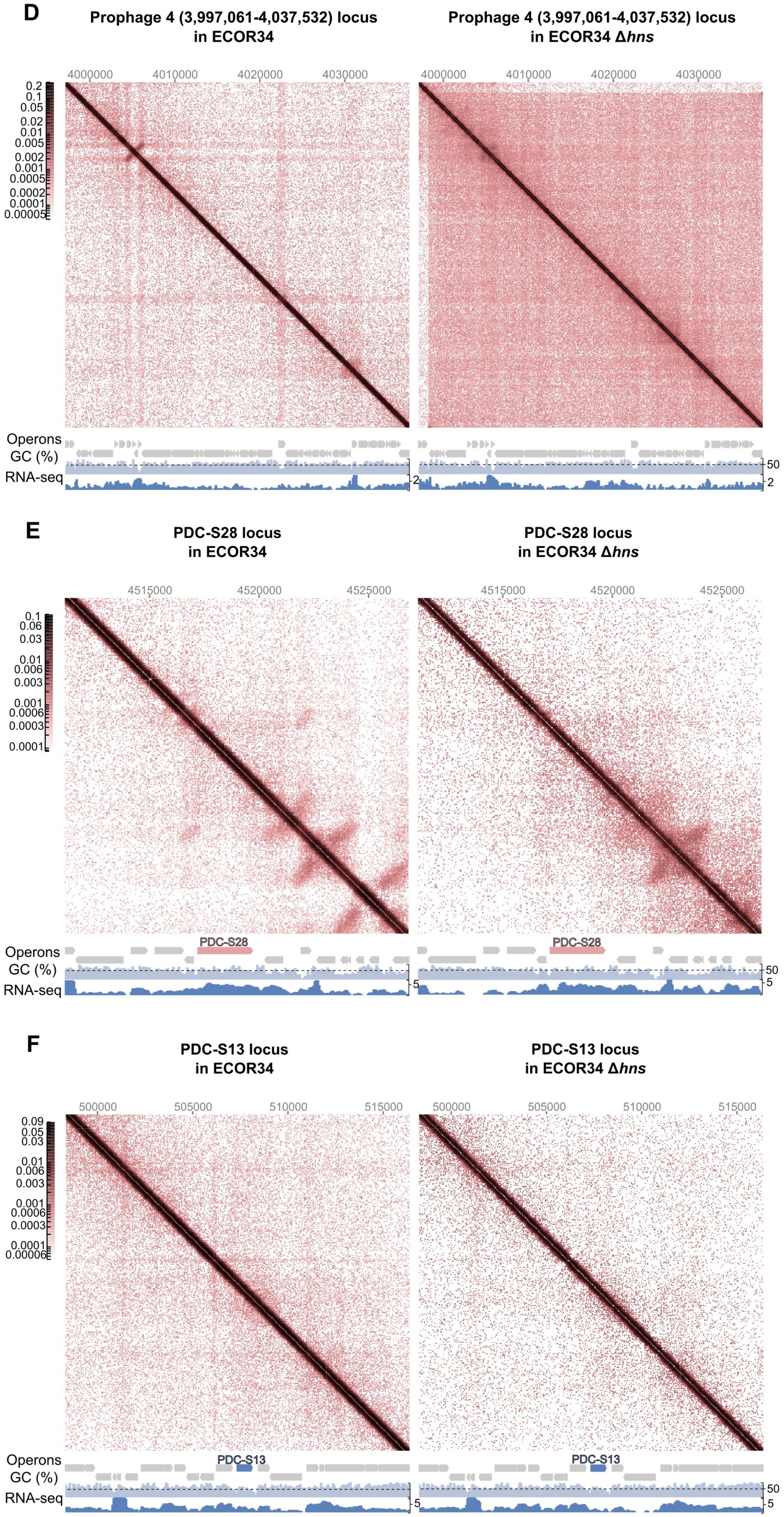
D
Prophage 4 (3,997,061-4,037,532) locus in ECOR34
Prophage 4 (3,997,061-4,037,532) locus in ECOR34 Δhns
4000000
4010000
4020000
4030000
0.2
0.1
0.05
0.02
0.01
0.005
0.002
0.001
0.0005
0.0002
0.0001
0.00005
Operons
GC (%)
RNA-seq
50
2
E
PDC-S28 locus in ECOR34
PDC-S28 locus in ECOR34 Δhns
4515000
4520000
4525000
0.1
0.06
0.03
0.01
0.006
0.003
0.001
0.0006
0.0003
0.0001
PDC-S28
Operons
GC (%)
RNA-seq
50
5
F
PDC-S13 locus in ECOR34
PDC-S13 locus in ECOR34 Δhns
500000
505000
510000
515000
0.09
0.05
0.03
0.01
0.006
0.003
0.001
0.0006
0.0003
0.0001
0.00006
PDC-S13
Operons
GC (%)
RNA-seq
50
5

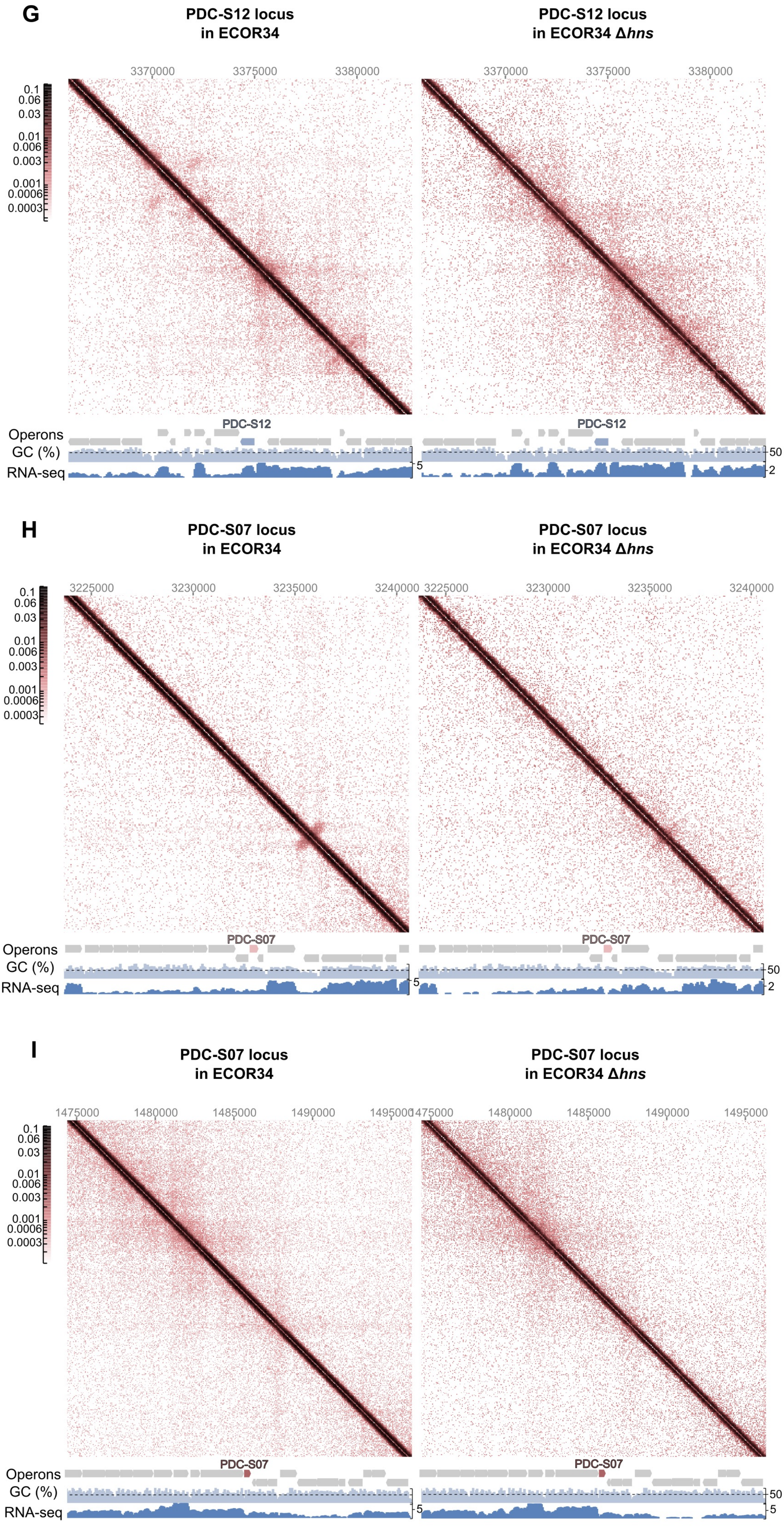

G
PDC-S12 locus in ECOR34
PDC-S12 locus in ECOR34 Δhns
3370000
3375000
3380000
0.1
0.06
0.03
0.01
0.006
0.003
0.001
0.0006
0.0003
Operons
GC (%)
RNA-seq
PDC-S12
50
5
2
H
PDC-S07 locus in ECOR34
PDC-S07 locus in ECOR34 Δhns
3225000
3230000
3235000
3240000
PDC-S07
I
PDC-S07 locus in ECOR34
PDC-S07 locus in ECOR34 Δhns
1475000
1480000
1485000
1490000
1495000

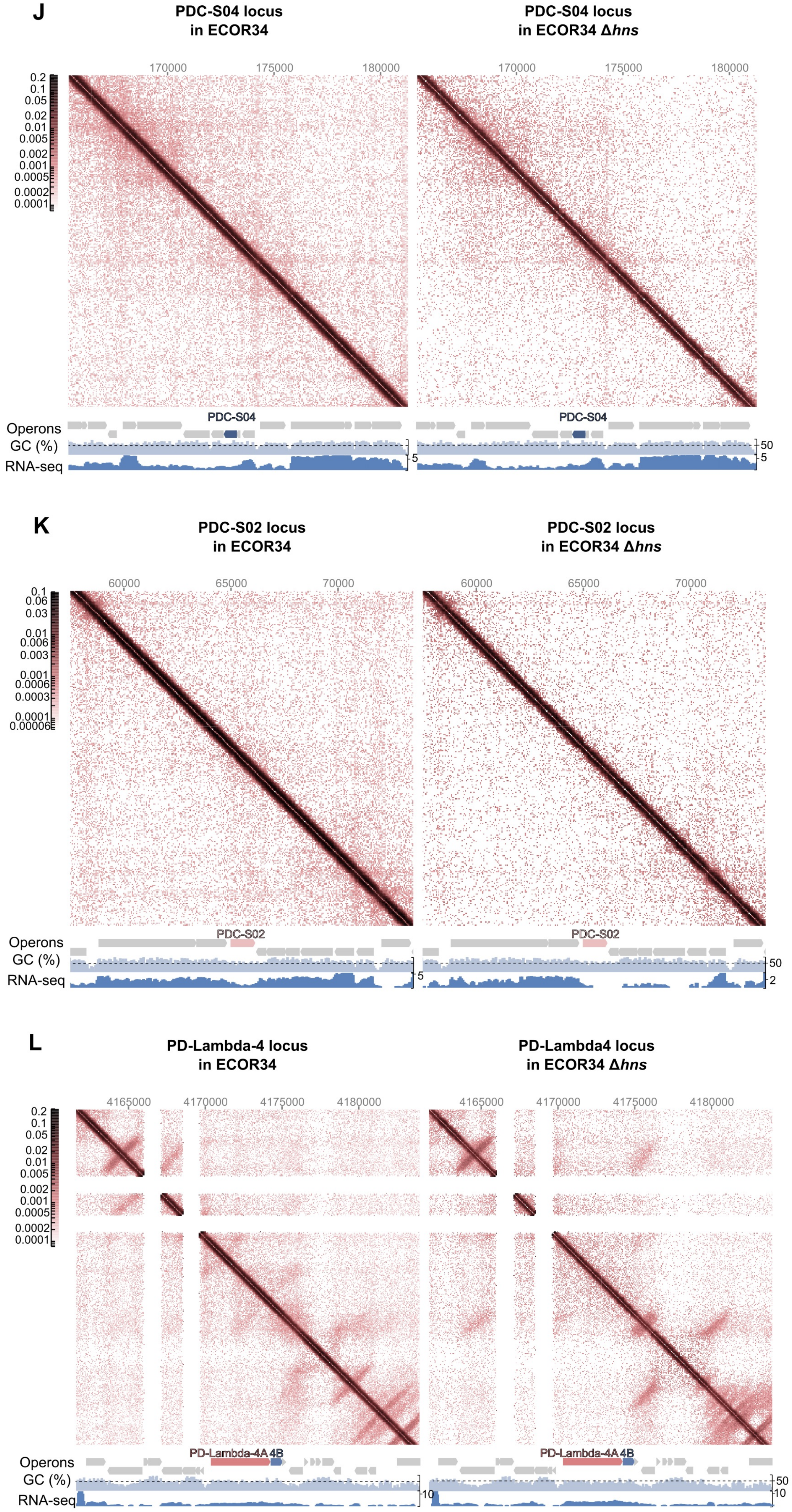
J
PDC-S04 locus in ECOR34
PDC-S04 locus in ECOR34 Δ*hns*
170000
175000
180000
0.2
0.1
0.05
0.02
0.01
0.005
0.002
0.001
0.0005
0.0002
0.0001
PDC-S04
Operons
GC (%)
RNA-seq
50
5
K
PDC-S02 locus in ECOR34
PDC-S02 locus in ECOR34 Δ*hns*
60000
65000
70000
0.1
0.06
0.03
0.01
0.006
0.003
0.001
0.0006
0.0003
0.0001
0.00006
PDC-S02
Operons
GC (%)
RNA-seq
50
5
2
L
PD-Lambda-4 locus in ECOR34
PD-Lambda4 locus in ECOR34 Δ*hns*
4165000
4170000
4175000
4180000
0.2
0.1
0.05
0.02
0.01
0.005
0.002
0.001
0.0005
0.0002
0.0001
PD-Lambda-4A 4B
Operons
GC (%)
RNA-seq
50
10

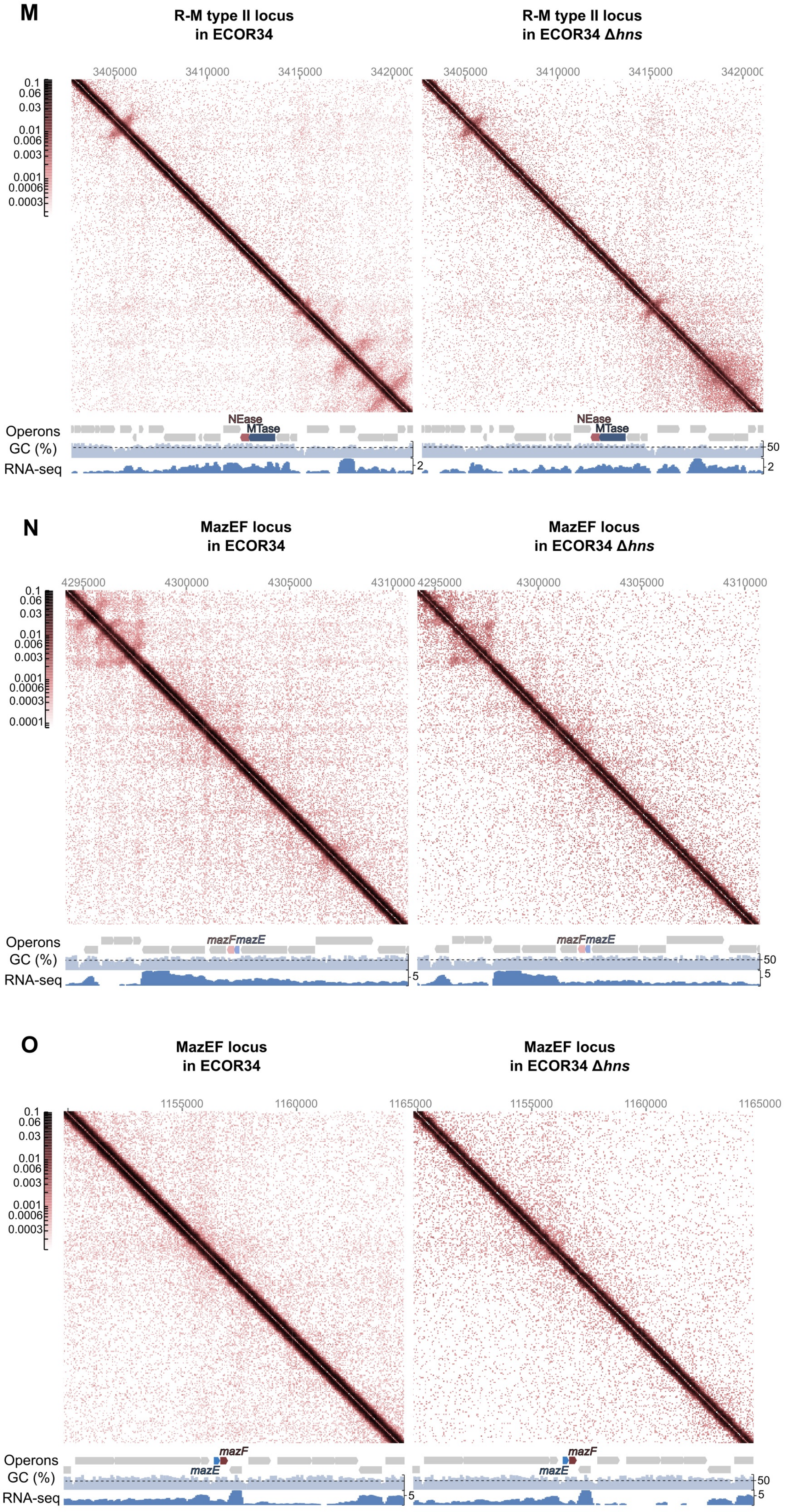
M
R-M type II locus in ECOR34
R-M type II locus in ECOR34 Δhns
3405000
3410000
3415000
3420000
0.1
0.06
0.03
0.01
0.006
0.003
0.001
0.0006
0.0003
NEase
MTase
Operons
GC (%)
RNA-seq
50
2
N
MazEF locus in ECOR34
MazEF locus in ECOR34 Δhns
4295000
4300000
4305000
4310000
0.0001
mazF
mazE
5
O
MazEF locus in ECOR34
MazEF locus in ECOR34 Δhns
1155000
1160000
1165000

P

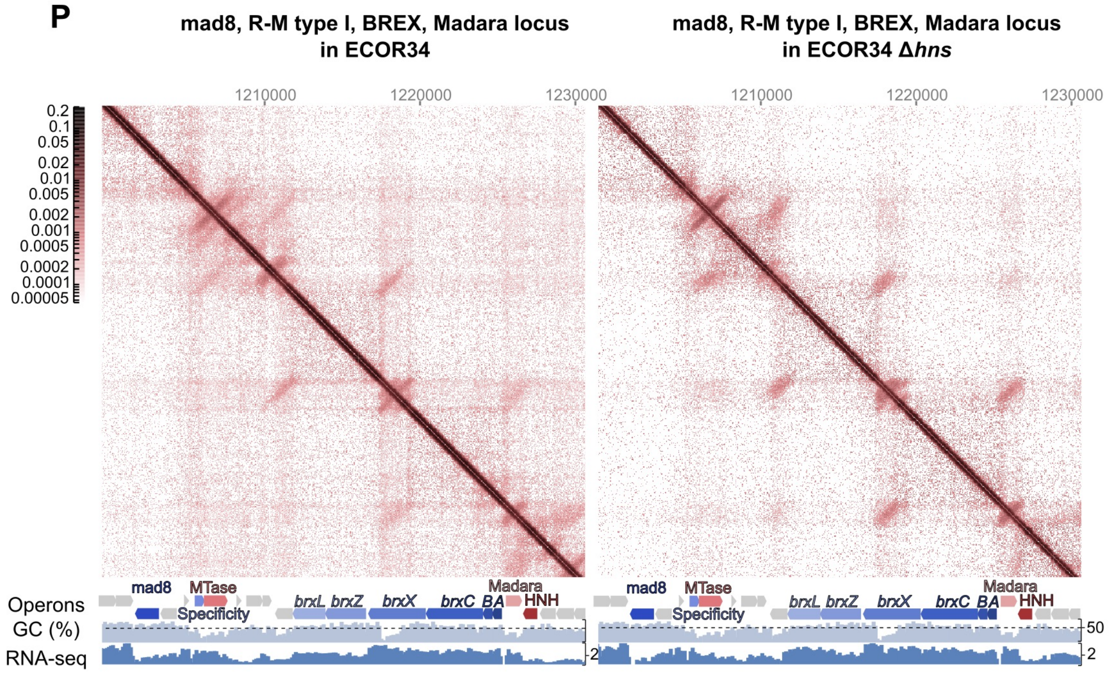
mad8, R-M type I, BREX, Madara locus
in ECOR34
mad8, R-M type I, BREX, Madara locus
in ECOR34 Δhns
1210000
1220000
1230000
0.2
0.1
0.05
0.02
0.01
0.005
0.002
0.001
0.0005
0.0002
0.0001
0.00005
Operons
GC (%)
RNA-seq
mad8
MTase
Specificity
brxL
brxZ
brxX
brxC
BA
Madara
HNH
50
2

Q

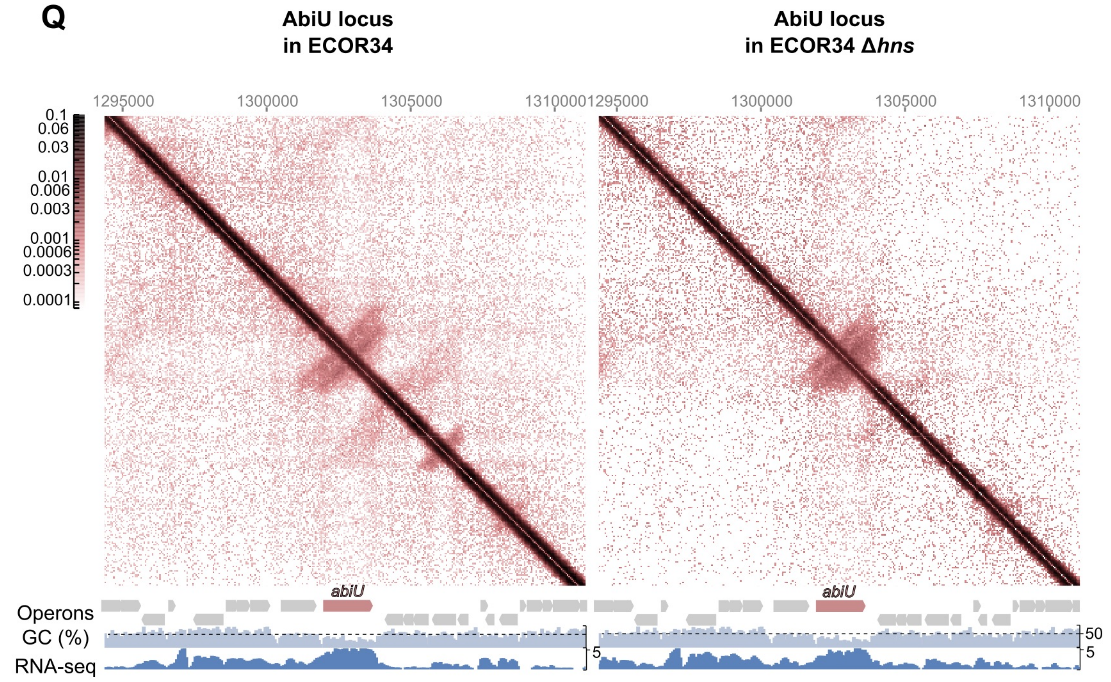
AbiU locus
in ECOR34
AbiU locus
in ECOR34 Δhns
1295000
1300000
1305000
1310000
0.1
0.06
0.03
0.01
0.006
0.003
0.001
0.0006
0.0003
0.0001
Operons
GC (%)
RNA-seq
abiU
50
5
