## Supplementary data for "H-NS silences antiviral immunity through 3D chromatin compaction"

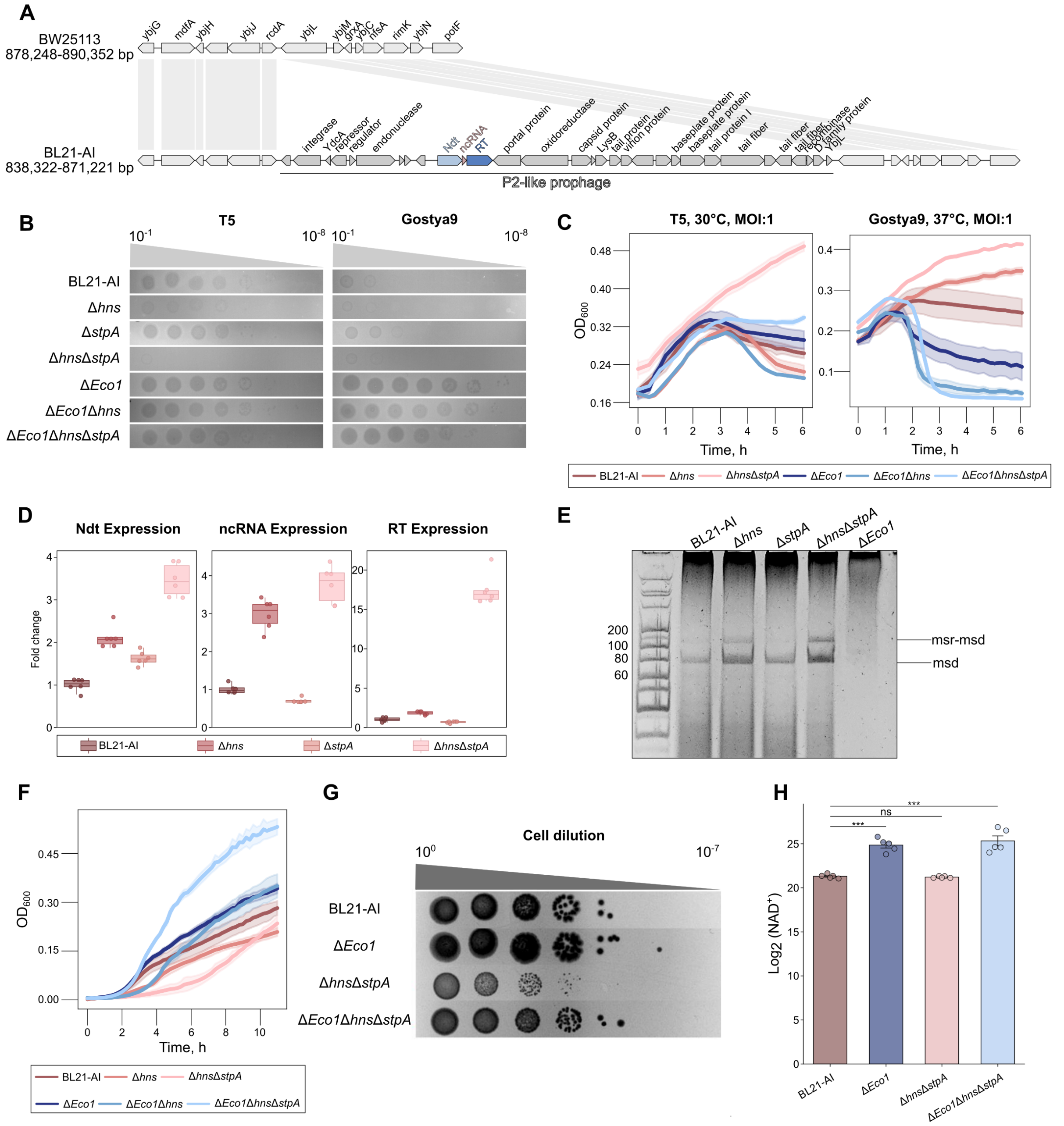


**Figure S1.** H-NS deletion activates Eco1 retron system in *E. coli* BL21-AI and causes Eco1-dependent toxicity.

**(A)** Alignment of the P2-like prophage loci between *E. coli* BL21-AI and prophage-free BW25113. **(B)** H-NS and the double H-NS/StpA deletions in BL21-AI enhance defense against T5 and Gostya9 phages in EOP assay (37°C). **(C)** Liquid culture infections of *E. coli* BL21-AI with phages T5 and Gostya9 demonstrate Eco1-dependent defense upon H-NS and the double H-NS/StpA deletions. **(D)** Expression levels of the Eco1 system components in BL21-AI and its derivatives, as determined by RT–qPCR. Signal is normalized to 16S rRNA expression level, with the expression level in WT cells set to 1 (37°C). **(E)** PAGE analysis of retron msDNA production (30°C) in *E. coli* BL21-AI and its derivatives (15% denaturing polyacrylamide gel) **(F)** Liquid culture growth curves of *E. coli* BL21-AI and indicated derivatives (30°C). **(G)** Spot-dilution toxicity assay of *E. coli* BL21-AI and indicated derivatives (30°C). **(H)** LC-MS $NAD^+$ concentration analysis in *E. coli* BL21-AI and indicated derivatives (30°C).

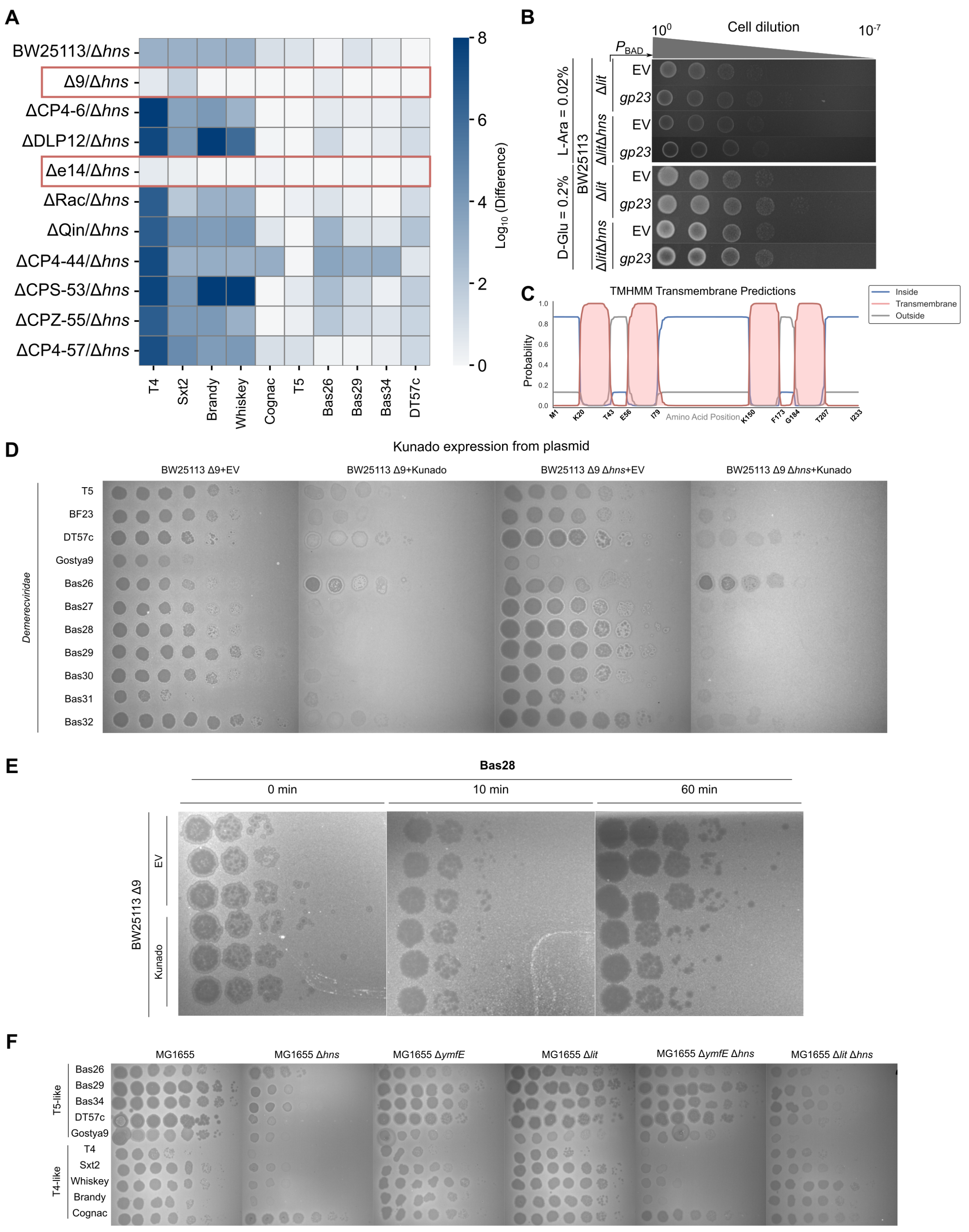


**Figure S2.** Supporting data for the Figure 2.

**(A)** Heatmap of the EOP assay demonstrating the difference in the phage titer for BW25113 strains with the deletion of each of the 9 prophages and its *Δhns* derivatives. **(B)** Toxicity assay BW25113 / *Δhns* / *Δlit* / *Δlit Δhns* + E.V / pBAD *gp23* on LB agar plates (+L-Ara 0.02% / D-Glu 0.02%). **(C)** TMHMM prediction of Kunado (YmfE) transmembrane regions. **(D)** EOP assay demonstrating the defense activity of Kunado (YmfE) with native promoter from pBR322 (37°C). **(E)** Titer of the free Bas26 phage particles in the presence of the Kunado system, measured at 0, 10, and 60 min after the phage addition. **(F)** EOP assay demonstrating the defense activity of native Kunado (YmfE) and Lit in the MG1655 strain (37°C).

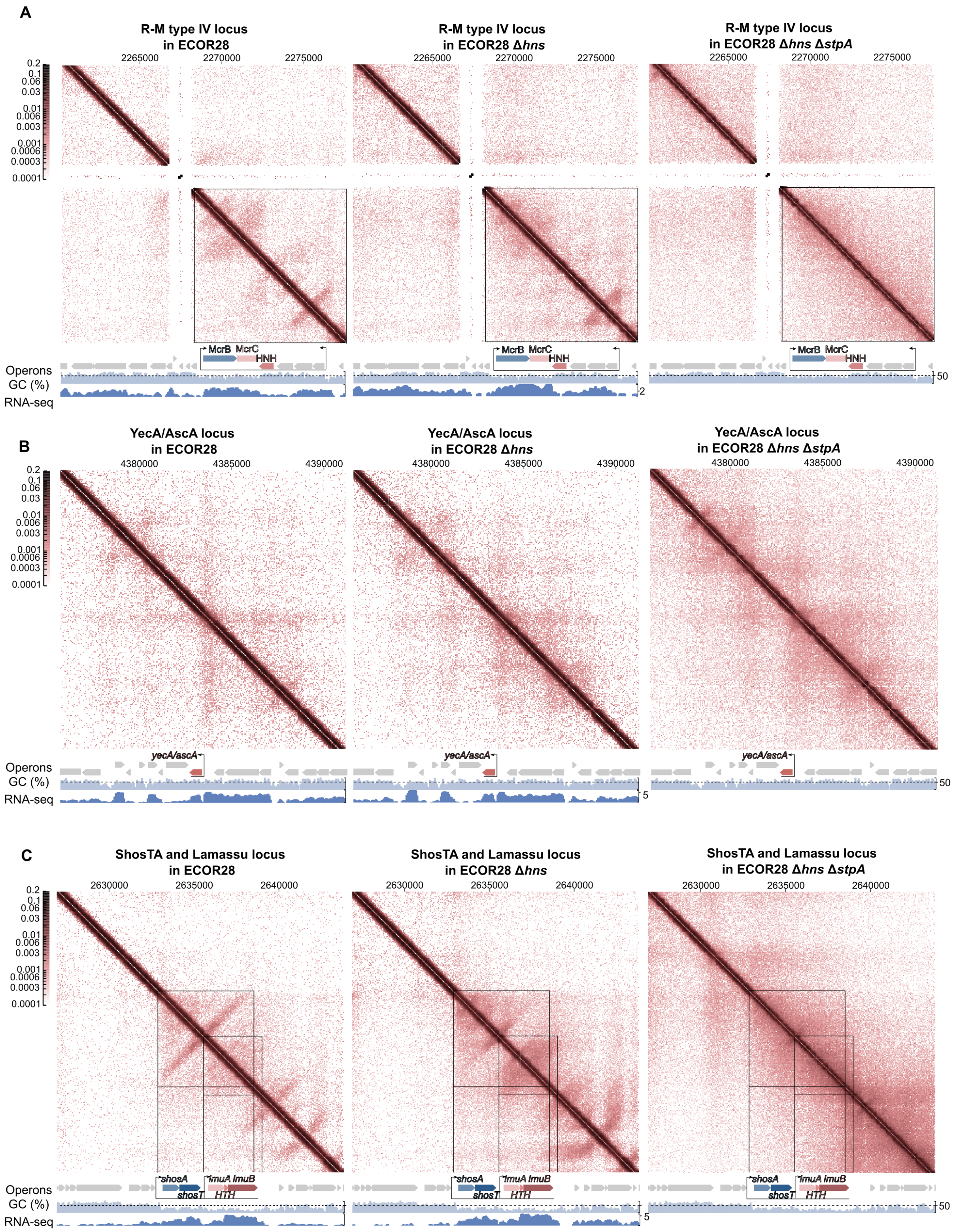

A
R-M type IV locus in ECOR28
R-M type IV locus in ECOR28 Δhns
R-M type IV locus in ECOR28 Δhns ΔstpA
2265000
2270000
2275000
McrB
McrC
HNH
Operons
GC (%)
RNA-seq
B
YecA/AscA locus in ECOR28
YecA/AscA locus in ECOR28 Δhns
YecA/AscA locus in ECOR28 Δhns ΔstpA
4380000
4385000
4390000
yecA/ascA
C
ShosTA and Lamassu locus in ECOR28
ShosTA and Lamassu locus in ECOR28 Δhns
ShosTA and Lamassu locus in ECOR28 Δhns ΔstpA
2630000
2635000
2640000
shosA
shosT
lmuA
lmuB
HTH

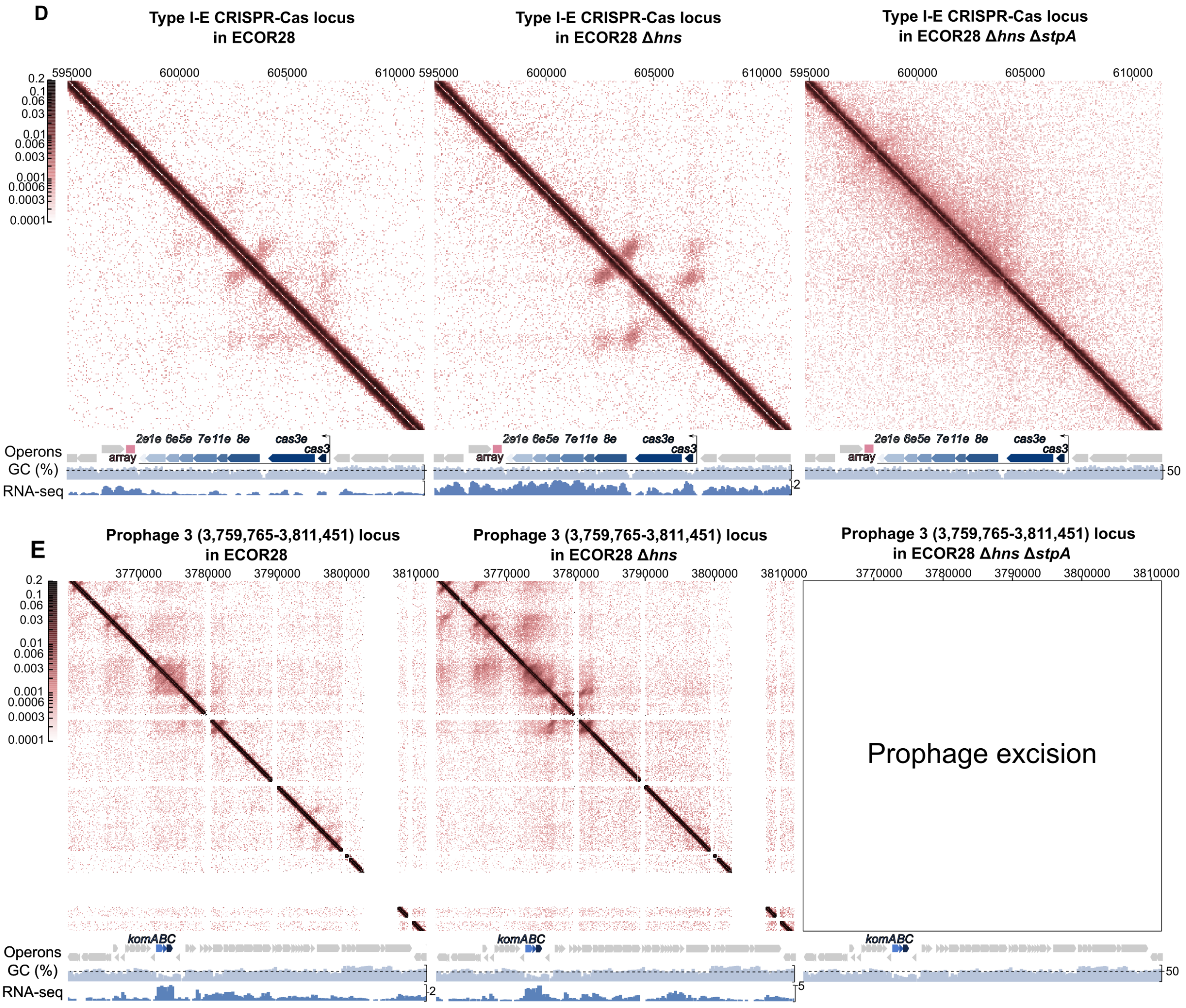


**Figure S3.** 3D chromosome compactization, GC content and gene expression levels ($\log_2$(CPM+1)) of the defense loci in the ECOR28, *Δhns*, and *Δhns/ΔstpA* derivatives: **(A)** R-M IV (*mrcBC*) locus. **(B)** PDC-S21(*yecA/ascA*) locus. **(C)** ShosTA and Lamassu loci. **(D)** CRISPR-Cas I-E locus. **(E)** Prophage 3 loci with Kongming.

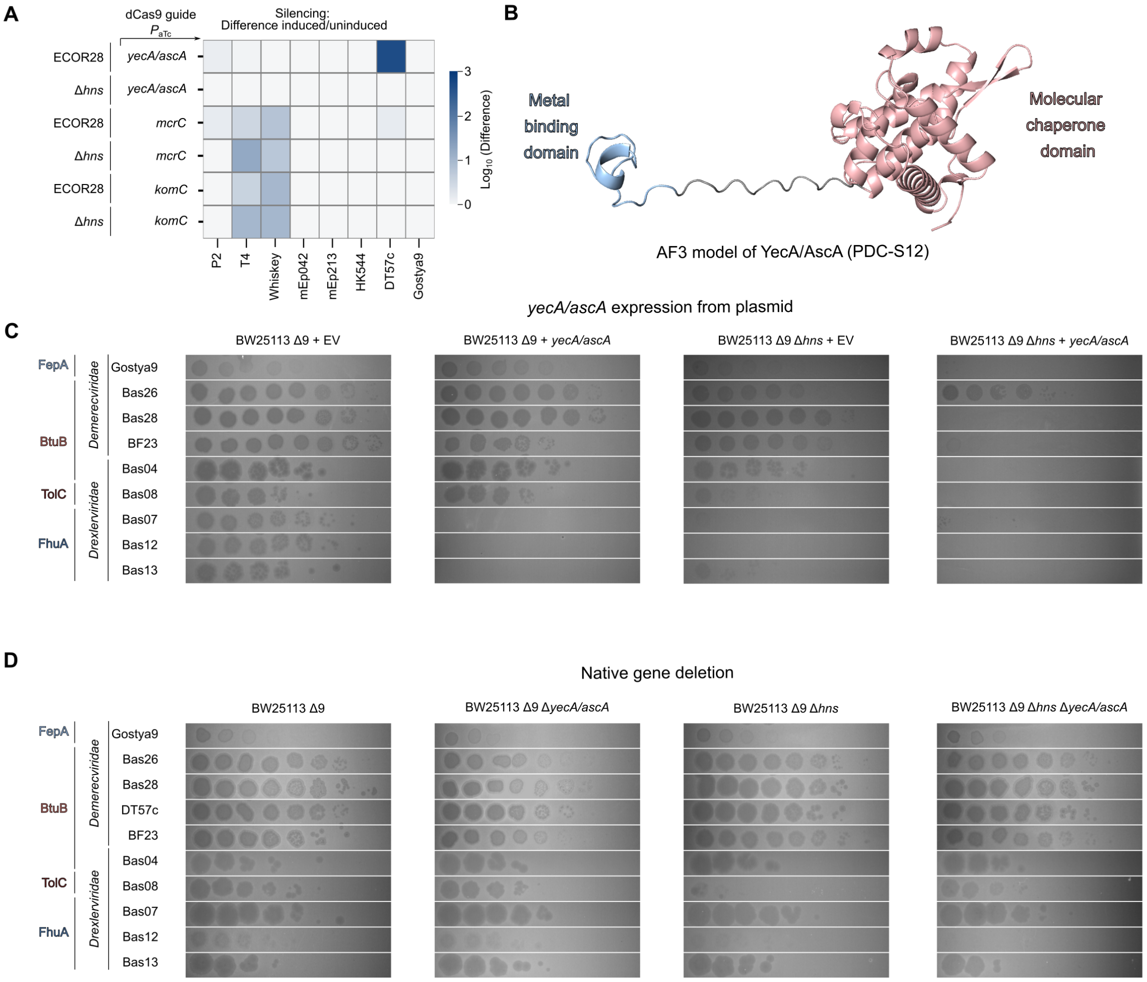


**Figure S4.** Supporting data for the Figure 3.

**(A)** Heatmap of the phage titers measured in conditions of ECOR28 dCas9 silencing. **(B)** AF3 model of the PDC-S12 (YecA/AscA) protein monomer. **(C)** EOP assay demonstrating the defense activity of PDC-S12 (*yecA/ascA*) expressed from a plasmid under its native promoter. **(D)** EOP assay demonstrating the defense activity of the native PDC-S12 (*yecA/ascA*) gene in *E.coli* BW25113.

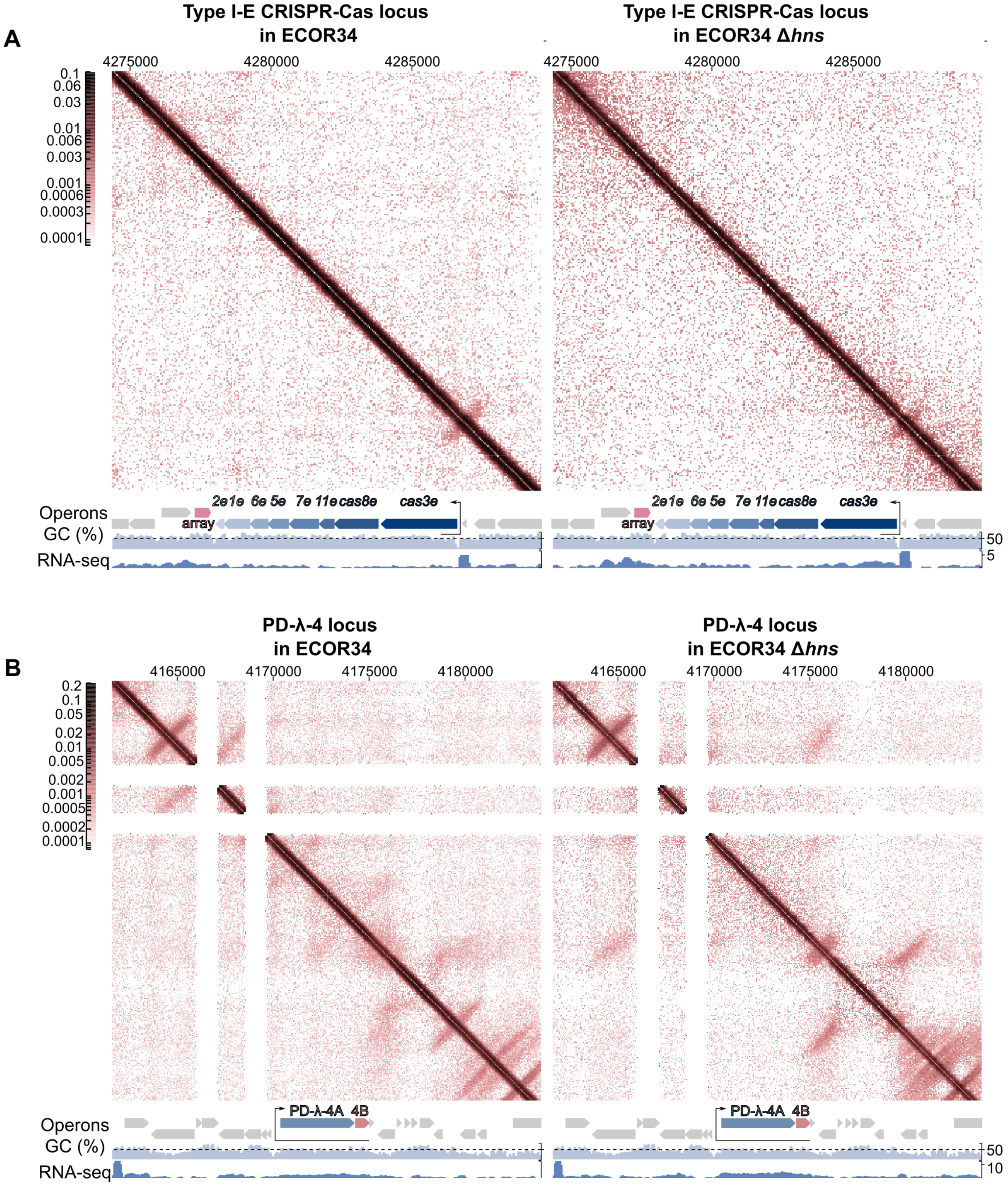


**Figure S5.** 3D chromosome compactization, GC content and gene expression levels ($\log_2$(CPM+1)) of the defense loci in the ECOR234, and *Δhns* derivative: **(A)** CRISPR-Cas I-E locus. **(B)** PD-λ-4 locus.

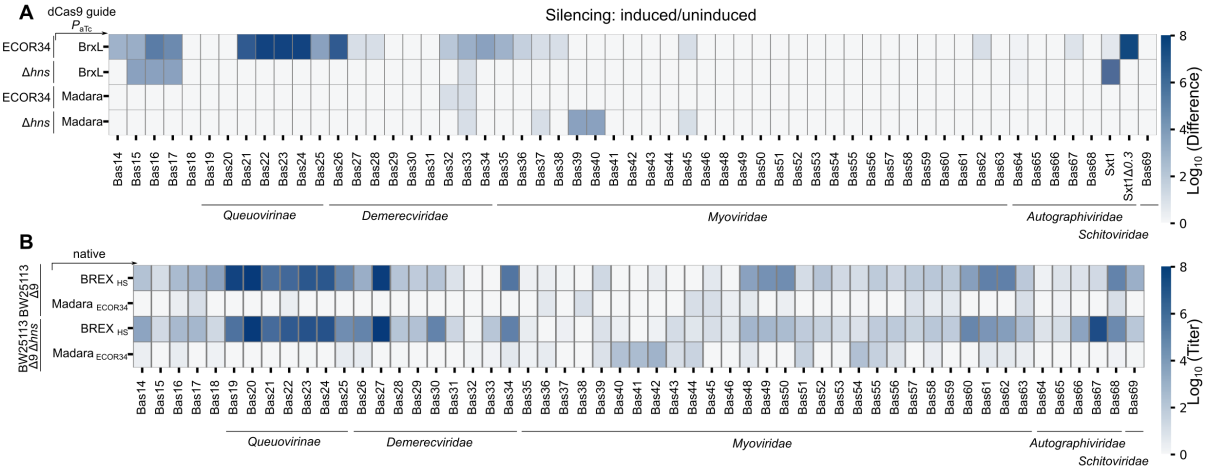


**Figure S6.** Supporting data for the Figure 4.

**(A)** Heatmap of the phage titers measured in conditions of ECOR34 dCas9 silencing. **(B)** Heatmap of the EOP assay performed with the BASEL phage collection against BW25113 in conditions of the BREX HS or Madara ECOR34 expression from the plasmid.

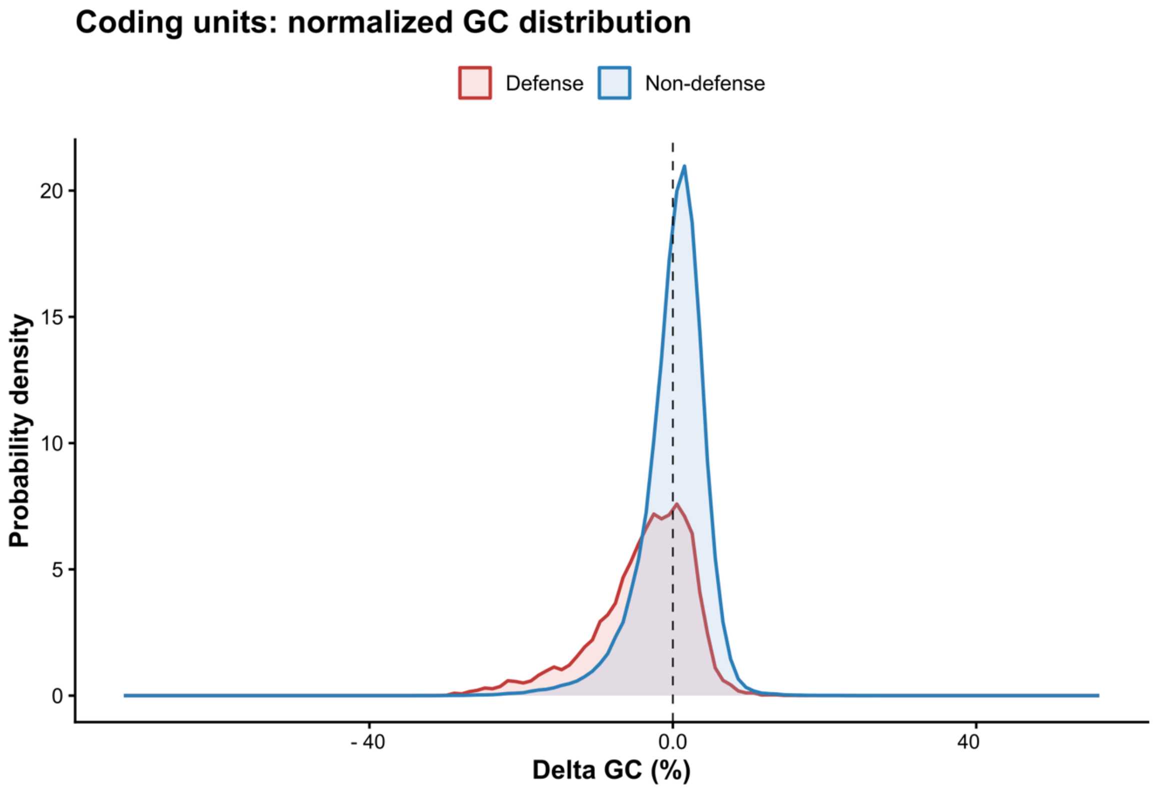


**Figure S7.** Phage defense genes have lower GC content across bacterial genomes.

Probability density of the distribution of GC-content of defense and non-defense genes across complete bacterial genomes. Delta GC (%) is the difference in GC content between a given gene and its host genome, calculated for each gene individually.

**Supplementary table 1.** List of *E.coli* strains used in this study.

| *E. coli* **strain** | **Description** | **Source** |
|---|---|---|
| BL21-AI | F-*ompT hsdSB (rB- mB-) gal dcm araB::T7RNAP-tetA* | Invitrogen |
| BL21-AI Δ*Eco1* | BL21-AI Δ*Eco1::FRT* | [1] |
| BL21-AI Δ*hns* | Bl21-AI Δ*hns::FRT* | This study |
| BL21-AI Δ*stpA* | Bl21-AI Δ*stpA::FRT* | This study |
| BL21-AI Δ*Eco1*Δ*hns* | BL21-AI Δ*Eco1::FRT* Δ*hns::FRT* | This study |
| BL21-AI Δ*Eco1*Δ*hns*Δ*stpA* | BL21-AI Δ*Eco1::FRT* Δ*hns::FRT* Δ*stpA::FRT* | This study |
| XL1-Blue | *E. coli K12 recA1 endA1 gyrA96 thi-1 hsdR17 supE44 relA1 lac [F'proAB lacIqZΔM15 Tn10], TetR* | Evrogen |
| BW25113 | *E. coli* K12 F– Δ(*araD-araB*)567 Δ*lacZ*4787(::*rrnB*-3) λ– *rph*-1 Δ(*rhaD-rhaB*)568 *hsdR*514 | Lab stock |
| BW25113 Δ*hns* | BW25113 Δ*hns::FRT* | KEIO collection |
| BW25113 Δ*stpA* | BW25113 Δ*hns::FRT* | KEIO collection |
| BW25113 Δ*hns*Δ*stpA* | BW25113 Δ*hns::FRT* Δ*stpA::FRT* | This study |
| BW25113 Δ*lit* | BW25113 Δ*lit::FRT* | KEIO collection |
| BW25113 Δ*mcrA* | BW25113 Δ*mcrA::FRT* | KEIO collection |
| BW25113 Δ*hns*Δ*lit* | BW25113 Δ*hns::FRT* Δ*lit::FRT* | This study |
| BW25113 Δ*hns*Δ*mcrA* | BW25113 Δ*hns::FRT* Δ*mcrA::FRT* | This study |

| | | |
|---|---|---|
| BW25113 Δ*9* | Deletion of 9 cryptic prophages | [2] |
| BW25113 Δ*Qin* | BW25113 Δ*Qin::FRT* | [2] |
| BW25113 Δ*e14* | BW25113 Δ*e14::FRT* | [2] |
| BW25113 Δ*LP12* | BW25113 Δ*LP12::FRT* | [2] |
| BW25113 Δ*CP4-6* | BW25113 Δ*CP4-6::FRT* | [2] |
| BW25113 Δ*CPz-55* | BW25113 Δ*CPz-55::FRT* | [2] |
| BW25113 Δ*CPS-53* | BW25113 Δ*CPS-53::FRT* | [2] |
| BW25113 Δ*rac* | BW25113 Δ*rac::FRT* | [2] |
| BW25113 Δ*CP4-44* | BW25113 Δ*CP4-44::FRT* | [2] |
| BW25113 Δ*CP4-57* | BW25113 Δ*CP4-57::FRT* | [2] |
| BW25113 Δ*9*Δ*hns* | BW25113 Δ*9* Δ*hns::FRT* | This study |
| BW25113 Δ*Qin*Δ*hns* | Δ*Qin::FRT* Δ*hns::FRT* | This study |
| BW25113 Δ*e14*Δ*hns* | Δ*e14::FRT* Δ*hns::FRT* | This study |
| BW25113 Δ*LP12*Δ*hns* | Δ*LP12::FRT* Δ*hns::FRT* | This study |
| BW25113 Δ*CP4-6*Δ*hns* | Δ*CP4-6::FRT* Δ*hns::FRT* | This study |
| BW25113 Δ*CPz-55*Δ*hns* | Δ*CPz-55::FRT* Δ*hns::FRT* | This study |
| BW25113 Δ*CPS-53*Δ*hns* | Δ*CPS-53::FRT* Δ*hns::FRT* | This study |
| BW25113 Δ*rac*Δ*hns* | Δ*rac::FRT* Δ*hns::FRT* | This study |
| BW25113 Δ*CP4-44*Δ*hns* | Δ*CP4-44::FRT* Δ*hns::FRT* | This study |

| | | |
|---|---|---|
| BW25113 Δ*CP4-57*Δ*hns* | Δ*CP4-57::FRT* Δ*hns::FRT* | This study |
| BW25113 Δ*9*Δ*hns*Δ*stpA* | BW25113 Δ*9* Δ*hns::FRT* Δ*stpA::FRT* | This study |
| BW25113 Δ*9*Δ*stpA* | BW25113 Δ*9* Δ*stpA::FRT* | This study |
| MG1655 Δ*stpA* | MG1655 Δ*stpA::FRT* | This study |
| MG1655 Δ*hns*Δ*stpA* | MG1655 Δ*hns::FRT* Δ*stpA::FRT* | This study |
| MG1655 Δ*lit* | MG1655 Δ*lit::FRT* | This study |
| MG1655 Δ*hns*Δ*lit* | MG1655 Δ*hns::FRT* Δ*lit::FRT* | This study |
| MG1655 Δ*stpA*Δ*lit* | MG1655 Δ*stpA::FRT* Δ*lit::FRT* | This study |
| MG1655 Δ*hns*Δ*stpA*Δ*lit* | MG1655 Δ*hns::FRT* Δ*stpA::FRT* Δ*lit::FRT* | This study |
| MG1655 Δ*ymfE* | MG1655 Δ*ymfE::FRT* | This study |
| MG1655 Δ*hns*Δ*ymfE* | MG1655 Δ*hns::FRT* Δ*ymfE::FRT* | This study |
| ECOR28 | | Lab stock |
| ECOR28 Δ*hns* | ECOR28 Δ*hns::FRT* | This study |
| ECOR28 Δ*stpA* | ECOR28 Δ*stpA::FRT* | This study |
| ECOR28 Δ*hns*Δ*stpA* | ECOR28 Δ*hns::FRT* Δ*stpA::FRT* | This study |
| ECOR34 | | Lab stock |
| ECOR34 Δ*hns* | ECOR34 Δ*hns::FRT* | This study |
| ECOR34 Δ*stpA* | ECOR34 Δ*stpA::FRT* | This study |

**Supplementary table 2.** List of phages used in this study.

| Phage | Comment | Source |
|---|---|---|
| T5 | | Lab stock |
| T5Mos | Deletions of 8 kb fragment | [3] |
| Gostya9 | | [4] |
| BF23 | | [5] |
| Bas01-Bas69 | | [6] |
| DT57c | | Lab stock |
| Msk8 | | Lab stock |
| Msk10 | | Lab stock |
| Msk11 | | Lab stock |
| Msk12 | | Lab stock |
| Msk14 | | Lab stock |
| P21a | | Lab stock |
| LF82 | | Lab stock |
| 9g | | Lab stock |
| 3.2 | | Lab stock |
| Quenovirus | dPreQ0 mutated deoxyguanosine | Lab stock |
| HK75 | | Lab stock |

| | | |
|---|---|---|
| HK97 | | Lab stock |
| HK106 | | Lab stock |
| HK140 | | Lab stock |
| HK225 | | Lab stock |
| HK446 | | Lab stock |
| HK544 | | Lab stock |
| mEp042 | | Lab stock |
| mEp213 | | Lab stock |
| HK573 | | Lab stock |
| T4 | | Lab stock |
| T4 gp2 | | Lab stock |
| T6 | | Lab stock |
| Sxt2 (T4) | Obtained from Sextaphage mix (Microgene) | Lab stock |
| RB49 | | [7] |
| Brandy | | [7] |
| Whiskey | | [7] |
| Cognac | | [7] |
| Felix01 | | [7] |
| P1 | | Lab stock |

| | | |
|---|---|---|
| P2 | | Lab stock |
| T7 | | Lab stock |
| T7Δ*0.3* | | Lab stock |
| T7Δ*5.9* | | Lab stock |
| Sxt1 | Obtained from Sextaphage mix (Microgene) | [8] |
| Sxt1Δ*0.3* | | Lab stock |
| λ Vir | | Lab stock |
| λ 872 | | Lab stock |

**Supplementary table 3.** List of plasmids used in this study.

| Plasmid Name | Plasmid Map | Comment | Source |
|---|---|---|---|
| pBAD | https://benchling.com/s/seq-FZKEYBxvFNpLmZCe9L4x?m=slm-wx17YqwMMNNPiXKliKTX | *araBAD* promoter, $Amp^R$ | Lab stock |
| pBAD gp23 T4 | https://benchling.com/s/seq-aVM1lAmTrEjYYRGKC2JE?m=slm-R2WMyI6bERuc7o4CKFCQ | Gp23 T4 under araBAD promoter, $Amp^R$ | This study |
| pBR322 | https://benchling.com/s/seq-rRdpvoDDeuxwv3nX2a78?item_id=seq_tUf4vEM2ND&m=slm-uxWc4ONSOqFim2CLgzRu | $Amp^R$ deletion, $Tet^R$ | Lab stock |

| | | | |
|---|---|---|---|
| pBR322 Madara | https://benchling.com/s/seq-7bq25vBDLZedgDz3XDrh?m=slm-JDBbeoUf2XLUXF4ot0GQ | Madara defense system under native promoter, $Tet^R$ | This study |
| pBR322 Madara D63A | https://benchling.com/s/seq-yPYFhjjoDCroC0WqDOSt?item_id=seq_a1IqpBwCrn&m=slm-9to0iEI62RzQhXMEPwAU | Madara defense system under native promoter, D63A point mutation, $Tet^R$ | This study |
| pBR322 Kunado | https://benchling.com/s/seq-7Zm80Bk1aO0kl07fo3es?m=slm-4e0w02zRjVaE2ieZF0xR | ymfE gene under native promoter, $Tet^R$ | This study |
| pBR322 yecA/ascA | https://benchling.com/s/seq-Nc3erdwD0H1i8KEz3wuG?item_id=seq_WnIaWYoNuH&m=slm-RHEzxZgD6yh6j1c0thM3 | yecA/ascA gene under native promoter, $Tet^R$ | This study |
| pFD152 dCas9 | https://benchling.com/s/seq-2WyXAwSfEnMOZ1g1ctVr?m=slm-f4EUUVhtwAMS5IemCHuh | $Cm^R$ | Lab stock |
| pFD152 brxL | https://benchling.com/s/seq-uS1ZVmVr7PhB1yFuPzez?m=slm-ZfPzsy5idiWcLtHIlYtp | brxL spacer, $Cm^R$ | This study |
| pFD152 yhfG | https://benchling.com/s/seq-ymPgiDebMQlSksNmeYIi?m=slm-6cwIPwuAfGo6UZDTVpzX | yhfG spacer, $Cm^R$ | This study |
| pFD152 shosT | https://benchling.com/s/seq-geOoXI4njdSTUNWoOOFN?m=slm- | shosT spacer, $Cm^R$ | This study |

| | | | |
|---|---|---|---|
| | tlpSYAZHwEyD6xRK8meh | | |
| pFD152 pdc-s04 | https://benchling.com/s/seq-BeVxWNvx4axrRxOlryEs?m=slm-vMvvH0wfTyLv0SyuxFWk | pdc-s04 spacer, $Cm^R$ | This study |
| pFD152 pdc-s07 | https://benchling.com/s/seq-mLh5IWphLxm0JpJfmLi3?m=slm-wnyPyHuu66EbkRRTqUbA | pdc-s07 spacer, $Cm^R$ | This study |
| pFD152 pdc-s11 | https://benchling.com/s/seq-x2x0trKSTpxp3xHEN45u?m=slm-QdkMtow102qrwrFlCzNC | pdc-s11 spacer, $Cm^R$ | This study |
| pFD152 pdc-s12 | https://benchling.com/s/seq-oJdPmgat6c9Rq9JL5xIk?m=slm-em3PQ4g0TlWoxbxS2Kgl | pdc-s12 spacer, $Cm^R$ | This study |
| pFD152 pd-lambda-1 | https://benchling.com/s/seq-63THbnyAOeM0gujEGxQE?m=slm-WynnEE0UkEsmuH2xx92w | pd-lambda-1 spacer, $Cm^R$ | This study |
| pFD152 pd-lambda-4a | https://benchling.com/s/seq-7X8ldmptbk1An23TdDPQ?m=slm-UgMA2gPWpCLBmLfFoHb0 | pd-lambda-4a spacer, $Cm^R$ | This study |
| pFD152 RM IV | https://benchling.com/s/seq-b5Z4TzK79Fe4tnMEMXoT?m=slm-Au2PlzTATwu7ydtBKJby | RM IV spacer, $Cm^R$ | This study |
| pFD152 lmuB | https://benchling.com/s/seq-avzwdQ8N9IKeYRNoBDfC?m=slm- | lmuB spacer, $Cm^R$ | This study |

| | | | |
|---|---|---|---|
| | kqNmAICHwbYhH55AwGzA | | |
| pFD152 komC | https://benchling.com/s/seq-qr5t10khx49KAaNyTzEU?item_id=seq_gBu7rYQKbi&m=slm-AB6acZ3ktaJ7GUfdk93G | komC spacer, $Cm^R$ | This study |

**Supplementary table 4.** List of oligos used in this study.

| **Oligo name** | **Sequence (5' to 3')** | **Length, nt** |
|---|---|---|
| qRT2_F | ggtgctccatcatcacctaaa | 21 |
| qRT2_R | tgactacgaggcccactaata | 21 |
| qEff_F | accgatgagcagcaacaa | 18 |
| qEff_R | cgacagcttcagccagaata | 20 |
| qE1 | gtcagaaaaaacgggtttcctggttgg | 27 |
| qE2 | tctgagttactgtctgttttccttgttgg | 29 |
| qE3 | cctctggatgttgtttcggca | 21 |
| rpoC_F | cgtactttacgccatgaacgtactaggc | 28 |
| rpoC_R | gccatgcggacgatcgtgtcatag | 24 |
| pBAD_lin_F | taaattattcccctagcg | 19 |

| | | |
|---|---|---|
| pBAD_lin_R | ggttaattcctcctgttag | 19 |
| pBAD check_F | atgccatagcatttttatcc | 20 |
| pBAD check_R | gatttaatctgtatcaggctg | 21 |
| pBAD Gp23 T4 F | cgagctctaaggaggttataaaaaatgactatcaaaactaaagctg | 47 |
| pBAD Gp23 T4 R | caagcttgcatgcctgcaggtcgacttagatacctttaacatatacacg | 49 |
| pBR322 Madara F | gtcatgagattatcaaaaaggatcttttattgttcagctttttcaac | 47 |
| pBR322 Madara R | atacattcaaatatgtatccgctcattgtgagagggatgatcg | 43 |
| pBR322 Madara D63A F | tcgccgacgactatcttc | 18 |
| pBR322 Madara D63A R | attgcgttattggcaatcgataaa | 24 |
| pFD152 brxL F | tagttgaaacaggtaaatcatgat | 24 |
| pFD152 brxL R | aaacatcatgatttacctgtttca | 24 |
| pFD152 yhfG F | tagtacgctgaagctcccagagac | 24 |
| pFD152 yhfG R | aaacgtctctgggagcttcagcgt | 24 |
| pFD152 Pdcs11 F | tagtctgtaatccttccagagtaa | 24 |
| pFD152 Pdcs11 R | aaacttactctggaaggattacag | 24 |
| pFD152 Pdcs04 F | tagttcaaggccaggataaagata | 24 |
| pFD152 Pdcs04 R | aaactatctttatcctggccttga | 24 |

| | | |
|---|---|---|
| pFD152 Pdcs07 F | tagttcacgcaccaccgcctgttc | 24 |
| pFD152 Pdcs07 R | aaacgaacaggcggtggtgcgtga | 24 |
| pFD152 Pdcs12 F | tagttccaactcactttcgtttaa | 24 |
| pFD152 Pdcs12 R | aaacttaaacgaaagtgagttgga | 24 |
| pFD152 Pdlambda4a F | tagtcagctcttccagctggatat | 24 |
| pFD152 Pdlambda4a R | aaacatatccagctggaagagctg | 24 |
| pFD152 ShosT F | tagttcgctatcgatctttccctt | 24 |
| pFD152 ShosT R | aaacaagggaaagatcgatagcga | 24 |
| pFD152 LmuB F | tagtccccatctttattggctatc | 24 |
| pFD152 LmuB R | aaacgatagccaataaagatgggg | 24 |
| pFD152 RM IV F | tagtttcgaacacagtgaaatgac | 24 |
| pFD152 RM IV R | aaacgtcatttcactgtgttcgaa | 24 |
| pFD152 komC F | tagttcaaccaattgactccatga | 24 |
| pFD152 komC R | aaactcatggagtcaattggttga | 24 |
| Lit check F | ttaatcgttcccagtgtgcc | 20 |
| Lit check R | agcaagaattgactggggag | 20 |
| mcrA check F | aggtttcctgcaggggataa | 20 |

| | | |
|---|---|---|
| mcrA check R | taacaatcaggtcagccgct | 20 |
| ymfE check F | ttgctctctttcaagttgcc | 20 |
| ymfE check R | ctggacactctgtcatcatt | 20 |
| hns check F | tgtcttaaaccggacaataaaaaatc | 26 |
| hns check R | ctggctattgcacaactgaatttaag | 26 |
| stpA check F | actgtttgcaggaatcagcg | 20 |
| stpA check R | ctgaaataatctcgcgcagg | 20 |
| yecA check F | tgacctggcaaagcagaaag | 20 |
| yecA check R | gtggaatcggtagacacaag | 20 |

**Supplementary table 5.** List of predicted BL21-AI defense systems and prophages.

| BL21-AI | |
|---|---|
| **Defense system name** | **Location, nucleotide positions** |
| PDC-S07 (mutT) | 116295-116684 |
| Retron Eco1 | 849501-851548 |
| PDC-S07 (nudG) | 1757477-1757884 |
| PDC-S12 (yecA) | 1907707-1908372 |
| MazEF | 2714956-2715539 |
| PDC-S04 | 3322073-3322675 |
| PDC-S13 | 3656162- 3656986 |
| R-M type IV | 4454833- 4457258 |
| R-M type I | 4459470-4465485 |
| **Prophage** | **Location, nucleotide positions** |
| Prophage 1 | 535387-553713 |
| Prophage 2 | 746487-756988 |
| P2-like prophage | 844110-866781 |
| Prophage 4 | 1368779-1381394 |
| Prophage 4 | 1548424-1572481 |

**Supplementary table 6.** List of predicted MG1655 defense systems and prophages.

| MG1655 | |
|---|---|
| **Defense system name** | **Location, nucleotide positions** |
| PDC-S07 (mutT) | 111043-111433 |
| Kunado (yfmE) | 1197333-1198237 |
| Lit | 1198495-1199588 |
| R-M type IV | 1841489-1841897 |
| PDC-S07 (nudG) | 1201633-1203135 |
| PDC-S12 | 1990954-1991619 |

| | |
|---|---|
| VSPR | 2030448-2032317 |
| Hachiman | 2761351-2765153 |
| RnlAB | 2765918-2767355 |
| CRISPR-Cas I-E | 2878569-2887219 |
| MazEF | 2910756-2911339 |
| PDC-S02 | 3367827-3368954 |
| PDC-S04 | 3490861-3491463 |
| PDC-S13 | 3817760-3818584 |
| MazEF/ChpSB | 4448447-4449042 |
| R-M type IV | 4576912-4579337 |
| R-M type I | 4580067-4586761 |
| R-M type IV | 4586948-4587863 |
| **Prophage** | **Location, nucleotide positions** |
| CP4-6 | 262898-297205 |
| DLP12 | 564755-586056 |
| e14 | 1196209-1211412 |
| Rac | 1411899-1434958 |
| Qin | 1632277-1652745 |
| CP4-44 | 2066020-2078748 |
| CPS-53 | 2466369-2476583 |
| CPZ-55 | 2558691-2565480 |
| CP4-57 | 2755941-2777971 |

**Supplementary table 7.** List of predicted BW25113 defense systems and prophages.

| **BW25113** | |
|---|---|
| **Defense system name** | **Location, nucleotide positions** |
| PDC-S02 | 64970-66097 |
| PDC-S04 | 172626-173228 |
| PDC-S13 | 507288-508112 |
| MazEF (1) | 1161371-1161622 |

| | |
|---|---|
| Mad8 | 1201633-1203135 |
| R-M I | 1205470-1207546 |
| BREX | 1211771-1225208 |
| PDC-S11 (Madara) | 1225502-1226500 |
| HNH-nuclease | 1226611-1227483 |
| AbiU | 1301950-1303650 |
| PDC-S07 (mutT) | 1485641-1486030 |
| PDC-S07 (nudG) | 3232765-3233172 |
| PDC-S12 (yecA) | 3374292-3374957 |
| R-M II | 3411774-3413643 |
| PD-Lambda-4 | 4170348-4174991 |
| CRISPR-Cas I-E | 4277969-4286530 |
| MazEF (2) | 4301955-4302538 |
| PDC-S28 (Hna) | 4517187-4519661 |
| **Prophage** | **Location, nucleotide positions** |
| CP4-6 | 258669-292976 |
| DLP12 | 560258-581559 |
| e14 | 1191676-1206868 |
| Rac | 1406156-1429215 |
| Qin | 1626543-1647000 |
| CP4-44 | 2060835-2072512 |
| CPS-53 | 2459864-2470078 |
| CPZ-55 | 2552058-2558820 |
| CP4-57 | 2749316-2771345 |

**Supplementary table 8.** List of predicted ECOR28 defense systems and prophages.

| ECOR28 | |
|---|---|
| **Defense system name** | **Location, nucleotide positions** |
| CRISPR-Cas I-E | 598159-606809 |

| | |
|---|---|
| MazEF (1) | 632126-632709 |
| MazEF (2) | 2200151-2200746 |
| R-M type IV | 2268891-2272252 |
| PDC-S07 (mutT) | 2474295-2474684 |
| ShosTA | 2633129-2635349 |
| Lamassu | 2635800-2638714 |
| PD-Lambda-1 | 2914031-2915533 |
| Kongming | 3772523-3774848 |
| PDC-S07 (nudG) | 4232694-4233101 |
| PDC-S12 (yecA) | 4382678-4383343 |
| VSPR | 4420118-4421987 |
| **Prophage** | **Location, nucleotide positions** |
| Prophage 1 | 2624084-2653799 |
| Prophage 2 | 2903392-2950437 |
| Prophage 3 | 3759765-3811451 |
| Prophage 4 | 4009055-4047213 |

**Supplementary table 9.** List of predicted ECOR34 defense systems and prophages.

| **ECOR34** | |
|---|---|
| **Defense system name** | **Location, nucleotide positions** |
| PDC-S02 | 64970-66097 |
| PDC-S04 | 172626-173228 |
| PDC-S13 | 507288-508112 |
| MazEF (1) | 1161371-1161622 |
| Mad8 | 1201633-1203135 |
| R-M type I | 1205470-1207546 |
| BREX type I | 1211771-1225208 |
| PDC-S11 (Madara) | 1225502-1226500 |
| HNH-nuclease | 1226611-1227483 |
| AbiU | 1301950-1303650 |

| | |
|---|---|
| PDC-S07 (mutT) | 1485641-1486030 |
| PDC-S07 (nudG) | 3232765-3233172 |
| PDC-S12 (yecA) | 3374292-3374957 |
| R-M type II | 3411774-3413643 |
| PD-Lambda-4 | 4170348-4174991 |
| CRISPR-Cas I-E | 4277969-4286530 |
| MazEF (2) | 4301955-4302538 |
| PDC-S28 (Hna) | 4517187-4519661 |
| **Prophage** | **Location, nucleotide positions** |
| Prophage 1 | 1300476-1341401 |
| Prophage 2 | 1946,154-1994040 |
| Prophage 3 | 2988991-3045047 |
| Prophage 4 | 3997061-4037532 |
